# Quantitative reconstitution of yeast RNA processing bodies

**DOI:** 10.1101/2022.08.13.503854

**Authors:** Simon L Currie, Wenmin Xing, Denise Muhlrad, Carolyn J Decker, Roy Parker, Michael K Rosen

**Author notes:** Correspondence (M.K.R.).

## Abstract

Many biomolecular condensates appear to form through liquid-liquid phase separation (LLPS). Individual condensate components can often undergo LLPS *in vitro*, capturing some features of the native structures. However, natural condensates contain dozens of components with different concentrations, dynamics, and contributions to compartment formation. Most biochemical reconstitutions of condensates have not benefitted from quantitative knowledge of these cellular features nor attempted to capture natural complexity. Here, we build on prior quantitative cellular studies to reconstitute yeast RNA processing bodies (P bodies) from purified components. Individually, five of the seven highly-concentrated P-body proteins form homotypic condensates at cellular protein and salt concentrations, using both structured domains and intrinsically disordered regions. Combining the seven proteins together at their cellular concentrations with RNA, yields phase separated droplets with partition coefficients and dynamics of most proteins in reasonable agreement with cellular values. The dynamics of most proteins in the reconstitution are also comparable to cellular values. RNA delays the maturation of proteins within, and promotes reversibility of, P bodies. Our ability to quantitatively recapitulate the composition and dynamics of a condensate from its most concentrated components suggests that simple interactions between these components carry much of the information that defines the physical properties of the cellular structure.

## Introduction

Biomolecular condensates are cellular compartments that lack an enclosing membrane barrier, yet are compositionally distinct from the surrounding environment (1, 2). Many condensates have liquid-like behaviors in cells, for example fusion and rounding due to surface tension, and rapid dynamics of key molecules. Some also have been shown to form through liquid-liquid phase transitions, based on sharp appearance/disappearance with changes in physical parameters (e.g. concentration, environmental conditions), dynamic behaviors and *in vitro*-*in vivo* correlations (3-11). Many studies have demonstrated that condensates can be formed *in vitro* using simple components from their natural counterparts—through phase separation of only a single, or a few, types of molecules. Such studies have demonstrated that relatively few components are required to form condensates, at least *in vitro (5-7, 12-20)*. Further, retention of mechanisms that regulate condensate formation and/or of condensate functions indicate that *in vitro* condensates can retain key features of the more complex cellular structures (5, 6, 18-29).

However, recent proteomic and sequencing studies have demonstrated that cellular condensates consist of tens to hundreds of different types of molecules, including proteins and nucleic acids (30-36). This complexity of composition is coupled to more complex mechanisms of formation than observed for individual molecules. For example, while for a given set of environmental conditions a single protein will phase separate at a specific concentration producing droplets of a specific concentration, in multi-component systems both the phase separation boundary and the composition of the condensate depend on specific nature of the interactions between the various components and thus on their relative and absolute concentrations (8, 9, 37, 38). Moreover, some components, by virtue of their interactions (valency, interaction partners, binding affinities), play more important roles in defining these parameters than others. Due to cooperativities between molecular interactions in multi-component systems, even when many/most components are known, it is not currently possible to quantitatively predict the compositions and physical properties of condensates from knowledge of their molecular parts and binary interactions thereof. As a step toward such a quantitative view, recent systematic surveys of cellular condensates to understand the concentrations of components and/or their requirements for condensate formation, have been reported (10, 39). Complex reconstitutions, containing all of the molecule types that are highly enriched in cellular condensates, provide an additional approach to a fuller understanding of the mechanisms that control the formation, properties, and functions of natural condensates.

RNA processing bodies (P-bodies) are archetypal biomolecular condensates consisting of RNA and proteins that are important in RNA metabolism, including enzymes with RNA-helicase (Dhh1), -decapping (Dcp2) and -exonucleolytic (Xrn1) activities (40-45). P bodies are thought to serve as sites of enhanced RNA degradation and/or of RNA and protein storage (30, 45-49). Previous biochemical reconstitutions have found that several different combinations of P-body proteins are sufficient to form condensates *in vitro* under various experimental conditions including: Dcp2, Edc3, Dcp1, and RNA (13, 21, 50); Dhh1, Pat1, and RNA (14, 51); and Dcp1, Dcp2, Lsm1-7, Pat1, and RNA (52). In some instances RNA reduced the saturation concentration (C_sat_) of P-body proteins (50), whereas in other cases RNA had no impact (21). Genetic studies indicate that individual P-body proteins can either promote or repress P-body formation, though no single protein is absolutely required to create the compartments (53, 54). Pat1, Edc3, and to a lesser extent Dhh1 promote yeast P-body formation, likely through the multivalent protein-protein and protein-RNA interactions that these proteins engage in (Table S1; Fig. S1)(14, 53, 55-57). In contrast, Dcp1, Xrn1, and Lsm1 repress P-body formation, as deletion of any of these components results in larger P bodies and/or higher partitioning of other components (39, 53, 58). For all three of these proteins, this repression may occur via their functions in RNA degradation, as RNA can contribute to P-body formation (53, 59), although RNA does not appear to be required for the maintenance of P-bodies (60). Dcp2 represses P-body formation in cells growing in mid-log phase by limiting the available deadenylated mRNAs required for P-body assembly, but promotes P-body formation under glucose starvation, or when decapping is blocked by deletion of Dcp1 (*dcp1Δ*), conditions that cause accumulation of large pools of deadenylated mRNAs independent of Dcp2 (36). This demonstrates how a single component of a condensate can have multiple inputs into condensate formation that are context dependent. These biochemical and genetic investigations suggest that P bodies are formed through multiple, partially-redundant interactions (54, 61).

Understanding the thermodynamics of condensate formation, including which interactions strongly contribute to it, requires knowledge about both the identity and the abundance of molecules within a given condensate (62). We recently performed a systematic analysis of *S. cerevisiae* P bodies and identified eight proteins (of 31 P body components examined) that *in vivo* are highly concentrated and have large partition coefficients (PC, concentration ratio of condensate to bulk cytoplasm) (Fig. 1A and S1)(39). This group included all of the aforementioned proteins that contribute to P-body formation in genetic experiments and Upf1. All other proteins, except Sbp1, exhibited at least twofold lower concentrations in P bodies, and all had at least twofold smaller PC values (compared to the lowest concentration and PC values of the other group)(39). The concentrations and dynamic properties of many (but not all) of the proteins were qualitatively correlated with the P body connectivity network, where network centrality was associated with higher concentration and lower dynamics (39).

**Figure 1.**
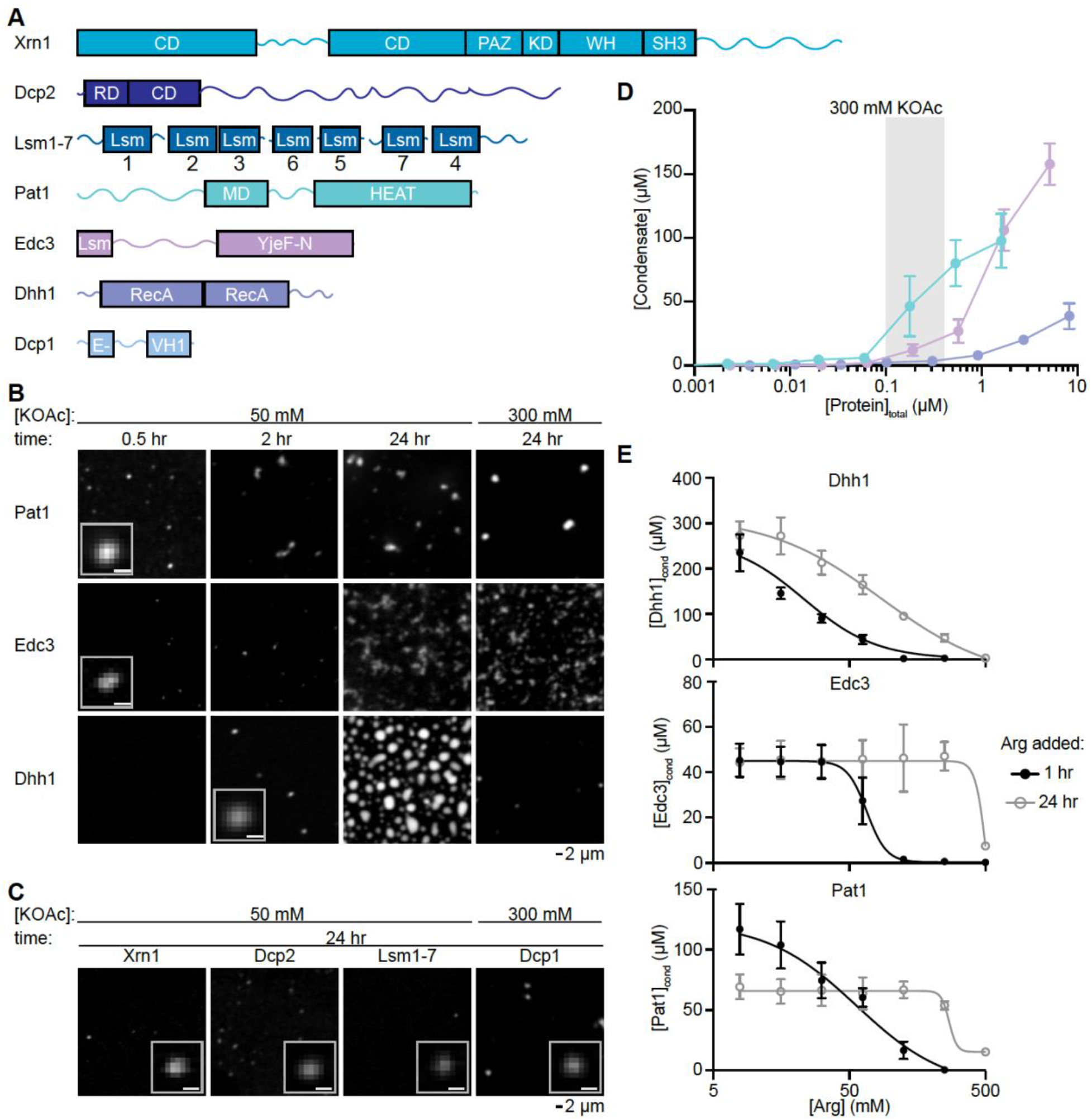
Individual P-body proteins phase separate at physiological conditions. (A) Domain structure of P-body proteins. Boxes indicate structured domains and lines represent IDRs. Abbreviations of domains are: CD, <u>C</u>atalytic <u>D</u>omain; PAZ, <u>P</u>iwi, <u>A</u>rgonaut, and <u>Z</u>wille domain; KD, <u>K</u>yprides, Ouzounis, Wouse <u>D</u>omain; WH, <u>W</u>inged <u>H</u>elix motif; SH3, <u>S</u>RC <u>H</u>omology <u>3</u> domain; RD, <u>R</u>egulatory <u>D</u>omain; Lsm, <u>L</u>ike <u>Sm</u> domain; MD, <u>M</u>iddle <u>D</u>omain; HEAT, <u>H</u>untingtin, <u>E</u>longation factor 3, protein phosphatase 2<u>A</u>, and <u>T</u>OR1 domain; YjeF-N; <u>YjeF N</u>-terminal domain; RecA, <u>RecA</u>-like domain; EVH1, <u>E</u>na/<u>V</u>asp <u>H</u>omology domain <u>1</u>. (B) Fluorescent micrographs for individual P-body proteins in a buffer at pH 7 with low (50 mM, left) and physiologically-relevant (300 mM, right) KOAc concentrations. Scale bar for inset corresponds to 0.5 *μ*m. Pat1 is at 0.5 *μ*M and Edc3 and Dhh1 are at 1 *μ*M total concentration. Proteins are labeled with EGFP. (C) Fluorescent micrographs for Dcp2, Lsm1-7, and Dcp1 at 1 *μ*M and Xrn1 at 0.4 *μ*M total concentration, and with indicated KOAc concentrations that promoted phase separation. Dcp1 and Dcp2 are labeled with EGFP, Lsm1-7 and Xrn1 are labeled with Alexa Fluor 488. (D) Graph of condensate concentration vs total protein concentration for Pat1, Edc3, and Dhh1 in a buffer at pH 7 with 300 mM KOAc. Gray rectangle corresponds to the estimated cellular concentration of these proteins. (E) Condensate protein concentration as a function of arginine added for Dhh1 (left), Edc3 (middle), and Pat1 (left). Black closed circles and gray open circles correspond to arginine added 1 hour and 24 hours after condensate initiation, respectively. Before adding arginine condensates were formed with 50 mM KOAc present. See also Fig. S2-S6.

Here we sought to understand whether and how the most concentrated proteins in P bodies could produce condensates with RNA *in vitro* that have properties similar to their cellular counterparts. We find that five proteins, Pat1, Dhh1, Edc3, Dcp1, and Xrn1 can form homotypic condensates at cellular protein and salt concentrations. Molecular dissections indicate that condensate formation by these proteins occurs through interactions of both folded domains and predicted disordered regions. Acidic pH, mimicking the dormancy state during which large yeast P-bodies form (63, 64), promotes condensate formation by enhancing homotypic phase separation and enhancing the affinity of P-body proteins for RNA. Combining all proteins with RNA at their cellular concentrations, and under cellular environmental conditions, produces heterotypic condensates whose PC values and dynamics are quantitively similar to those of cellular P bodies. Thus, the capacity to generate these cellular structures may reside critically in the most concentrated components (which is consistent with the prior genetic analyses). Comparisons of the single-component and eight-component condensates suggest that competing homotypic and heterotypic interactions are integrated within the multicomponent droplets. Our quantitative reconstitution is a valuable system for future examinations of the formation and biochemical function(s) of P bodies, and provides a general framework for understanding the formation of multicomponent condensates.

## Results

### Individual P-body proteins phase separate under physiologic conditions

We examined seven of the eight proteins that are highly concentrated in *S. cerevisiae* P bodies: Dcp1, Dcp2, Dhh1, Edc3, Lsm1-7, Pat1, and Xrn1 (Fig. 1A)(39). We did not include Upf1 because this protein does not affect the targeting of most normal mRNAs to P bodies, but rather only a subset of those undergoing aberrant translational termination and nonsense-mediated decay (NMD)(65). Therefore Upf1 is more of an accessory factor, as opposed to a core factor, of P-bodies. These proteins possess many features that have been associated with condensate formation including multivalent protein-protein and protein-RNA interactions, and intrinsically disordered regions (IDRs)(Fig. S1-3). Full length Dcp1, Dcp2, Dhh1, Edc3, and Pat1 were expressed individually in bacteria with N-terminal maltose binding protein (MBP) and C-terminal His_6_ tags. Double affinity purification yielded full length reagents (Fig. S4A). To increase solubility we retained the MBP fusion throughout the purification and used TEV cleavage of both tags to initiate condensate formation (Fig. S4B and Methods). P-body proteins with Enhanced Green Fluorescent Protein (EGFP) inserted immediately before the C-terminal TEV cleavage site were also expressed and purified. We used previously established expression plasmids and purification protocols for the Lsm1-7 complex and Xrn1 (40, 66, 67). Alexa Fluor 488-conjugated versions of Lsm1-7 and Xrn1 were also purified for quantitative analyses of these proteins.

We initially examined individual P-body proteins labeled with EGFP or Alexa Fluor 488 (see Methods) at 0.4 to 1 µM concentration and pH 7, similar to the cytoplasm of unstressed yeast in log phase growth (64). Each of them formed micron-sized condensates with distinct properties under different salt conditions (Fig. 1B and 1C). Pat1, Edc3, and Dhh1 rapidly formed small spherical condensates in 30 minutes to a few hours after TEV addition, with partition coefficients (ratio of droplet to bulk concentration) greater than 40 in low salt conditions (50 mM KOAc). Comparatively, Dcp1, Dcp2, Lsm1-7, and Xrn1 slowly formed condensates over many hours after TEV addition, with lower partition coefficients (less than 5). Physiologically-relevant salt concentrations [300 mM KOAc: (68, 69)] eliminated Dcp2, Lsm1-7, and Xrn1 condensates, reduced the size and number of Dhh1 condensates, had minimal impact on Edc3 and Pat1 condensates, and slightly enhanced Dcp1 condensates. EGFP alone does not form condensates under the same experimental conditions (Fig. S4C), suggesting that the P-body proteins are driving condensate formation. The 0.4 to 1 µM protein concentrations in these assays were roughly two- to tenfold higher than estimated cellular concentrations of these molecules (Fig. 1B, 1C and S5)(70). However, we still observed condensate formation at pH 7 with physiological protein and salt concentrations for Pat1, Edc3, and Dhh1 (Fig. 1D), consistent with these proteins playing important roles in forming P-bodies in cells (14, 53, 55, 56).

Although Pat1, Edc3 and Dhh1 all initially formed small spherical homotypic condensates, different proteins transitioned to distinct morphologies, with different physical properties, over time. Dhh1 condensates grew into larger objects that are mostly spherical, whereas Pat1 and Edc3 formed aspherical networks of condensates (Fig. 1B). One interpretation of this result is that homotypic condensates of Edc3 and Pat1 rapidly mature into structures that cannot fuse and round, while those of Dhh1 remain more dynamic (see below), with surface tension that produces spherical shape. One approach to probe the maturation of condensates is to examine whether they become more resistant to dissolution over time. To develop such an assay we screened for molecules that disrupt homotypic condensate formation. Dhh1 condensates were disrupted by a variety of molecules, including salt (NaCl), glycerol, and arginine; in contrast, Edc3 and Pat1 were selectively disrupted by only arginine (Fig. S6A). Although the concentration of arginine required to dissolve homotypic condensates varied by protein, we observed that arginine more potently disrupted each of the homotypic condensates when aged for 1 hour as compared to 24 hours (Fig. 1E and S6B). These data suggest that in the absence of other factors (e.g. RNA, see Fig. 6 and Discussion) homotypic P-body protein condensates change over time into a less reversible state.

### Acidic pH enhances homotypic condensate formation and protein-RNA interactions

P bodies form in response to cellular stresses that elicit cell cycle arrest and entry into a dormant state (59). This transition results in several changes to the cellular environment including acidification of the cytosol (63, 64). We therefore wondered whether P-body proteins might respond to changes in pH. Indeed, at 0.4 to 1.0 µM concentration all P-body proteins exhibited more numerous condensates and higher PCs at pH 5.8 in lower salt conditions (50 mM KOAc; Fig. 2A and S7). The magnitude of pH-sensitivity of PC values ranged from one hundred-fold for Dcp1 to two-fold for Pat1 (Fig. 2B, S7B-C). Higher salt (300 mM KOAc) reduced the pH-sensitivity of PCs for all proteins except for Dhh1, suggesting that ionizable groups may be responsible for sensing the change in pH. We hypothesized that histidine residues may play an important role in pH-sensitive condensate formation because the pH values that we tested bracket the pK_A_ of the imidazole side chain (71). Analysis of a previously-published crystal structure of Dcp1 (72) suggested that one particular histidine residue, H206, might be important for Dcp1 condensate formation as it is centrally located within the homodimer interface for both Dcp1 molecules (Fig. 2C). Indeed, mutation of H206 to alanine specifically inhibits oligomerization and homotypic condensate formation by Dcp1 at pH 5.8, whereas mutation of other histidine residues had less impact (Fig. 2D-E and S8). In contrast to this single important histidine in Dcp1, truncations and point mutants in Dcp2 (described further below) indicated that elements distributed in both the structured N-terminal domains and in the C-terminal IDR of the protein contribute to pH-sensitive condensate formation (Fig. 2F-G and S9A). There is a correlation between the frequency of histidine residues and the pH-sensitive partitioning (at 300 mM KOAc) of P-body proteins other than Dcp1 (Fig. 2H-I and S9B). In contrast, the isoelectric points of proteins do not correlate with the pH sensitivity of homotypic PCs (Fig. S9C). This suggests that similar to Dcp2, multiple distributed histidine residues likely contribute to pH-sensitive LLPS behaviors of other P-body proteins as well. Together, the data suggest that the protonation of histidine residues at acidic pH may promote the oligomerization and homotypic condensate formation of P-body proteins. Further *in vivo* experiments will be necessary to determine which of these proteins play the greatest roles in pH sensing by P-bodies in yeast.

**Figure 2.**
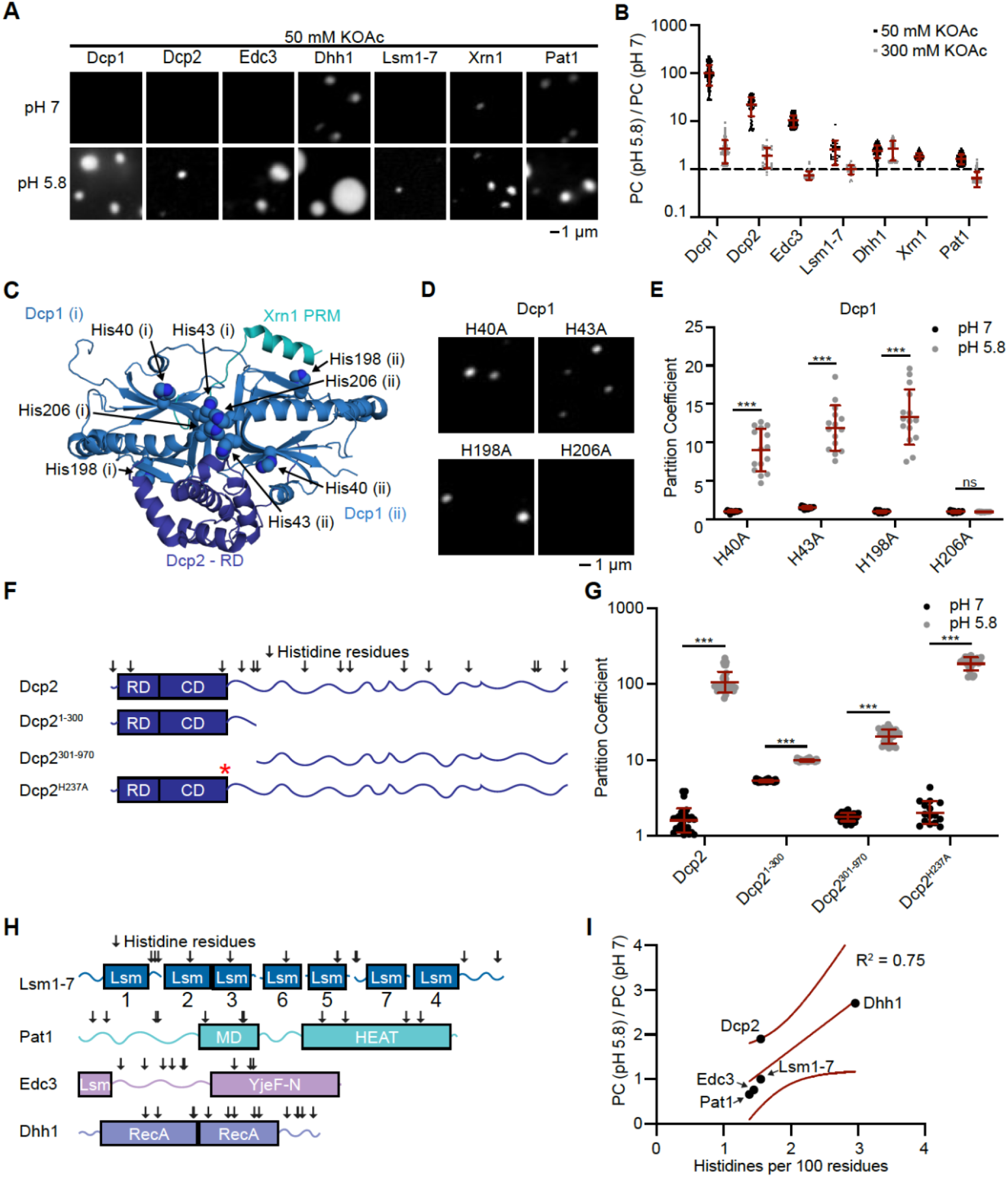
Acidic pH promotes homotypic phase separation and RNA binding of P-body proteins. (A) Representative micrographs for individual P-body proteins at pH 7 (top) or 5.8 (bottom) and 50 mM KOAc. Proteins labeled as described in Fig. 1. (B) Ratio of partition coefficients for individual P-body proteins at pH 5.8 and pH 7. Data points are partition coefficients for individual condensates from two replicate experiments. (C) Crystal structure of Dcp1 dimer (72) annotated with mutated histidine residues. Interactions with Xrn1 (100) and Dcp2 (101) were modeled onto the structure. (i) and (ii) indicate distinct Dcp1 molecules. (D) Representative micrographs of Dcp1 histidine mutants at pH 5.8 and 300 mM KOAc. Dcp1 mutants are labeled with EGFP. (E) Partition coefficients of Dcp1 histidine mutants with 300 mM KOAc, pH 7 (black) and pH 5.8 (gray). Data points are partition coefficients for individual condensates from two replicate experiments. (F) Schematic of Dcp2, truncations, and point mutations tested. Arrows above Dcp2 indicate histidine residues. Red asterisk indicates location of His237 point mutations. (G) Partition coefficients for Dcp2 truncations and mutations with 300 mM KOAc, pH 7 (black) and pH 5.8 (gray). Data points are partition coefficients for individual condensates from two replicate experiments. Dcp2 truncations and mutants are labeled with EGFP. (H) Location of histidine residues in Lsm1-7, Pat1, Edc3, and Dhh1. (I) Ratio of partition coefficients (pH 5.8 / pH 7) with 300 mM KOAc versus the frequency of histidine residues (number of histidine residues per 100 amino acids) for all P-body proteins that were tested. Solid line indicates linear fit and dotted lines indicate 95% confidence interval. See also Fig. S7-S9.

Having established an importance for pH in homotypic interactions, we queried whether pH might also regulate protein interactions with RNA. Previous studies have demonstrated that dimerization can increase affinity of RNA binding proteins for RNA through avidity effects (73). Therefore we reasoned that the pH-enhanced oligomerization of P-body proteins might increase their affinity for RNA. We used Electrophoretic Mobility Shift Assays (EMSAs) to measure the equilibrium dissociation constants (*K*_*D*_) for binding of each protein to an RNA oligonucleotide at pH 7 and pH 5.8 (Fig. 3A and S10). For these experiments we used a synthetic 60-nucleotide RNA consisting of portions of the RPL41A mRNA, a known resident of yeast P bodies (Fig. S11)(74). Acidic pH increased the RNA-binding affinity of all proteins, except for Dcp1 which did not appreciably bind to RNA at either pH (Table S2; Fig. 3B-C and S10). The Dcp2-RNA interaction was greater than 16-fold tighter at pH 5.8 versus pH 7.0, whereas the other proteins showed more modest pH-dependent enhancements (two- to fivefold). The Hill coefficients for all individual proteins were higher at pH 5.8, indicating that RNA binding is more cooperative under acidic conditions (Table S2). The enhanced cooperativity of RNA-binding at pH 5.8 may be a result of the higher oligomerization state of P-body proteins in these conditions. These data indicate that the RNA-binding of P-body proteins is enhanced at acidic pH (Fig. 3D).

**Figure 3.**
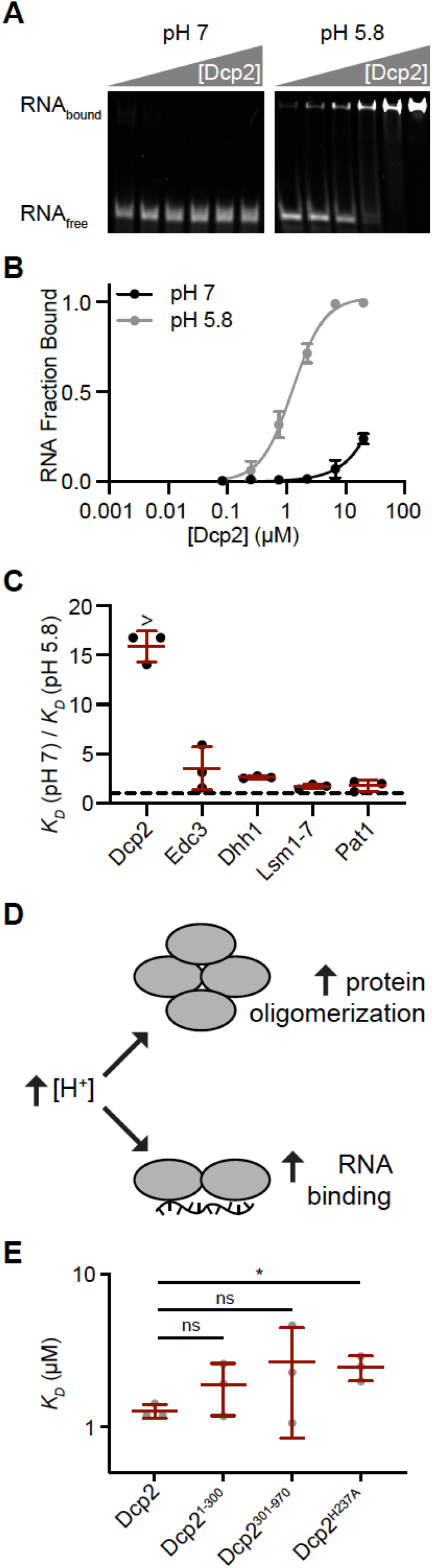
Acidic pH promotes P-body proteins binding to RNA. (A) Representative EMSA gels for Dcp2 binding to RPL41A RNA with 300 mM KOAc, pH 7 (left) and 5.8 (right). (B) Quantitation of Dcp2 binding to RPL41A RNA at pH 7 (black) and pH 5.8 (gray). Data points are the average and standard deviation from three replicate experiments. Lines show fits to the Hill equation. (C) Ratio of equilibrium dissociation constants, *K*_*D*_, for individual P-body proteins at pH 7 and 5.8. Ratios for Dcp2 are lower limits as RNA binding at pH 7 was too poor to be accurately quantified. (D) Cartoon summarizing that acidic pH promotes protein oligomerization and homotypic LLPS (top) and tighter binding to RNA (bottom). (E) Equilibrium dissociation constants for Dcp2 truncations and point mutants binding to RPL41A RNA with 300 mM KOAc. See also Tables S2 and S3; Fig. S10-S11.

We next examined the location of pH-sensitive RNA-binding elements within Dcp2 since it exhibited the largest enhancement in RNA-binding affinity. A previous study suggested that a basic patch on the surface of the regulatory and catalytic domains (Dcp^1-300^) is important for RNA binding (Fig. 2F and S9A)(75). Within this patch we mutated histidine 237 to alanine (Dcp2^H237A^). This mutation resulted in a modest two-fold decrease in the affinity of Dcp2 for RNA at pH 5.8 (Table S3; Fig. 3E), suggesting that additional pH-sensitive RNA-binding elements are present within Dcp2. Indeed, we found that both the structured domains (Dcp2^1-300^) and the IDR (Dcp2^301- 970^) of Dcp2 each bind to RNA with similar affinity as full-length Dcp2 (Table S3; Fig. 2F and 3E). Furthermore, the RNA binding of both regions are also pH sensitive (Table S3). The similar RNA-binding affinities of full-length Dcp2 and each of these regions indicates that the RNA-binding of Dcp2 may be autoinhibited, as has been previously reported for Dcp2 RNA-decapping activity (76). Thus, similar to the condensate-forming elements of Dcp2, the pH-sensitive RNA-binding elements are also distributed in both the N-terminal structured domains and the C-terminal IDR.

### Structured domains and IDRs contribute to homotypic LLPS of P-body proteins

We next sought to determine which regions of individual P-body proteins are responsible for homotypic condensate formation. We created truncations that separated structured domains from IDRs, using truncation boundaries from previous studies where available (Fig. 4A-E) (42, 55, 57). We used condensate-promoting buffer conditions of acidic pH (5.8) and low salt (50 mM KOAc) because we hypothesized that truncations may have reduced condensate forming capacity relative to full-length proteins. Interestingly, every protein tested contained multiple regions that are sufficient to form condensates (Fig. 4F-J and S12-16). Only the N-terminal IDR of Pat1 and the structured Lsm domain of Edc3 did not form condensates at any tested concentration. Most P-body truncations formed spherical structures, with the exception of the disordered linker of Dcp1 (Dcp^182-129^) whose formations resembled interconnected fibrils (Fig. 4J and S16). By comparing each full-length protein with its constituent regions, we can infer how the condensate-forming behavior of different regions contributes to that of each full-length protein (Fig. 4A-E and S12- 16). Pat1 has a similar C_sat_ as the middle domain, Pat1^241-422^, yet Pat1^241-422^ has a higher condensed phase concentration (Fig. 4L and S13B). Pat1^423-796^ also self assembles, but only at the highest concentrations tested (Fig. 4L inset and S13). Together these data suggest that Pat1^241-422^ contributes most strongly to phase separation whereas Pat1^1-240^ and/or Pat1^423-796^ modulate the degree of concentration within the condensate. For Dcp2 and Edc3 the full-length proteins exhibit lower C_sat_ and lower condensed phase concentrations compared to each of their individual regions (Fig. 4K, 4M, S12, and S14). Edc3^1-66^ did not phase separate at any concentration tested (Fig. 4H, 4M inset, and S14). Individually, Edc3^67-282^ and Edc3^283-551^ both phase separate, suggesting that these two regions contribute to phase separation of the full- length protein (Fig. 4H, 4M inset, and S14). Dhh1 and Dcp1 have similar condensed phase concentrations as their structured domains alone, but the adding the IDR drives the C_sat_ lower for Dcp1 (Fig. 4N, 4O, S15, S16). These data suggest that in Dcp1, Dcp2, and Edc3 multiple regions synergize to lower their C_sat_ (Fig. 4A and 4C-E). The region that contributes more strongly to condensate formation is a structured domain for Pat1, Dhh1, and Dcp1, whereas structured domains and IDRs are both major contributors to Dcp2 and Edc3 condensate formation. Together, the data suggest that multiple regions act together to drive homotypic condensate formation of full-length P-body proteins, and that both structured domains and IDRs are important contributors to this process.

**Figure 4.**
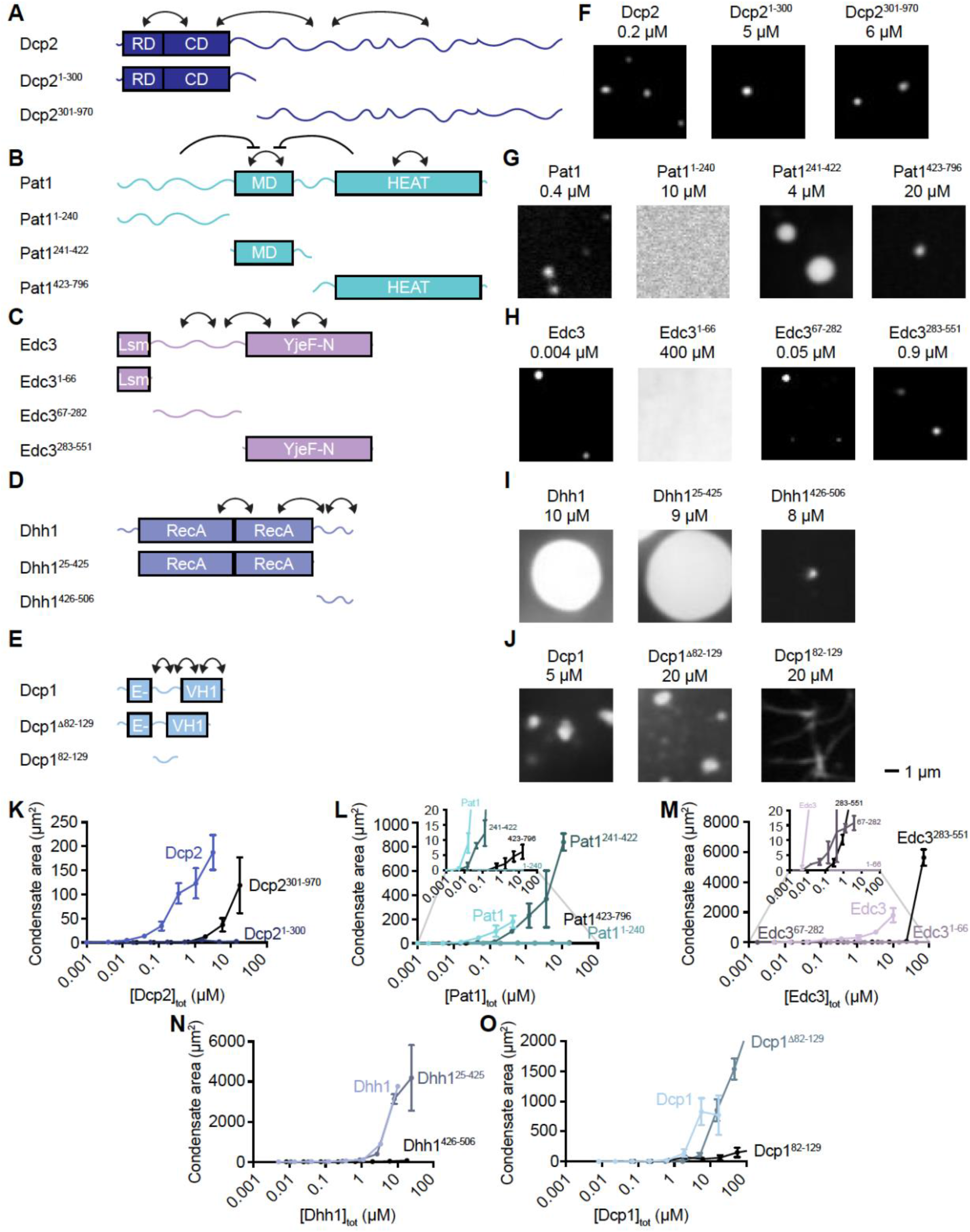
Structured domains and IDRs contribute to homotypic condensate formation. (A-E) Schematic of full-length and truncated P-body proteins. Arrows indicate inferred relationships between protein domains and/or IDRs as suggested from data in K-O. Arrows within a domain/IDR indicate that homotypic interactions within that region are sufficient to drive condensate formation. Arrows between domains/IDRs represent our interpretation that these domains synergize in the full-length protein to drive lower saturation concentrations and/or higher condensed phase concentrations; inhibitory arrows represent a negative relationship between these domains. Arrows between domains/IDRs do not necessarily indicate that a physical interaction occurs between these regions, as other explanations are possible, though such interactions are known between Dcp2^1-300^ and Dcp2^301-970^, Pat1^241-422^ and Pat1^423-796^, and Edc3^67-282^ and Edc3^283-551^ (50, 57, 76). (F-J) Representative images for a single concentration of each full-length and truncated P-body protein. Condensates were formed with 50 mM KOAc and pH 5.8. All truncations were labeled with EGFP. (K-O) Plot of condensate protein concentration vs total concentration for each full-length and truncated P-body protein. Insets are shown for Pat1 and Edc3 for clarity of datapoints with small area values. See also Fig. S12-S17.

There is reasonable agreement between P-body protein regions that exhibit LLPS *in vitro* (Fig. 4), and contribute to P-body formation *in vivo* (Fig. S17). Our *in vitro* results are consistent with a synergy between multiple regions within Dcp2, and within Dhh1, contributing to P-body formation in cells (23, 39). Furthermore, our results are consistent with the importance of Edc3^283-551^ and Pat1^423-796^ contributing to P-body formation (55, 56). However, there are discrepancies for some P-body protein regions (Fig. S17). Edc3^1-66^ does not form homotypic condensates, yet is important for P-body formation *in vivo* (55). This discrepancy is likely due to heterotypic interactions between Edc3^1-66^ and helical leucine motifs in Dcp2 that contribute to P-body formation (13, 39). Dcp1 forms homotypic condensates *in vitro*, but represses P-body formation in cells by activating the RNA-decapping of Dcp2 (53) which is an activity not present in homotypic Dcp1 condensates. Edc3^67-282^ and Pat1^241-422^ form homotypic condensates *in vitro*, but are not important for P-body formation *in vivo* (55, 56). We do not currently understand these latter discrepancies, however there are some inherent limitations to our *in vitro* - *in vivo* comparisons for P-body protein regions (Fig. S17). Since the threshold concentration for Edc3^67-282^ is at least tenfold higher than Edc3 *in vitro* (Fig. S14), one possibility is that Edc3^67-282^ may contribute to P body formation, but only at higher expression levels than were tested *in vivo* (55). Furthermore, our *in vitro* assays are quantitative whereas some of the previous cellular data is more qualitative in scoring for P-body formation based on one or two molecular markers (55, 56). Thus, protein regions with weaker contributions to phase separation, such as Edc3^67-282^, may be missed by qualitative examinations in cells. While investigating the LLPS of isolated protein regions informs on the importance of homotypic interactions, this approach does not always correlate with condensate formation in cells. For example, regions whose contributions to P-body formation are mediated via heterotypic interactions with other components will be missed by this approach (see also Fig. 6 and Discussion).

### Proteins have similar partitioning and dynamics in reconstituted and cellular P bodies

Having characterized homotypic condensates of individual P-body proteins, we next examined heterotypic condensates combining all P-body proteins and RNA. All of the proteins and RNA were mixed together and incubated for 2 hours at 30 °C before cleaving protein tags with TEV to initiate condensate formation. Initiating condensate formation with less preincubation of components resulted in heterogeneous condensates and multiphase condensates with complex architectures that have not been observed in cellular P bodies (Fig. S18). Condensates were incubated for twenty-four hours before imaging, unless stated otherwise. We used Pat1-mCherry and RNA-AlexaFluor647 as markers and examined colocalization with GFP-tagged or AlexaFluor488-conjugated versions of other P-body proteins. We used a fluorescence intensity cut-off of threefold above background for each fluorophore, and assigned the center of mass for condensates in each of the microscope channels. Condensates with centers of mass within 1 µm were considered to be overlapping to account for slight discrepancies in the designation of the center of mass and for stage drift during imaging.

Initially we examined the full collection of molecules under acidic conditions (pH 5.8), as the previous inventory of P body components in wild type yeast was determined during glucose starvation, where intracellular pH is approximately 5.8 (39, 64). Additionally, under these conditions LLPS of the individual components is favored and RNA binding affinity is tighter (see above), providing technical advantages to the assay. In these acidic conditions, Pat1 frequently overlaps with all other P-body proteins (> 90%), and all P-body proteins frequently overlap with RNA (> 90%: Fig. 5A-B and S19). These results indicate that at pH 5.8 the vast majority of condensates contain all eight components. EGFP by itself did not partition into heterotypic condensates (Fig. S20), suggesting that formation of, and recruitment into, condensates is dependent on specific protein-protein and protein-RNA interactions. Furthermore, all condensates appeared to have homogeneous fluorescence (at the resolution limit of our confocal microscope) regardless of which components were labeled. Thus, under conditions favoring protein-protein and protein-RNA interactions, all eight P-body components together form condensates.

**Figure 5.**
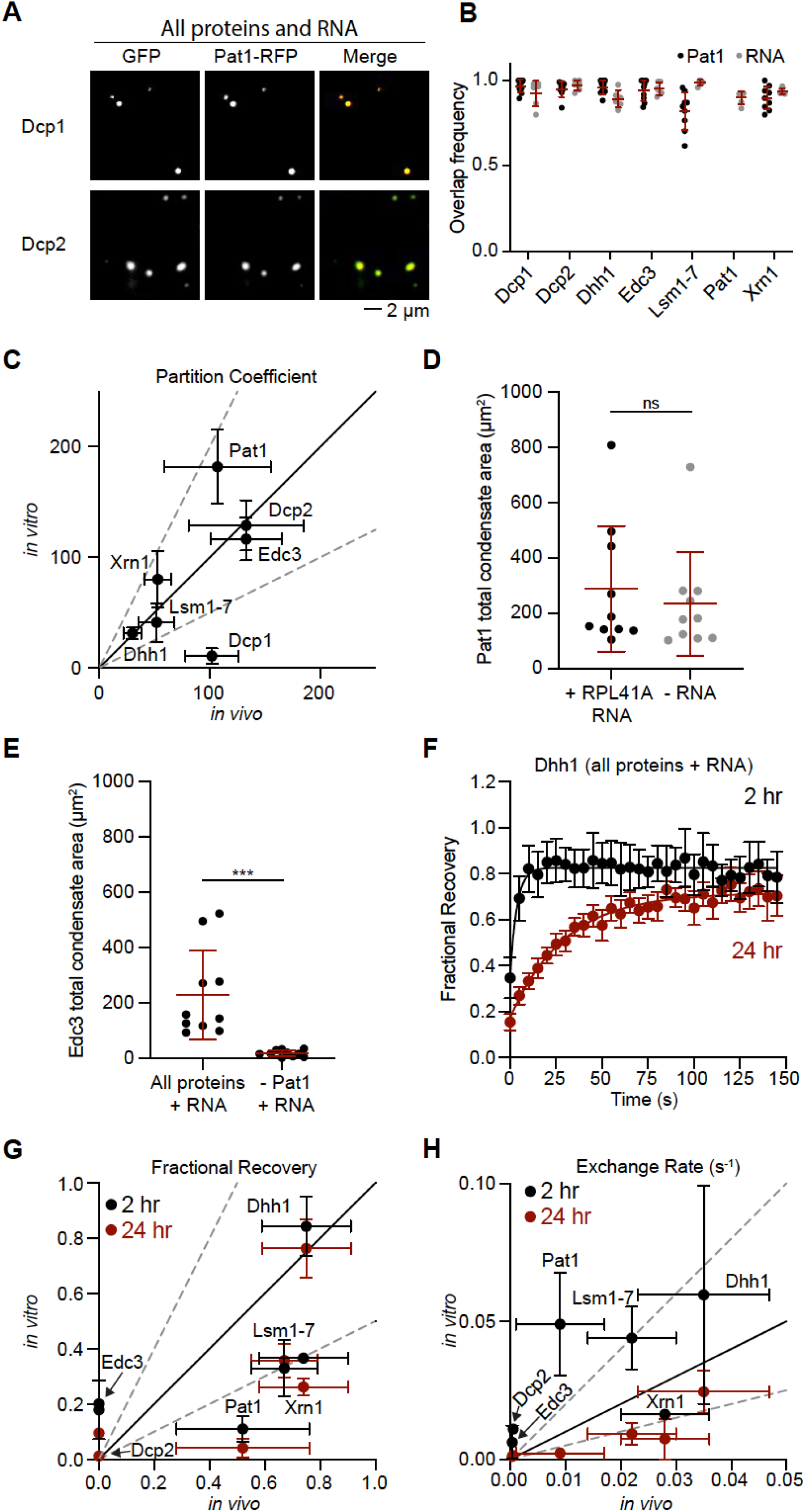
Quantitative reconstitution of P-bodies reveals two classes of scaffolds. (A) Representative micrographs examining overlap between Pat1 and Dcp1 (top) or Dcp2 (bottom). All P-body molecules are included in the samples at estimated cellular concentrations. All experiments in Fig. 5 are at pH 5.8 and 300 mM KOAc. (B) Fraction overlap between different P-body proteins with Pat1 (black) and RNA (gray). Data points correspond to the mean values for individual micrographs from at least two replicate experiments. (C) Comparison of partition coefficients for P-body proteins *in vitro* values (this study) with *in vivo* values from a previous study (39). Solid black line indicates equivalent values and dashed gray lines indicate twofold differences. Data points and error are the mean and SD from at least four replicate experiments. (D) Comparison of condensate area per micrograph, using Pat1 as a marker, for the reconstitution with RPL41A RNA, left, and without RNA, right. (E) Comparison of condensate area per micrograph, using Edc3 as a marker, for the reconstitution with all proteins and RNA, left, and in the absence of Edc3, right. (F) FRAP recovery curves for Dhh1-EGFP in condensates with all P-body proteins and RNA. Experiment was performed after condensates had formed for 2 hours (black) and 24 hours (red). Data points and error bars correspond to the mean and SD from fifteen condensates in a single experiment. At least two replicate experiments were performed. (G) Comparison of fractional recovery values for each P-body protein with *in vivo* values (39). Black solid line is where values are equivalent and dotted gray lines correspond to twofold differences. Data points and error bars correspond to the mean and SD from two replicate experiments. (H) Comparison of exchange rate values for each P-body protein with *in vivo* values (39). Data presented as in (G). See also Table S4; Fig. S5 and S18-S26.

LLPS can be exquisitely sensitive to parameters such as pH, temperature, and protein concentrations (77). In order to more rigorously examine which experimental conditions generate *in vitro* P-bodies that most closely mimic their *in vivo* counterparts, we measured partition coefficients for each P-body protein while varying pH, temperature, or total protein concentrations. We compared partition coefficients for these proteins with previously measured values in *S. cerevisiae (39)*. Using the estimated cellular protein concentrations (Fig. S5), at pH 5.8, and incubating at 30 °C resulted in the closest match to cellular P bodies (Table S4; Fig. 5C and S21). Indeed, under these conditions the partition coefficients for most proteins were within twofold of their *in vivo* values, with the exception of Dcp1. Changing to neutral pH 7, incubating the reactions at 4 °C, or using equal protein concentrations (150 nM) for every protein all resulted in greater deviation from cellular values (Fig. S21). Each perturbation affected a unique subset of proteins, with Dcp2 being the lone protein whose partitioning was strongly impacted by all changes. Incubating the reactions at 4 °C affected the partitioning of all proteins except for Dhh1. Incubation at pH 7 resulted in a distinct composition of condensates with RNA, Dcp1, and Xrn1 less frequently overlapping with the other P-body proteins (Fig. S22) Thus, we conclude that the *in vitro* experimental conditions of pH 5.8, 30 °C, and cellular protein concentrations most accurately recapitulate the cellular stoichiometry of P-bodies.

We next investigated the role of RNA in our reconstitution. The *in vitro* P bodies described above were formed in the presence of RPL41A RNA, a 60 nucleotide RNA with secondary structure (Fig. S11). Removing this RNA from the reconstitution had little impact on condensate area (Fig. 5D). We tested additional RNAs including a 10 nucleotide single stranded RNA [RNA10, (78)], full-length Mating Factor A (MFA2) mRNA which is 348 nucleotides long and localizes to cellular P bodies (59), and total RNA from yeast. None of these RNAs strongly impacted condensate formation or partitioning of Pat1 into the structures (Fig. S23A-B). All RNAs were more enriched in condensates at pH 5.8 than at pH 7.0 (Fig. S23C-D), consistent with higher affinity of several of the proteins for RNA under acidic conditions (Fig. 3). The enrichment of the defined RNA species in reconstituted P bodies correlates with their length (Fig. S23E). Thus, in our reconstitution, RNA does not strongly contribute to P-body formation, but rather is recruited in a length-dependent manner.

Genetic studies have shown that Pat1 is an important contributor to P-body formation in cells (53, 56). To examine whether Pat1 is similarly important to our reconstitution we held all other P-body components at their cellular concentrations but left out Pat1. The absence of Pat1 strongly impaired condensate formation and reduced partitioning of all other molecules into the few remaining condensates (Fig. 5E and S24). These results confirm the importance of Pat1 for P-body formation and suggest that our reconstitution is dependent on similar molecular interactions as cellular P bodies.

Finally, to compare the dynamics of the reconstituted P-bodies with their cellular counterparts, we monitored fluorescent recovery after photobleaching (FRAP) for each P-body protein in the context of the full complement of components. In preliminary experiments we observed that ATP is required for Dhh1 to have similar dynamics *in vitro* as in cells, likely due to the role of the nucleotide hydrolytic cycle in binding and release of RNA by the protein (Fig. S25) (39). We monitored the recovery of each protein at an early timepoint (2 hours) and at a late timepoint (24 hours) (Fig. 5F and S26). At the later timepoint most proteins demonstrated reduced fractional recovery and all proteins exhibited slower exchange rates (Fig. 5G-H and S26). Importantly, with the lone exception of Pat1, P-body proteins exhibited fractional recoveries and exchange rates comparable (within a factor of 2-4) to those observed in cells (39). The fractional recovery of Pat1, however, was lower in our *in vitro* reconstitution than in cells. Collectively, these data indicate that our *in vitro* reconstitution with seven proteins and RNA is sufficient to recapitulate the partitioning and dynamics observed in cellular P bodies for most proteins.

### Homotypic and heterotypic condensates have different physical properties

Having established experimental conditions (pH 5.8, 300 mM KOAc, 30 °C, cellular concentrations of proteins) that produce native-like P bodies, we next queried whether homotypic condensates of individual proteins could form under these conditions (note that homotypic condensates in Fig. 2 were formed with higher protein concentrations, lower salt concentrations, and at room temperature). Individually, Dcp2, Lsm1-7, and RNA did not form homotypic condensates, indicating that their recruitment into these compartments is heterotypic in nature (Fig. 6A and S27). In contrast, Dhh1, Edc3, Xrn1, Dcp1, and Pat1 all formed homotypic condensates, raising the question of whether there are any differences between homotypic condensates made up of a single P-body protein and heterotypic condensates that include all of the P-body proteins and RNA. All of the proteins that phase separated individually exhibited a higher partition coefficient in homotypic condensates than in heterotypic condensates (two- to tenfold), though this difference was only statistically significant for Edc3, Dcp1, and Pat1 (Fig. 6A). These results suggest that heterotypic interactions between all molecules promote a single multicomponent condensate, instead of distinct homotypic condensates. Heterotypic interactions recruit individual components that do not create condensates on their own (Dcp2, Lsm1-7, and RNA), and some heterotypic interactions temper homotypic interactions that drive Dhh1, Edc3, Xrn1, Dcp1, and Pat1 into condensates, likely through competitive binding. These results are conceptually consistent with recent descriptions of multicomponent LLPS through the framework of polyphasic linkage (see Discussion) (38, 79, 80).

**Figure 6.**
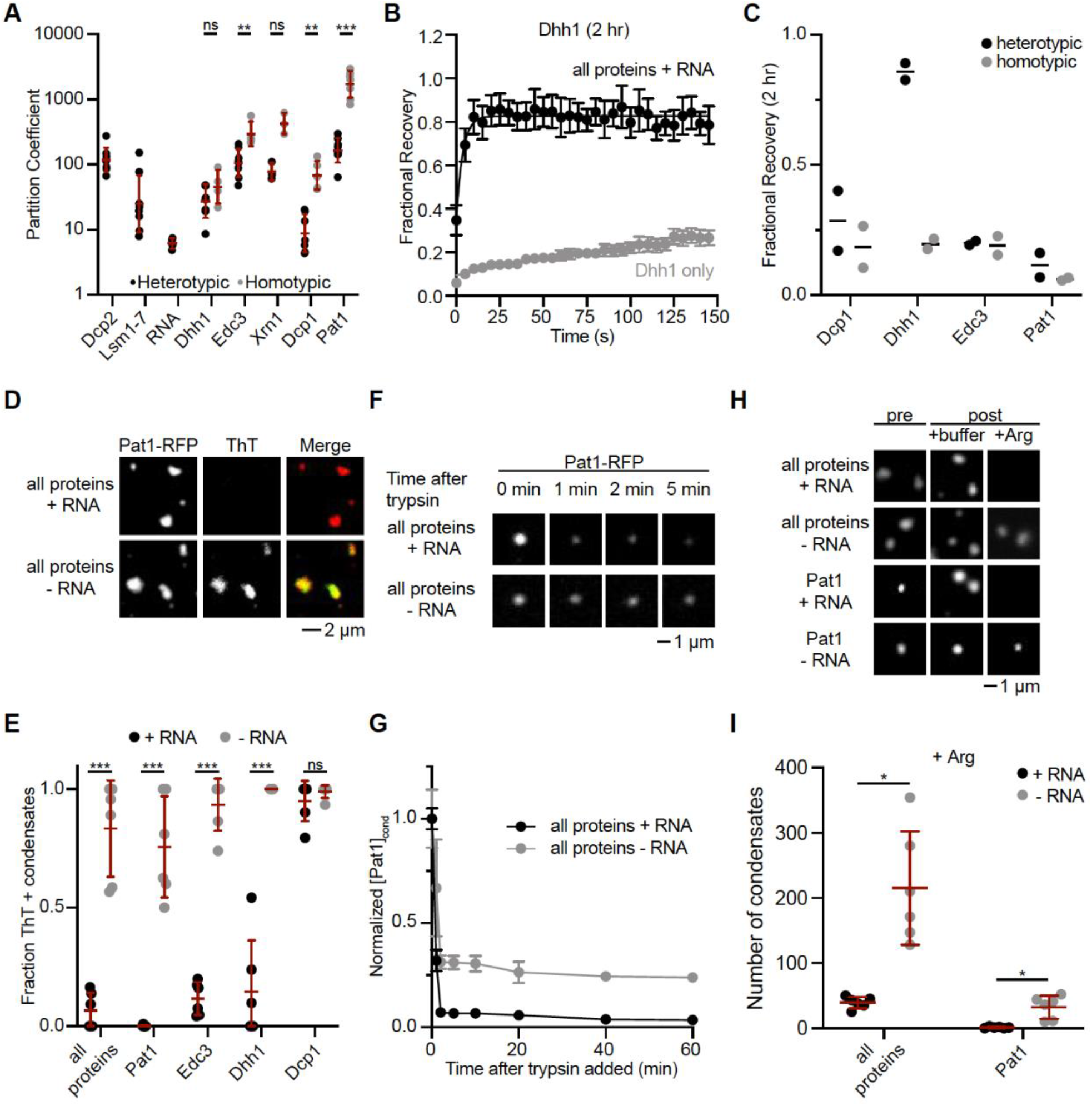
RNA prevents the maturation of P-body proteins and promotes reversibility. (A) Comparison of partition coefficients for P-body molecules in heterotypic (black) and homotypic (gray) condensates. Data points are the mean values from at least four replicate experiments. All experiments in Fig. 6 were conducted with 300 mM KOAc. (B) FRAP recovery curves for Dhh1 in heterotypic condensates (black) and in homotypic condensates (gray). Experiment was performed after condensates had formed for 2 hours. Data points and error bars correspond to the mean and SD from fifteen condensates in a single experiment. At least two replicate experiments were performed. (C) Fractional recovery values for P-body proteins in heterotypic (black) and homotypic (gray) condensates. Data points and line correspond to two replicate experiments and the mean. (D) Representative micrographs for condensates with all proteins and with RNA (top) or without RNA (bottom). Condensates were imaged with Pat1-RFP (left) and with ThT (middle). (E) Fraction of ThT positive condensates with RNA (black) and without RNA (gray). The proteins included in each condensate are listed on the x-axis. (F) Time course of micrographs with trypsin digestion of condensates. (G) Relative quantification of Pat1 intensity in condensates during trypsin digestion. (H) Representative micrographs for formation and dissolution of condensates with arginine. (I) Pat1 partition coefficient after arginine dissolution, normalized to a buffer control. See also Fig. S27-S31.

We next used FRAP experiments to determine whether homotypic and heterotypic condensates have different material properties. We examined the individual P-body proteins that form homotypic condensates under cellular conditions (Fig. 6A). Dhh1 is clearly less dynamic in homotypic condensates compared to heterotypic condensates at 2 hours (Fig. 6B-C). Dcp1, Edc3, and Pat1 were slightly less dynamic in homotypic condensates, though the relative differences for these proteins are small because they also show low recovery in heterotypic condensates (Fig. 6C and S28). We conclude that proteins have reduced dynamics in homotypic condensates compared to heterotypic condensates, though the difference is subtle for Dcp1, Edc3, and Pat1.

### RNA delays the maturation and promotes the reversibility of *in vitro* P-bodies

We used additional approaches to further dissect differences in material states between heterotypic and homotypic condensates. First, we monitored incorporation of Thioflavin T (ThT) into condensates over time. ThT is a stain for amyloid fibers, and it has been observed in other systems that *in vitro* condensates increasingly stain with ThT as they mature (12, 15, 81). However, we only rarely observe ThT staining of the complete condensates containing all proteins and RNA, even at longer incubations (24 hr: Fig. 6D-E). Conversely, the vast majority of condensates with all P-body proteins, but without RNA, incorportated ThT at 24 hr (Fig. 6D-E). Thus, RNA prevents maturation of reconstituted P bodies to a state that binds ThT (likely preventing formation of amyloid fibers). Pat1, Edc3, and Dhh1 homotypic condensates behaved similarly at 24 hr; the individual protein condensates frequently stained positive for ThT in the absence of RNA, but much less frequently in the presence of RNA (Fig. 6E and S29). Dcp1 was the lone outlier, whose homotypic condensates stained with ThT with or without RNA, consistent with the protein lacking measurable binding to RNA (Fig. S10). With the caveat that some amyloid-like fibers may not bind ThT, these results suggest that P-body protein condensates begin to form amyloid fibers over time, and that the presence of RNA inhibits or slows this transition.

Amyloid formation can result in increased resistance to proteolysis; therefore, we next investigated whether the inclusion of RNA affected trypsin digestion of condensates. We formed condensates with all P-body proteins, either with or without RNA, and used Dhh1-GFP and Pat1- RFP as markers for the structures. We imaged the condensates over a sixty-minute time course after adding trypsin. Condensates in the presence of RNA were more rapidly and fully degraded by trypsin digestion (Fig. 6F-G and S30). These results further support the idea that *in vitro* P-bodies with RNA are less amyloid-like and more reversible.

As an additional measure of P-body dissolution we used arginine (500 mM) to disrupt condensates, with and without RNA, using Pat1-RFP as a marker. After forming the condensates for two hours, we added either arginine or an equivalent volume of buffer as a control. Condensates with RNA were dissolved by arginine, but not by the buffer control (Fig. 6H-I and S31). In contrast, arginine was less effective at dissolving condensates that lacked RNA. Condensates with Pat1 as the only protein behaved similarly: arginine dissolved Pat1 condensates with RNA, whereas Pat1 condensates without RNA were refractory to arginine dissolution (Fig. 6H-I and S31). These data also support the idea that RNA enhances the reversibility of condensates.

## Discussion

### Reconstitution of P bodies

In this study we have reconstituted RNA processing bodies using seven proteins that are highly concentrated in cellular P bodies (39), and RNA. When mixed at their cellular concentrations and under experimental conditions that mimic the cellular environment these molecules assemble together to form condensates that contain all seven proteins and RNA. Moreover, the condensates are quantitatively consistent with cellular P bodies in terms of protein partitioning and dynamics, and show a similar dependence on Pat1 (Fig. 5) (39). These results suggest that the interactions between the molecules, including binary binding and higher-order cooperativities, are sufficient to encode these cellular properties (Fig. 7). While many cellular inputs could modulate the behaviors of P bodies, including additional lower-expressed components, post-translational modifications, and chemical or mechanical energy, they are not necessary to produce a compartment that is native-like in its composition and dynamics. Below we discuss how our reconstitution compares to previous cellular and theoretical experiments, and we propose future directions for using the system to further probe multicomponent condensate formation and function.

**Figure 7.**
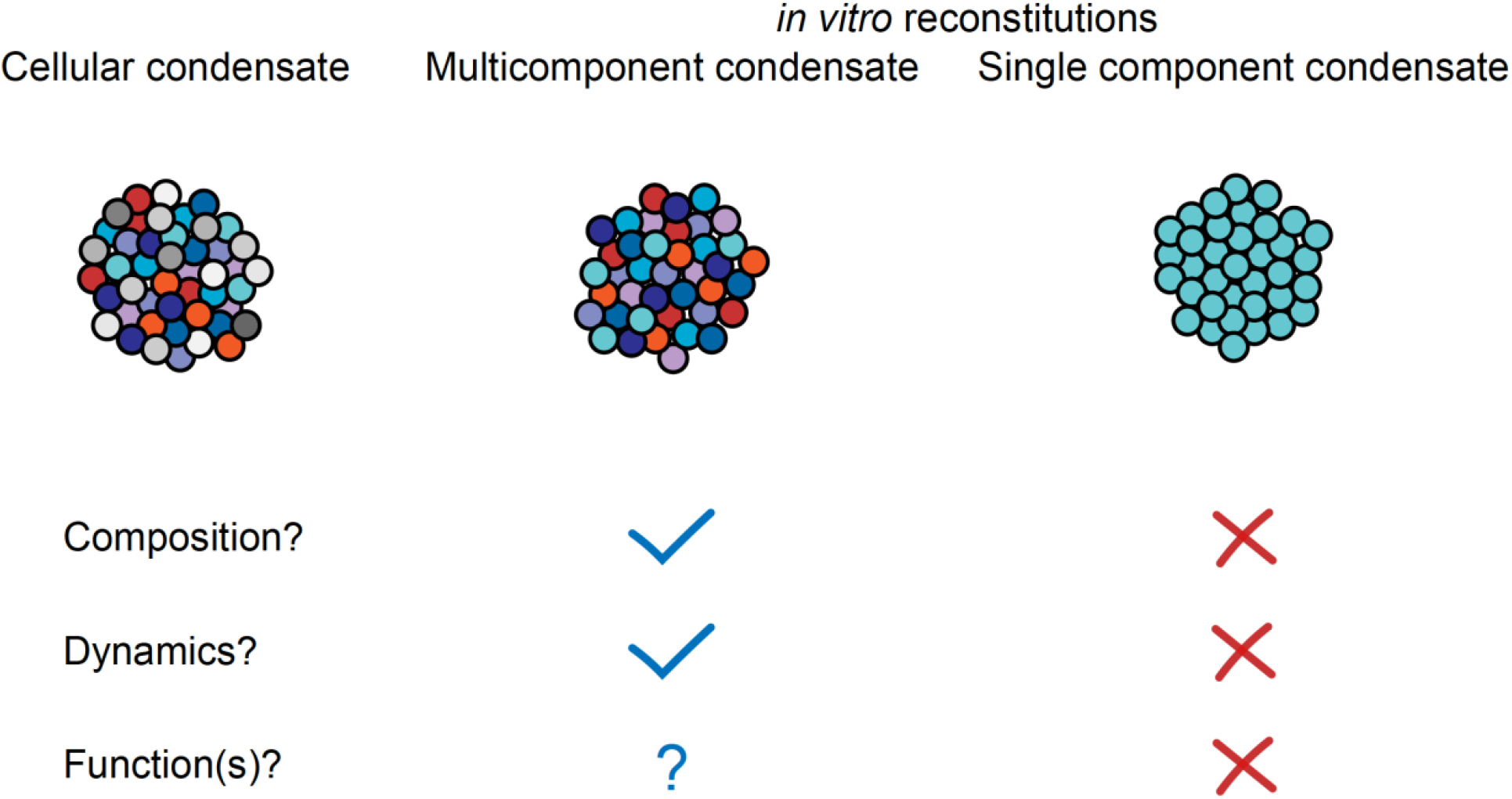
Multicomponent condensates recapitulate the composition and dynamics of highly-concentrated components. Cellular condensates consist of highly-concentrated molecules, represented by different colored circles, and less concentrated molecules, represented by different shades of gray. In this study we found that reconstituting P bodies *in vitro* with all of the highly-concentrated proteins resulted in condensates with partitioning and dynamics values of individual components in quantitative agreement with cellular P bodies (39). Reconstitutions using single components, or more generally using arbitrarily-reduced complexity, often still form condensates *in vitro*, but these condensates are distinct from cellular condensates in terms of their composition and/or material properties (62). While we have not yet assayed the function of our *in vitro* P bodies, we suggest that this level of reconstitution – including three enzymes and many regulators of these enzymes – will allow for the examination of multiple relevant biochemical activities, and the interplay between these activities (see Discussion).

### Integration of homotypic and heterotypic interactions for P-body assembly

Using conditions that mimic the cellular environment we are able to form condensates that contain all seven P-body proteins and RNA. However, under the same conditions, five proteins (Dhh1, Edc3, Xrn1, Edc3, and Pat1) are capable of forming homotypic condensates on their own, and these proteins tend to have higher partitioning in homotypic compared to the heterotypic condensates. These data suggest that there is a balance between homotypic and heterotypic interactions that contribute to P-body formation. Recent theoretical descriptions have combined a polyphasic linkage formalism (82) with coarse-grained linear polymer modeling to investigate which molecular features regulate the formation of multicomponent condensates (38, 79, 80). In this framework, interactions between stickers in multi-valent scaffold molecules drive condensate formation whereas spacers modulate condensate formation. Ligands/clients (low-valent, non-scaffold molecules that are enriched in condensates but do not promote their formation) can modulate condensate formation by binding to stickers or spacers, with distinct expectations for the change in molecule partitioning based on the nature of the ligand and its binding site on the scaffold. In our system, Dhh1, Edc3, Xrn1, Dcp1, and Pat1 can all self-associate, as each form homotypic condensates under cellular conditions (Fig. 6). Multiple stickers, consisting both of structured domains and IDRs, are present within these proteins (Fig. 4 and S1). Various subsets of P-body proteins have been shown to cooperatively phase separate, including: Dcp1, Dcp2 and Edc3 (21, 50); Dcp2, Pat1 and Lsm1-7 (52); and Dhh1, Pat1, and RNA (51). Interactions, between the different molecules (39, 54) drive the complete collection to form a single condensate including all molecule types. Based on the lower partitioning of many of the proteins in heterotypic condensates, there are likely some heterotypic interactions that disrupt sticker-sticker interactions. For example, Pat1 is an important scaffold-like protein for P-body formation (53, 56), and Pat1 interacts with itself and with all of the other highly-concentrated P body proteins and RNA (57, 83-85). In contrast, since Lsm1-7 only interacts with RNA and Pat1 (57, 86), Lsm1-7 binding to Pat1 may compete with other Pat1-mediated interactions, and thus limit the overall valency of the system. Cellular data potentially supports this limiting function of Lsm1-7 as genetic deletion of Lsm1 enhances P-body formation in yeast cells (53). However, this strain also exhibits a RNA-decapping defect and accumulates deadenylated mRNAs which may also contribute to P-body formation (87, 88). Other heterotypic interactions will likely have minimal effect on scaffold partitioning, either by binding to spacers within the scaffold molecules, or by forming a new set of interactions that is comparable in valency to those disrupted in the scaffold molecule. The Dhh1-Pat1 interaction (57, 84) appears to be in this latter class as deletion of Dhh1 has minimal impact on P-body formation and the partitioning of other P-body proteins in yeast cells (14, 53). A general and important point here is that the complexity of interactions among highly concentrated P-body components is not likely to be recapitulated in simplified systems examining only a few components at a time. Thus, formation of homotypic condensates will not always be relevant in the context of cellular multicomponent condensates where a combination of homotypic and heterotypic interactions impart thermodynamic and material properties to the system. Additional experiments are needed to further interrogate which heterotypic interactions strongly modulate P-body formation and composition.

Our results suggest that acidification of the cytoplasm may contribute to P-body formation. Prior *in vivo* studies indicated that cytoplasmic acidification restricted the mobility of mRNAs (63, 64). This reduced mobility may be due to the formation of P bodies, as we found that acidic pH promotes homotypic protein-protein interactions and heterotypic protein-RNA interactions through protonation of a key histidine residue on Dcp1, and likely multiple histidine residues dispersed throughout the other P-body proteins (Fig. 2 and 3). Importantly, acidic pH is also required to generate molecular stoichiometry in our reconstitution that is consistent with cellular P bodies measured in glucose starvation conditions, where the cytoplasm is acidic (Fig. 5) (39, 64). It is possible that the absence of macroscopic P-bodies during cellular proliferation may be due in part to the more basic cytoplasmic pH under these conditions, but there are likely additional cellular mechanisms that contribute to repressing P-body formation. However, no additional regulatory mechanisms are apparently required for the formation of P-bodies with appropriate molecular stoichiometry and material properties (Fig. 5 and 7).

### Reconstitution mimics the redundant nature of cellular assemblies

Our *in vitro* results are consistent with previous genetic experiments indicating that no single protein is absolutely required for P body formation *in vivo*, although some are more important than others (53-56). Our results are also consistent with previous reconstitutions that used a more limited set of P-body proteins, in that several distinct minimal collections of P-body proteins are able to form condensates *in vitro* (14, 21, 50-52). These results suggest that there are multiple, somewhat redundant, protein scaffolds that can act collectively to assemble many other P-body components (54). Excluding Pat1 restricts, but does not completely eliminate, the formation of condensates. Why does the loss of Pat1 dampen condensate formation when remaining individual proteins (Dhh1, Edc3, Xrn1, and Dcp1) robustly form homotypic condensates on their own? Our current interpretation of these results is that heterotypic interactions restrict the homotypic interactions of the remaining scaffold-like proteins when Pat1 is removed. This interpretation suggests that there is epistasis in P-body formation; that is, the contribution of any given molecule to condensate formation is dependent on which other molecules are present (9, 12, 37, 38, 58). If epistasis is a general feature of condensate formation, then reconstitutions that approach the molecular complexity of cellular condensates will more accurately reveal the contributions that each molecule makes to condensate formation. This reconstitution best mimics cellular P bodies when using physiological protein concentrations (Fig. 5). Using different absolute or relative concentrations of components, and/or a different collection of molecules, will likely influence the epistatic interactions between components. Thus, an important future direction will be to understand quantitatively how the constituent molecules influence the formation of P bodies (i.e. the threshold concentrations for appearance of the compartments) and their composition.

The role of RNA in formation and maintenance of P bodies is not fully understood. In biochemical reconstitutions, one report described a strong effect of RNA on the threshold concentrations of Dcp1/Dcp2/Edc3 for phase separation (50). Another, however, observed that RNA does not change the partitioning of Dcp1/Dcp2/Edc3 enrichment in phase separated droplets, except when Edc3 is in large (16-fold) excess over the other proteins, suggesting that RNA does not strongly promote phase separation (21). Here, we have similarly observed that although RNA is recruited into reconstituted P bodies in a length-dependent manner, it does not change the partitioning of proteins into the structures and is not required for their formation at cellular concentrations. Similar ambiguity has been reported in more physiological systems. In cell lysates, digestion of RNA causes dissolution of P bodies (59), and in cells, P bodies form in proportion to the pool of deadenylated mRNAs (45, 58). Yet, it was recently demonstrated that acute RNA degradation by RNAse L does not disrupt P bodies in intact human cells (60). One potential resolution to these apparent contradictions is that RNA may be important for the initiation of P bodies *in vivo*, but not for their maintenance. Thus, long timescale genetic perturbations may prevent P-body formation, while acute depletion of mRNA does not affect existing P bodies. Additional studies of the time dependence of RNA perturbations *in vivo* and *in vitro* may help resolve this issue.

Previous studies have shown that in different circumstances RNA can either prevent or enhance condensate formation by RNA-binding proteins (12, 61, 89-92). Here, we found that RNA is important for promoting the reversibility of P bodies and delaying their maturation into an irreversible, non-dynamic, amyloid-like state (Fig. 6). This function may be related to the ability of RNA to chaperone aggregation-prone proteins (93). Many of the regions of P-body proteins that interact with RNA also make homotypic interactions (12, 50, 55-57, 73). For example, the central IDR of Edc3 binds to RNA, interacts with the C-terminal YjeF-N domain, and promotes LLPS (50, 73). Thus, one mechanism by which RNA promotes a reversible state of P bodies may be by restricting homotypic protein interactions. Consistent with our findings, P bodies are relatively liquid-like in yeast and only progress to a solid state when individual proteins are artificially overexpressed (81). This suggests that the stoichiometry of resident protein and RNA molecules regulates the material properties of P bodies, and likely other condensates. In this way, stoichiometry could also regulate P-body functions, as was previously observed for reconstituted membrane-associated signaling condensates, where stoichiometry controlled membrane dwell time and consequently activity (20, 22).

### Potential implications of multiple scaffold-like molecules

One feature enabled by a multi-scaffold system is the colocalization and coordination of multiple enzymatic activities, as well regulators of those activities. In the case of P bodies, the highly concentrated P-body proteins possess RNA-helicase (Dhh1), RNA-decapping (Dcp2), and RNA- exonucleolytic degradation (Xrn1) activities within distinct polypeptides (40-43). Dcp1, Dhh1, Edc3, and Pat1 are known to stimulate the activity of Dcp2 (43, 57). Thus, forming a heterotypic condensate that includes all of these proteins could in theory serve as a concentrated microreactor to increase RNA degradation. Recent work demonstrated that Dcp2-mediated RNA decapping is enhanced within *in vitro* condensates containing Dcp1, Dcp2, and Edc3, but is inhibited in condensates that only had Dcp1 and Dcp2 (21). This example illustrates how composition can modulate enzymatic activities within condensates, and highlights the importance of efforts to mimic cellular complexity when investigating the biochemical activities of condensates. In addition to individual biochemical activities being increased within P bodies, the transfer of substrate RNA between different types of enzymes could also be enhanced, particularly if certain molecules are spatially organized within the condensates (94, 95), potentially resulting in a significant increase in the overall rate of RNA degradation. This reconstitution, which includes all highly enriched P-body proteins and many activators, is suitable for future examination of enzymatic activities and pathway flux within P bodies.

Combined with genetic experiments, our *in vivo* (39) and *in vitro* examinations of P bodies provide a route to systematically determine which components play important roles in forming condensates and specifying their physical properties. The ability to form homotypic condensates *in vitro* is often considered evidence that a molecule plays an important scaffolding role *in vivo*. However, given the prevalence of proteins that can undergo phase separation *in vitro* under different conditions (96-99), the redundant scaffolding observed in P-bodies may be a common feature of many cellular condensates. This view is consistent with recent network-based models for condensate formation, where multiple macromolecules can contribute to different degrees, based on their positions in the molecular interaction network and degree of connectivity to other components (8, 10, 39). In future biochemical studies it will be important to examine candidate scaffolds in experimental contexts that closely mimic cellular conditions in order to understand how multiple molecule types can be integrated to form multicomponent condensates. Using quantitative, multicomponent reconstitutions will enable a better understanding of the formation mechanisms, and ultimately biochemical and cellular functions, of diverse biomolecular condensates.

## Materials and Methods

### Lead Contact and Materials Availability

Requests for reagents should be directed to the Lead Contact, Michael K. Rosen. Plasmids generated for this study are available by contacting the Lead Author.

### Cloning of expression plasmids

Genes for Dcp1, Dcp2, Dhh1, Edc3, and Pat1 were cloned into a modified pMAL plasmid (New England Biolabs) using standard methods and NdeI and BamHI restriction sites. In order to use this strategy, silent mutations were first inserted into the coding sequences to eliminate restriction sites in Dcp2 (one NdeI site), Edc3 (one BamHI site), and Pat1 (two NdeI sites), using standard site directed mutagenesis. The modified pMAL plasmid (pMTTH) contains TEV-cleavable N-terminal MBP and C-terminal His_6_ tags. These genes were also cloned into a modified MTTH vector with monomeric GFP cloned in after the P-body gene and before the second TEV cleavage site/His_6_ tag.

Expression plasmids for Lsm1-7 and Xrn1 were generous gifts from Yigong Shi and Lionel Benard, respectively.

### Protein expression and purification

Lsm1-7 and Xrn1 were expressed and purified as described previously (40, 66, 67). The general protocol for expressing and purifying the other P-body proteins is described below, with protein-specific information provided thereafter. Plasmids were transfected into *Escherichia Coli* BL21 (DE3) cells and grown overnight at 37 °C on Luria Broth (LB)/Ampicillin agar plates. Individual colonies were resuspended in a 50 mL culture of LB/Ampicillin and grown overnight at 37 °C. Cells were collected by centrifugation (3,400 x *g*, 10 minutes), resuspended in LB, and added to six, one-liter cultures of LB/Ampicillin in four-liter unbaffled flasks. Cultures were grown at 37 °C until an OD_600_ of ∼ 0.5 at which point the temperature was decreased to 18 °C. Cultures were induced with 1 mM IPTG at an OD_600_ of ∼ 1.0 and grown overnight at 18 °C. Cells were collected by centrifugation (4,700 x *g*, 40 minutes) and the pellet was resuspended in 25 mM Tris pH 8, 10% (V:V) glycerol, 500 mM NaCl, 10 mM imidazole, and 5 mM *β*-mercaptoethanol (BME). Suspensions were transferred to 50 mL conical tubes and stored at -80 °C for future use.

Cell suspensions were thawed in cold water and lysed using a cell homogenizer (10,000 psi, 3 passes on ice). Lysate was centrifuged (45,000 x *g*, 30 minutes) and the cleared supernatant was added to ∼ 15 mL of Ni^2+^ agarose resin (Biorad), and incubated for 30 minutes with circular rotation in 50 mL conical tubes. Proteins were isolated using gravity columns and the following buffers. Ni^2+^ wash 1: 25 mM Tris pH 8, 10% glycerol, 2.5 M NaCl, 10 mM imidazole, 5 mM BME; Ni^2+^ wash 2: 25 mM Tris pH 8, 10% glycerol, 500 mM NaCl, 10 mM imidazole, 5 mM BME; Ni^2+^ elution: 25 mM Tris pH 8, 10% glycerol, 500 mM NaCl, 500 mM imidazole, 5 mM BME. The eluate was added to ∼ 30 mL of amylose resin (New England Biolabs) and incubated for 30 minutes with circular rotation in 50 mL conical tubes. Proteins were isolated using gravity columns and the following buffers. Amylose wash 1: 25 mM Tris pH 8, 10% glycerol, 150 mM NaCl, 5 mM BME; amylose elution: 25 mM Tris pH 8, 10% glycerol, 150 mM NaCl, 5 mM BME, and 50 mM maltose. The amylose eluate was diluted threefold into 25 mM Tris pH 8, 0 mM NaCl, and 5 mM BME for a final concentration of 50 mM NaCl, and filtered through a 0.45 µm Whatman filter (GE Healthcare). Filtrate was loaded onto a Q sepharose ion-exchange column and eluted using a 50 – 500 mM NaCl gradient over ten column volumes, taking fractions every 2.5 mL. Fractions containing the desired protein were were loaded onto an SD200 26/600 size exclusion column equilibrated with 10 mM MES pH 7, 5% glycerol, 300 mM KOAc, and 5 mM BME. Fractions containing the desired protein were concentrated by ultrafiltration (Amicon centricon) with 3k (Lsm1-7) and 30k (all other proteins) molecular weight cutoffs. Single-use aliquots were flash frozen in liquid nitrogen and stored at -80 °C for future use.

Cleaving the MBP-fusion tag before ion-exchange and size-exclusion columns resulted in a partial loss of protein for Dhh1 and a total loss of protein for Edc3 and Pat1. Thus, to simplify procedures and maintain consistency, the MBP-tag was retained on all proteins during purification, and condensate formation was initiated by TEV cleavage.

Protein-specific deviations from the general protocol are as follows. Only two liters of culture were used for Dcp1 and Dhh1. Dcp2 was expressed in terrific buffer (TB) instead of LB due to low expression of the full-length protein. 10 mM ATP was included in the wash buffers for the Ni^2+^ column for Dhh1 in order to disrupt interactions with RNA. Without ATP, Dhh1 eluted as a series of peaks from the ion-exchange column presumably due to an inhomogeneous distribution of bound RNA molecules. 1M urea was included in the ion-exchange buffers for Pat1 to prevent aberrant oligomerization and/or precipitation. The size exclusion buffer for Pat1 included 1 M KOAc to similarly eliminate aberrant oligomerization and/or precipitation.

See Table S5 for protein sequences.

### RNA reagents

RPL41A and RNA10 RNA (Fig. S11) were purchased from Integrated DNA Technologies, resuspended in 10 mM Tris-HCl, 1 mM EDTA, pH 7.0 buffer, and single-use aliquots were flash frozen in liquid nitrogen and stored at -80 °C for future use. Total yeast RNA was purchased from Sigma-Aldrich and solubilized and stored as described above. MFA2 RNA was generated using *in vitro* transcription. Template DNA was generated by PCR out of the pRP802 plasmid (Parker lab) using the following oligos: TAATACGACTCACTATAGCGAGC and TTTTTCATGAAAAAATCTGTTAAAGTGATAACTAC. MFA2 RNA was transcribed from this DNA template using the Invitrogen Ambion MEGAscript T7 Transcription Kit by following the standard protocol with 15% Cy5-UTP included. The reaction was allowed to proceed overnight, then treated with the Megaclear Transcription Clean-Up Kit (Thermo Scientific). A sample of RNA was run on a 5% polyacrylamide/8 M urea gel to confirm correct size and specificity. Extent of Cy5 labeling was determined using a NanoDrop spectrophotometer. MFA2 RNA samples were flash frozen in liquid nitrogen and stored at -80 °C for future use.

### Protein structure and disorder predictions

Structured and disordered regions were determined based on the following criteria (Fig. 1A and S1). Known three-dimensional structures were curated from the literature (73, 83-85, 100-102). Secondary structure and protein disorder were predicted using PSIPRED and Disprot VSL2, respectively (103, 104). Pat1 middle domain (MD: Pat1^241-422^) was the only region in these P-body proteins that had predicted secondary structure without a previously determined three-dimensional structure (Fig. S2 and S3). We predicted the structure of Pat1^241-422^ using AlphaFold (Fig. S2F)(105, 106). This model confidently predicts multiple *α*-helices within Pat1^241-422^, consistent with PSIPRED secondary structure prediction (data not shown) and disorder prediction (Fig. S3). However, we note that while the AlphaFold model confidently predicts the presence of *α*-helices within Pat1^241-422^, their orientation relative to one another and relative to the rest of Pat1 is unclear (see Fig. S2 for further details). The Dcp1 model (Fig. 2C) was made by overlapping structures for the Dcp1 dimer (72) with Dcp1-Xrn1 (100) and Dcp1-Dcp2 (101) interactions using the main chain of Dcp1 for alignment. The Dcp2 homology model (Fig. S9A) was generated by mutating *S. cerevisiae*-specific residues onto the *Schizosaccharomyces pombe* structure (75). All protein structures were visualized using PyMol.

### Microscopy

Microscopy experiments were carried out in 384-well glass bottom microwell plates (Brooks Life Science Systems: MGB101-1-2-LG-L). Prior to use the plates were cleaned with 5% Hellmanex III, then 1M NaOH, and then passivated with mPEG silane, with extensive milliQ water washes between each step. On the day of use individual wells were blocked with 2% bovine serum albumin, then rinsed thoroughly with reaction buffer. Single-use aliquots of proteins were quickly thawed in cold water, centrifuged at 16,000 x *g* for 10 min at 4 °C, then stored on ice. Protein concentrations were quantified using UV-Vis spectroscopy (absorbance at 280 nm) and rechecked before each experiment. Single-use RNA aliquots were thawed on ice and heated to 70 °C for 90 s immediately before use. For complex reactions, the components were added to the well in the following order to promote heterotypic condensate formation and minimize homotypic condensate formation: buffer, RNA, Dcp2, Lsm1-7, Xrn1, Dhh1, Pat1, Edc3, and Dcp1. Pipetting into plate wells was carried out at room temperature as quickly as possible, then plates were incubated at 30 °C for 2 hours. TEV was added at a 1:50 molar ratio, relative to total protein and RNA concentrations, to initiate condensate formation. We note that different procedures, e.g. involving TEV cleavage before mixing and/or insufficient preincubation of P-body components, yielded heterogeneous condensates and poor reproducibility. Reactions were incubated and imaged at 30 °C unless stated otherwise.

Condensate images were captured on a Nikon Eclipse Ti microscope base with a Yokogawa CSU-X1 spinning disk confocal scanner unit, 100 × 1.49 NA objective, and Andor EM- CCD camera. Fluorescence Recovery After Photobleaching (FRAP) was performed with a TIRF/iLAS2 FRAP Module (Biovision) and Rapp UGA-40 Phototargeter. Z-stacks were collected with the exception of FRAP experiments as the rapid collection of subsequent timepoints did not allow for the collection of a Z-stack at each time point.

Micrographs were analyzed using FIJI (107). For Z-stacks the Z plane with the highest intensity was selected for analysis. A threshold was set at three-fold above the background signal for each channel. Concentrations were determined from fluorescence intensities using standard curves for RFP (Pat1), Alexa Fluor 488 (Lsm1-7 and Xrn1), and GFP (Dcp1, Dcp2, Dhh1, Edc3, and Pat1) (Fig. S32-34). Small condensates, that are close in size (xy area) to the point spread function (PSF) of a microscope exhibit diluted intensities (39). We empirically determined that the intensity of condensates smaller than 2.5 µm in diameter exhibit a linear dependence on condensate size, whereas the intensities of condensates larger than 2.5 µm in diameter were independent of condensate size (Fig. S35). Thus condensates larger than 2.5 µm in diameter were selected for further analysis of condensate concentration. For calculating fraction overlap, condensates with centers of mass within 1 µm of each other in the different channels were considered to be overlapping.

50 µM ThT was added to samples to image ThT staining of condensates. For imaging condensates with all P-body proteins, with or without RNA, and for Pat1 homotypic condensates, Pat1-RFP was used as a marker for the condensates. Condensates were considered to stain positive for ThT if the centers of mass from the RFP and ThT channels were within 1 µm of each other (Fig. 6D and S29A). ThT staining cannot be used when GFP is the marker for a condensate due to the overlap between GFP and ThT fluorescence spectra. Instead, for Dcp1, Dhh1, and Edc3 homotypic condensates we estimated the percentage of ThT positive condensates as follows. We performed parallel experiments with either GFP-tagged proteins only to count the number of condensates, or with unlabelled proteins and ThT to count the number of ThT positive condensates. We estimated the percentage of ThT positive condensates by dividing the number of ThT positive condensates by the number of total condensates (Fig. S29B).

For the proteolysis experiments 0.8 µg trypsin was added to each reaction. The condensed phase concentration for Pat1-RFP and Dhh1-GFP were monitored before and after trypsin addition. Relative condensed phase protein concentrations based on pre-trypsin levels are displayed (Fig. 6G and S30).

### Electrophoretic Mobility Shift Assays

Reactions included 10 nM Alexa Fluor647-labeled RPL41A RNA. P-body proteins were titrated over >100-fold range, with exact values depending on expression/purification yield and affinity for RNA (Fig. S10), in 300 mM KOAc, 10 mM MES pH 7 or pH 5.8, and 5 mM BME. Reactions were incubated on ice for 2 hr, then resolved on a 6% native PAGE gel. 0.5x MBE buffer (MES, pH 7 or 5.8; Borate; EDTA) and gels were equilibrated and run at 4 °C. Gels were imaged using Bio-Rad ChemiDoc MP. Bands were detected using Bio-Rad Image Lab software (v. 6.1) and curves were fit using the Specific Binding with Hill Slope equation on Graphpad Prism (v. 9.0.0 for Mac).

### Native Gels

Reactions with Dcp1 were incubated at room temperature at the indicated pH (5.8 or 7) without TEV cleavage for 2 hours. 2x sample buffer (62.5 mM MES pH 7, 25% glycerol, 1% bromophenol blue) was then added and samples were loaded onto a native polyacrylamide (37.5:1 acrylamide:bis-acrylamide) gel with 4% stacking and 6% separating portions. Gels used 25 mM MES pH 7 as the buffer. Running buffer with 25 mM MES pH 7 and 192 mM Glycine was used and gels were pre run for 30 minutes before loading samples.

### Quantification and Statistical Analyses

Unless stated otherwise, data points and error bars in figures correspond to the mean and standard deviation (SD) from each of six total micrographs from two replicate experiments. All statistical tests were performed using Microsoft Excel. A two-tailed paired t test was performed on the data in Fig. 2E, 2G, 6A, 6I, S23, and S31. A one-tailed paired t test was performed on Fig. 6E, and S18H. A two-tailed heteroscedastic t test was performed on the data in Fig. 3E. *, **, and *** symbols correspond to *p* values less than 0.05, 0.01, and 0.001, respectively.

## Acknowledgements

We thank Yigong Shi and Lionel Bénard for expression plasmids of Lsm1-7 and Xrn1, respectively. We thank past and present members of the Rosen lab and Furqan Dar, Kiersten Ruff, and Rohit Pappu for technical assistance and helpful discussions regarding this project. Research was supported by the Howard Hughes Medical Institute (to R.P. and M.K.R) and grants from the Welch Foundation (I-1544 to M.K.R.).

## Author Contributions

S.L.C. and M.K.R conceived the study and S.L.C., W.X., R.P., and M.K.R. designed the research program. S.L.C. performed the cloning, protein expression and purification, microscopy and biochemical experiments, and S.L.C., W.X., R.P., and M.K.R. analyzed the data. W.X., D.M., and C.J.D. provided critical reagents and intellectual contributions. R.P. and M.K.R. secured funding and supervised the work. S.L.C., R.P., and M.K.R. wrote the manuscript and all authors reviewed and edited the manuscript.

## Declaration of Interests

R.P. and M.K.R. are cofounders and consultants for Faze Medicines.

