## Supplemental Information for "Quantitative reconstitution of yeast RNA processing bodies"

This PDF file includes:

Tables S1 to S5

Figures S1 to S35

Supplemental References
**Table S1.**

**Effect of gene deletion on P-body formation**

| Gene | Effect on other proteins in P bodies^#^ | |
| --- | --- | --- |
| deletion | Increased | Decreased |
| pat1Δ | - | Dcp1, Dcp2, Dhh1, Edc3, Lsm1, Xrn1 |
| edc3Δ | - | Dcp1, Dcp2, Dhh1, Lsm1, Pat1, Xrn1 |
| dhh1Δ | - | Dcp2 |
| dcp2Δ^$^ | Dhh1, Edc3, Lsm1, Pat1, Xrn1 | Dcp1 |
| lsm1Δ | Dcp1, Dcp2, Dhh1, Edc3, Xrn1 | Pat1^%^ |
| xrn1Δ | Dcp1, Dcp2, Dhh1, Edc3, Lsm1, Pat1 | - |
| dcp1Δ | Dcp2, Dhh1, Edc3, Lsm1, Pat1, Xrn1 | - |

^#^ References: (1-3)

^$^ Dcp2 deletion increases P-body formation during mid-log phase but decreases P-body formation under stress conditions, suggesting it has competing scaffolding and RNA-degradation activities that are integrated to either form or dissolve P bodies.

^%^ Lsm1 deletion reduces Pat1 accumulation in P bodies by enhancing nuclear retention of Pat1.

**Table S2.**

**Curve fitting for individual P-body proteins binding to RNA (Figs. 3B and S10)**

| Protein | pH | *K_D_* (μM) | *K_D_* (pH 7) / *K_D_* (pH 5.8) | Hill Coefficient^a^ | B_max_^b^ | n |
| --- | --- | --- | --- | --- | --- | --- |
| Dcp1 | 7 | > 20 | - | - | - | 3 |
|  | 5.8 | > 20 |  | - | - | 3 |
| Dcp2 | 7 | > 20 | > 16 | - | - | 3 |
|  | 5.8 | 1.3 ± 0.1 |  | 1.7 ± 0.4 | 1.0 ± 0.1 | 3 |
| Dhh1 | 7 | 0.7 ± 0.1 | 2.6 ± 0.3 | 2.2 ± 0.1 | 1.0 ± 0.1 | 3 |
|  | 5.8 | 0.3 ± 0.1 |  | 2.7 ± 0.6 | 1.0 ± 0.1 | 3 |
| Edc3 | 7 | 2.1 ± 0.8 | 3 ± 1 | 1.7 ± 0.1 | 1.3 ± 0.3 | 3 |
|  | 5.8 | 0.7 ± 0.2 |  | 2.9 ± 0.9 | 1.0 ± 0.1 | 3 |
| Lsm1-7 | 7 | 0.5 ± 0.1 | 1.7 ± 0.6 | 3 ± 1 | 1.0 ± 0.1 | 3 |
|  | 5.8 | 0.3 ± 0.1 |  | 4.2 ± 0.6 | 1.0 ± 0.1 | 3 |
| Pat1 | 7 | 0.7 ± 0.2 | 1.8 ± 0.6 | 1.5 ± 0.8 | 1.0 ± 0.1 | 3 |
|  | 5.8 | 0.4 ± 0.1 |  | 2.5 ± 0.7 | 1.0 ± 0.1 | 3 |

^a^ Hill coefficient was not determined (-) for conditions that did not reach saturation.

^b^ B_max_ was not determined (-) for conditions that did not reach saturation.

**Table S3.**

**Curve fitting for Dcp2 truncations and point mutants binding to RNA (Fig. 3E).**

| Protein | pH | *K_D_* (μM) | Fold difference^a^ | Hill coefficient^b^ | B_max_^c^ |
| --- | --- | --- | --- | --- | --- |
| Dcp2^1-300^ | 7 | >20 | - | **-** | **-** |
|  | 5.8 | 1.7 ± 0.3 | 1.3 ± 0.3 | 2.2 ± 0.6 | 1.0 ± 0.1 |
| Dcp2^301-970^ | 7 | > 20 | - | - | - |
|  | 5.8 | 1.9 ± 0.5 | 1.5 ± 0.4 | 1.3 ± 0.4 | 1.0 ± 0.1 |
| Dcp2^H237A^ | 7 | >20 | - | **-** | **-** |
|  | 5.8 | 2.8 ± 0.8 | 2.2 ± 0.6 | 3 | 0.9 ± 0.1 |

^a^ Fold difference relative to Dcp2 (Table S2).

^b^ Hill coefficient was not determined (-) for conditions that did not reach saturation.

^c^ B_max_ was not determined (-) for conditions that did not reach saturation.

**Table S4.**

**Partition coefficients for P-body proteins in heterotypic condensates (Fig 5E).**

| Protein | Partition Coefficient | Fold difference^a^ | n |
| --- | --- | --- | --- |
| Dcp1 | 11 ± 7 | 0.10 ± 0.07 | 8 |
| Dcp2 | 130 ± 20 | 1.0 ± 0.2 | 8 |
| Dhh1 | 31 ± 5 | 1.0 ± 0.2 | 8 |
| Edc3 | 120 ± 20 | 0.9 ± 0.2 | 8 |
| Lsm1-7 | 40 ± 20 | 0.8 ± 0.3 | 8 |
| Pat1 | 180 ± 30 | 1.7 ± 0.4 | 6 |
| Xrn1 | 80 ± 30 | 1.5 ± 0.6 | 4 |

^a^ Fold difference relative to *in vivo* values (4).

**Table S5.**

**Detailed information on recombinant proteins used in this study.**

| Protein | Sequence |
| --- | --- |
| Dcp1 | MTGAATAAENSATQLEFYRKALNFNVIGRYDPKIKQLLFHTPHASLYKWDFKKDEWNKLEYQGVLAIYLRDVSQNTNLLPVSPQEVDIFDSQNGSNNIQVNSGSDNSNRNSSGNGNSYKSNDSLTYNCGKTLSGKDIYNYGLIILNRINPDNFSMGIVPNSVVNKRKVFNAEEDTLNPLECVGVEVKDELVIIKNLKHEVYGIWIHTVSDRQNIYELIKYLLEYEPRDSCA |
| Dcp1^H40A^ | MTGAATAAENSATQLEFYRKALNFNVIGRYDPKIKQLLFATPHASLYKWDFKKDEWNKLEYQGVLAIYLRDVSQNTNLLPVSPQEVDIFDSQNGSNNIQVNSGSDNSNRNSSGNGNSYKSNDSLTYNCGKTLSGKDIYNYGLIILNRINPDNFSMGIVPNSVVNKRKVFNAEEDTLNPLECVGVEVKDELVIIKNLKHEVYGIWIHTVSDRQNIYELIKYLLEYEPRDSCA |
| Dcp1^H43A^ | MTGAATAAENSATQLEFYRKALNFNVIGRYDPKIKQLLFHTPAASLYKWDFKKDEWNKLEYQGVLAIYLRDVSQNTNLLPVSPQEVDIFDSQNGSNNIQVNSGSDNSNRNSSGNGNSYKSNDSLTYNCGKTLSGKDIYNYGLIILNRINPDNFSMGIVPNSVVNKRKVFNAEEDTLNPLECVGVEVKDELVIIKNLKHEVYGIWIHTVSDRQNIYELIKYLLEYEPRDSCA |
| Dcp1^H198A^ | MTGAATAAENSATQLEFYRKALNFNVIGRYDPKIKQLLFHTPHASLYKWDFKKDEWNKLEYQGVLAIYLRDVSQNTNLLPVSPQEVDIFDSQNGSNNIQVNSGSDNSNRNSSGNGNSYKSNDSLTYNCGKTLSGKDIYNYGLIILNRINPDNFSMGIVPNSVVNKRKVFNAEEDTLNPLECVGVEVKDELVIIKNLKAEVYGIWIHTVSDRQNIYELIKYLLEYEPRDSCA |
| Dcp1^H206A^ | MTGAATAAENSATQLEFYRKALNFNVIGRYDPKIKQLLFHTPHASLYKWDFKKDEWNKLEYQGVLAIYLRDVSQNTNLLPVSPQEVDIFDSQNGSNNIQVNSGSDNSNRNSSGNGNSYKSNDSLTYNCGKTLSGKDIYNYGLIILNRINPDNFSMGIVPNSVVNKRKVFNAEEDTLNPLECVGVEVKDELVIIKNLKHEVYGIWIATVSDRQNIYELIKYLLEYEPRDSCA |
| Dcp1^Δ82-129^ | MTGAATAAENSATQLEFYRKALNFNVIGRYDPKIKQLLFHTPHASLYKWDFKKDEWNKLEYQGVLAIYLRDVSQNTNLLPVKTLSGKDIYNYGLIILNRINPDNFSMGIVPNSVVNKRKVFNAEEDTLNPLECVGVEVKDELVIIKNLKHEVYGIWIHTVSDRQNIYELIKYLLEYEPRDSCA |
| Dcp1^82-129^ | SPQEVDIFDSQNGSNNIQVNNGSDNSNRNSSGNGNSYKSNDSLTYNCG |
| Dcp2 | MSLPLRHALENVTSVDRILEDLLVRFIINCPNEDLSSVERELFHFEEASWFYTDFIKLMNPTLPSLKIKSFAQLIIKLCPLVWKWDIRVDEALQQFSKYKKSIPVRGAAIFNENLSKILLVQGTESDSWSFPRGKISKDENDIDCCIREVKEEIGFDLTDYIDDNQFIERNIQGKNYKIFLISGVSEVFNFKPQVRNEIDKIEWFDFKKISKTMYKSNIKYYLINSMMRPLSMWLRHQRQIKNEDQLKSYAEEQLKLLLGITKEEQIDPGRELLNMLHTAVQANSNNNAVSNGQVPSSQELQHLKEHSGEHNQQKDQQSSFSSQQQPSIFPSLSEPFANNKNVIPPTMPMANVFMSNPQLFATMNGQPFAPFPFMLPLTNNSNSANPIPTPVPPNFNAPPNPMAFGVPNMHNLSGPAVSQPFSLPPAPLPRDSGYSSSSPGQLLDILNSKKPDSNVQSSKKPKLKILQRGTDLNSIKQNNNDETAHSNSQALLDLLKKPTSSQKIHASKPDTSFLPNDSVSGIQDAEYEDFESSSDEEVETARDERNSLNVDIGVNVMPSEKDSRRSQKEKPRNDASKTNLNASAESNSVEWGPGKSSPSTQSKQNSSVGMQNKYRQEIHIGDSDAYEVFESSSDEEDGKKLEELEQTQDNSKLISQDILKENNFQDGEVPHRDMPTESNKSINETVGLSSTTNTVKKVPKVKILKRGETFASLANDKKAFDSSSNVSSSKDLLQMLRNPISSTVSSNQQSPKSQHLSGDEEIMMMLKRNSVSKPQNSEENASTSSINDANASELLGMLKQKEKDITAPKQPYNVDSYSQKNSAKGLLNILKKNDSTGYPRTEGGPSSEMSTSMKRNDATNNQELDKNSTELLNYLKPKPLNDGYENISNKDSSHELLNILHGNKNSSAFNNNVYATDGYSLASDNNENSSNKLLNMLQNRSSAINEPNFDVRSNGTSGSNELLSILHRK |
| Dcp2^1-300^ | MSLPLRHALENVTSVDRILEDLLVRFIINCPNEDLSSVERELFHFEEASWFYTDFIKLMNPTLPSLKIKSFAQLIIKLCPLVWKWDIRVDEALQQFSKYKKSIPVRGAAIFNENLSKILLVQGTESDSWSFPRGKISKDENDIDCCIREVKEEIGFDLTDYIDDNQFIERNIQGKNYKIFLISGVSEVFNFKPQVRNEIDKIEWFDFKKISKTMYKSNIKYYLINSMMRPLSMWLRHQRQIKNEDQLKSYAEEQLKLLLGITKEEQIDPGRELLNMLHTAVQANSNNNAVSNGQVPSSQE |
| Dcp2^301-970^ | LQHLKEQSGEHNQQKDQQSSFSSQQQPSIFPSLSEPFANNKNVIPPTMPMANVFMSNPQL  FATMNGQPFAPFPFMLPLTNNSNSANPIPTPVPPNFNAPPNPMAFGVPNMHNLSGPAVS  QPFSLPPAPLPRDSGYSSSSPGQLLDILNSKKPDSNVQSSKKPKLKILQRGTDLNSIKQNNND  ETAHSNSQALLDLLKKPTSSQKIHASKPDTSFLPNDSVSGIQDAEYEDFESSSDEEVETARDER  NSLNVDIGVNVMPSEKDSRRSQKEKPRNDASKTNLNASAESNSVEWGPGKSSPSTQSKQN  SSVGMQNKYRQEIHIGDSDAYEVFESSSDEEDGKKLEELEQTQDNSKLISQDILKENNFQDGE  VPHRDMPTESNKSINETVGLSSTTNTVKKVPKVKILKRGETFASLANDKKAFDSSSNVSSSKDL  LQMLRNPISSTVSSNQQSPKSQHLSGDEEIMMMLKRNSVSKPQNSEENASTSSINDANASEL  LGMLKQKEKDITAPKQPYNVDSYSQKNSAKGLLNILKKNDSTGYPRTEGGPSSEMSTSMKRN  DATNNQELDKNSTELLNYLKPKPLNDGYENISNKDSSHELLNILHGNKNSSAFNNNVYATDGYSLASDNNENSSNKLLNMLQNRSSAINEPNFDVRSNGTSGSNELLSILHRK |
| Dcp2^H237A^ | MSLPLRHALENVTSVDRILEDLLVRFIINCPNEDLSSVERELFHFEEASWFYTDFIKLMNPTLPSLKIKSFAQLIIKLCPLVWKWDIRVDEALQQFSKYKKSIPVRGAAIFNENLSKILLVQGTESDSWSFPRGKISKDENDIDCCIREVKEEIGFDLTDYIDDNQFIERNIQGKNYKIFLISGVSEVFNFKPQVRNEIDKIEWFDFKKISKTMYKSNIKYYLINSMMRPLSMWLRAQRQIKNEDQLKSYAEEQLKLLLGITKEEQIDPGRELLNMLHTAVQANSNNNAVSNGQVPSSQELQHLKEQSGEHNQQKDQQSSFSSQQQPSIFPSLSEPFANNKNVIPPTMPMANVFMSNPQLFATMNGQPFAPFPFMLPLTNNSNSANPIPTPVPPNFNAPPNPMAFGVPNMHNLSGPAVSQPFSLPPAPLPRDSGYSSSSPGQLLDILNSKKPDSNVQSSKKPKLKILQRGTDLNSIKQNNNDETAHSNSQALLDLLKKPTSSQKIHASKPDTSFLPNDSVSGIQDAEYEDFESSSDEEVETARDERNSLNVDIGVNVMPSEKDSRRSQKEKPRNDASKTNLNASAESNSVEWGPGKSSPSTQSKQNSSVGMQNKYRQEIHIGDSDAYEVFESSSDEEDGKKLEELEQTQDNSKLISQDILKENNFQDGEVPHRDMPTESNKSINETVGLSSTTNTVKKVPKVKILKRGETFASLANDKKAFDSSSNVSSSKDLLQMLRNPISSTVSSNQQSPKSQHLSGDEEIMMMLKRNSVSKPQNSEENASTSSINDANASELLGMLKQKEKDITAPKQPYNVDSYSQKNSAKGLLNILKKNDSTGYPRTEGGPSSEMSTSMKRNDATNNQELDKNSTELLNYLKPKPLNDGYENISNKDSSHELLNILHGNKNSSAFNNNVYATDGYSLASDNNENSSNKLLNMLQNRSSAINEPNFDVRSNGTSGSNELLSILHRK |
| Dcp2^H237R^ | MSLPLRHALENVTSVDRILEDLLVRFIINCPNEDLSSVERELFHFEEASWFYTDFIKLMNPTLPSLKIKSFAQLIIKLCPLVWKWDIRVDEALQQFSKYKKSIPVRGAAIFNENLSKILLVQGTESDSWSFPRGKISKDENDIDCCIREVKEEIGFDLTDYIDDNQFIERNIQGKNYKIFLISGVSEVFNFKPQVRNEIDKIEWFDFKKISKTMYKSNIKYYLINSMMRPLSMWLRRQRQIKNEDQLKSYAEEQLKLLLGITKEEQIDPGRELLNMLHTAVQANSNNNAVSNGQVPSSQELQHLKEQSGEHNQQKDQQSSFSSQQQPSIFPSLSEPFANNKNVIPPTMPMANVFMSNPQLFATMNGQPFAPFPFMLPLTNNSNSANPIPTPVPPNFNAPPNPMAFGVPNMHNLSGPAVSQPFSLPPAPLPRDSGYSSSSPGQLLDILNSKKPDSNVQSSKKPKLKILQRGTDLNSIKQNNNDETAHSNSQALLDLLKKPTSSQKIHASKPDTSFLPNDSVSGIQDAEYEDFESSSDEEVETARDERNSLNVDIGVNVMPSEKDSRRSQKEKPRNDASKTNLNASAESNSVEWGPGKSSPSTQSKQNSSVGMQNKYRQEIHIGDSDAYEVFESSSDEEDGKKLEELEQTQDNSKLISQDILKENNFQDGEVPHRDMPTESNKSINETVGLSSTTNTVKKVPKVKILKRGETFASLANDKKAFDSSSNVSSSKDLLQMLRNPISSTVSSNQQSPKSQHLSGDEEIMMMLKRNSVSKPQNSEENASTSSINDANASELLGMLKQKEKDITAPKQPYNVDSYSQKNSAKGLLNILKKNDSTGYPRTEGGPSSEMSTSMKRNDATNNQELDKNSTELLNYLKPKPLNDGYENISNKDSSHELLNILHGNKNSSAFNNNVYATDGYSLASDNNENSSNKLLNMLQNRSSAINEPNFDVRSNGTSGSNELLSILHRK |
| Dhh1 | MGSINNNFNTNNNSNTDLDRDWKTALNIPKKDTRPQTDDVLNTRGNTFEDFYLKRELLMGIFEAGFEKPSPIQEEAIPVAITGRDILARAKNGTGKTAAFVIPTLEKVKPKLNKIRALIMVPTRELALQTSQVVRTLGKHCGISCMVTTGGTNLRDDILRLNETVHILVGTPGRVLDLASRRVADLSDCSLFIMDEADKMLSRDFKTIIEQILSFLPPTHQSLLFSATFPLTVKEFVVKHLHKPYDINVMEELTLKGITQYYAFVEERQELHCLNTLFSKLQINQAIIFCNSTNRVELLAKKITDLGYSCYYSHARMKQQERNKVFHEFRQGKVRTLVCSDLLTRGIDIQAVNVVINFDFPKTAETYLHRIGRSGRFGHLGLAINLINWNDRFNLYKIEQELGTEIAAIPATIDKSLYVAENDETVPVPFPIEQQSYHQQAIPQQQLPSQQQFAIPPQQHHPQFMVPPSHQQQQAYPPPQMPSQQGYPPQQEHFMAMPPGQSQPQY |
| Dhh1^25-425^ | ALNIPKKDTRPQTDDVLNTKGNTFEDFYLKRELLMGIFEAGFEKPSPIQEEAIPVAITGRDILARAKNGTGKTAAFVIPTLEKVKPKLNKIRALIMVPTRELALQTSQVVRTLGKHCGISCMVTTGGTNLRDDILRLNETVHILVGTPGRVLDLASRKVADLSDCSLFIMDEADKMLSRDFKTIIEQILSFLPPTHQSLLFSATFPLTVKEFVVKHLHKPYDINLMEELTLKGITQYYAFVEERQKLHCLNTLFSKLQINQAIIFCNSTNRVELLAKKITDLGYSCYYSHARMKQQERNKVFHEFRQGKVRTLVCSDLLTRGIDIQAVNVVINFDFPKTAETYLHRIGRSGRFGHLGLAINLINWNDRFNLYKIEQELGTEIAAIPATIDKSLYVAENDET |
| Dhh1^426-506^ | VPVPFPIEQQSYHQQAIPQQQLPSQQQFAIPPQQHHPQFMVPPSHQQQQAYPPPQMPSQQGY |
| Edc3 | MSQFVGFGVQVELKDGKLIQGKIAKATSKGLTLNDVQFGDGGKSQAFKVRASRLKDLKVLTVASQSGKRKQQRQQQQQNDYNQNRGEHIDWQDDDVSKIKQREDFDFQRNLGMFNKKDVFAQLKQNDDILPENRLRGHNRKQTQLQQNNYQNDELVIPDAKKDSWNKISSRNEQSTHQSQPQQDAQDDLVLEDDEHEYDVDDIDDPKYLPITQSLNITHLIHSATNSPSINDKTRGTVINDKDQVLAKLGQMIISQSRSNSTSLPAANKQTTIRSKNTKQNIPMATPVQLLEMESITSEFFSINSAGLLENFAVNASFFLKQKLGGRARLRLQNSNPEPLVVILASDSNRSGAKALALGRHLCQTGHIRVITLFTCSQNELQDSMVKKQTDIYKKCGGKIVNSVSSLESAMETLNSPVEIVIDAMQGYDCTLSDLAGTSEVIESRIKSMISWCNKQRGSTKVWSLDIPNGFDAGSGMPDIFFSDRIEATGIICSGWPLIAINNLIANLPSLEDAVLIDIGIPQGAYSQRTSLRKFQNCDLFVTDGSLLLDL |
| Edc3^1-66^ | MSQFVGFGVQVELKDGKLIQGKIAKATSKGLTLNDVQFGDGGKSQAFKVRASRLKDLKVLTVASQS |
| Edc3^67-282^ | GKRKQQRQQQQQNDYNQNRGEHIDWQDDDVSKIKQQEDFDFQRNLGMFNKKDVFAQLKQNDDILPENRLRGHNRKQTQLQQNNYQNDELVIPDAKKDSWNKISSRNEQSTHQSQPQQDAQDDLVLEDDEHEYDVDDIDDPKYLPITQSLNITHLIHSATNSPSINDKTKGTVINDKDQVLAKLGQMIISQSRSNSTSLPAANKQTTIRSKNTKQNI |
| Edc3^283-551^ | PMATPVQLLEMESITSEFFSINSAGLLENFAVNASFFLKQKLGGRARLRLQNSNPEPLVVILASDSNRSGAKALALGRHLCQTGHIRVITLFTCSQNELQDSMVKKQTDIYKKCGGKIVNSVSSLESAMETLNSPVEIVIDAMQGYDCTLSDLAGTSEVIESRIKSMISWCNKQRGSTKVWSLDIPNGFDAGSGMPDIFFSDRIEATGIICSGWPLIAINNLIANLPSLEDAVLIDIGIPQGAYSQRTSLRKFQNCDLFVTDGSLLLDL |
| Lsm1-7 | Lsm1:  MSANSKDRNQSNQDAKRQQQNFPKKISEGEADLYLDQYNFTTTAAIVSSVDRKIFVLLRDGR  MLFGVLRTFDQYANLILQDCVERIYFSEENKYAEEDRGIFMIRGENVVMLGEVDIDKEDQPLE  AMERIPFKEAWLTKQKNDEKRFKEETHKGKKMARHGIVYDFHKSDMY  Lsm2:  MLFFSFFKTLVDQEVVVELKNDIEIKGTLQSVDQFLNLKLDNISCTDEKKYPHLGSVRNIFIRGS  TVRYVYLNKNMVDTNLLQDATRREVMTERK  Lsm3:  METPLDLLKLNLDERVYIKLRGARTLVGTLQAFDSHCNIVLSDAVETIYQLNNEELSESERRCE  MVFIRGDTVTLISTPSEDDDGAVEI  Lsm4:  MLPLYLLTNAKGQQMQIELKNGEIIQGILTNVDNWMNLTLSNVTEYSEESAINSEDNAESSKA  VKLNEIYIRGTFIKFIKLQDNIIDKVKQQINSNNNSNSNGPGHKRYYNNRDSNNNRGNYNRRN  NNNGNSNRRPYSQNRQYNNSNSSNINNSINSINSNNQNMNNGLGGSVQHHFNSSSPQKVEF  Lsm5:  MSLPEILPLEVIDKTINQKVLIVLQSNREFEGTLVGFDDFVNVILEDAVEWLIDPEDESRNEKV  MQHHGRMLLSGNNIAILVPGGKKTPTEAL  Lsm6:  MSGKASTEGSVTTEFLSDIIGKTVNVKLASGLLYSGRLESIDGFMNVALSSATEHYESNNNKLL  NKFNSDVFLRGTQVMYISEQKI  Lsm7:  MHQQHSKSENKPQQQRKKFEGPKREAILDLAKYKDSKIRVKLMGGKLVIGVLKGYDQLMNL  VLDDTVEYMSNPDDENNTELISKNARKLGLTVIRGTILVSLSSAEGSDVLYMQK |
| Pat1 | MSFFGLENSGNARDGPLDFEESYKGYGEHELEENDYLNDETFGDNVQVGTDFDFGNPHSSGSSGNAIGGNGVGATARSYVAATAEGISGPRTDGTAAAGPLDLKPMESLWSTAPPPAMAPSPQSTMAPAPAPQQMAPLQPILSMQDLERQQRQMQQQFMNFHAMGHPQGLPQGPPQQQFPMQPASGQPGPSQFAPPPPPPGVNVNMNHMPMGPVQVPVQASPSPIGMSNTPSPGPVVGATNMPLQSGRRSKRDVSPEEQRRLQIRHARVEKILKYSGIMTPRDKDFITRYQLSHIVTEDPYNEDFYFQVYKIIQRGGITSESNKGLIARAYLEHSGHRLGGRYKRTDIALQRMQSQVEKAVTVAKERPSKLKDQQAAAGNSSQDNKQANTVLGKISSTLNSKNPRRQLQIPRQQPSDPDALKDVTDSLTNVDLASSGSSSTGSSAAAVASKQRRRSSYAFNNGNGATNLNKSGGKKFILELIETVYEEILDLEANLRNGQQTDSTAMWEALHIDDSSYDVNPFISMLSFDKGIKIMPRIFNFLDKQQKLKILQKIFNELSHLQIIILSSYKTTPKPTLTQLKKVDLFQMIILKIIVSFLSNNSNFIEIMGLLLQLIRNNNVSFLTTSKIGLNLITILISRAALIKQDSSRSNILSSPEISTWNEIYDKLFTSLESKIQLIFPPREYNDHIMRLQNDKFMDEAYIWQFLASLALSGKLNHQRIIIDEVRDEIFATINEAETLQKKEKELSVLPQRSQELDTELKSIIYNKEKLYQDLNLFLNVMGLVYRDGEISELK |
| Pat1^1-240^ | MSFFGLENSGNARDGPLDFEESYKGYGEHELEENDYLNDETFGDNVQVGTDFDFGNPHSSGSSGNAIGGNGVGATARSYVAATAEGISGPRTDGTAAAGPLDLKPMESLWSTAPPPAMAPSPQSTMAPAPAPQQMAPLQPILSMQDLERQQRQMQQQFMNFHAMGHPQGLPQGPPQQQFPMQPASGQPGPSQFAPPPPPPGVNVNMNQMPMGPVQVPVQASPSPIGMSNTPSPGPVVGAT |
| Pat1^241-422^ | KMPLQSGRRSKRDLSPEEQRRLQIRHAKVEKILKYSGLMTPRDKDFITRYQLSQIVTEDPYNEDFYFQVYKIIQRGGITSESNKGLIARAYLEHSGHRLGGRYKRTDIALQRMQSQVEKAVTVAKERPSKLKDQQAAAGNSSQDNKQANTVLGKISSTLNSKNPRRQLQIPRQQPSDPDALK |
| Pat1^423-796^ | DVTDSLTNVDLASSGSSSTGSSAAAVASKQRRRSSYAFNNGNGATNLNKSGGKKFILELIETVYEEILDLEANLRNGQQTDSTAMWEALHIDDSSYDVNPFISMLSFDKGIKIMPRIFNFLDKQQKLKILQKIFNELSHLQIIILSSYKTTPKPTLTQLKKVDLFQMIILKIIVSFLSNNSNFIEIMGLLLQLIRNNNVSFLTTSKIGLNLITILISRAALIKQDSSRSNILSSPEISTWNEIYDKLFTSLESKIQLIFPPREYNDHIMRLQNDKFMDEAYIWQFLASLALSGKLNHQRIIIDEVRDEIFATINEAETLQKKEKELSVLPQRSQELDTELKSIIYNKEKLYQDLNLFLNVMGLVYRDGEISELK |
| Xrn1 | MGIPKFFRYISERWPMILQLIEGTQIPEFDNLYLDMNSILHNCTHGNDDDVTKRLTEEEVFAKI  CTYIDHLFQTIKPKKIFYMAIDGVAPRAKMNQQRARRFRTAMDAEKALKKAIENGDEIPKGEPFDSNSITPGTEFMAKLTKNLQYFIHDKISNDSKWREVQIIFSGHEVPGEGEHKIMNFIRHLKSQKDFNQNTRHCIYGLDADLIMLGLSTHGPHFALLREEVTFGRRNSEKKSLEHQNFYLLHLSLLREYMELEFKEIADEMQFEYNFERILDDFILVMFVIGNDFLPNLPDLHLNKGAFPVLLQTFKEALLHTDGYINEHGKINLKRLGVWLNYLSQFELLNFEKDDIDVEWFNKQLENISLEGERKRQRVGKKLLVKQQKKLIGSIKPWLMEQLQEKLSPDLPDEEIPTLELPKDLDMKDHLEFLKEFAFDLGLFITHSKSKGSYSLKMDLDSINPDETEEEFQNRVNSIRKTIKKYQNAIIVEDKEELETEKTIYNERFERWKHEYYHDKLKFTTDSEEKVRDLAKDYVEGLQWVLYYYYRGCPSWSWYYPHHYAPRISDLAKGLDQDIEFDLSKPFTPFQQLMAVLPERSKNLIPPAFRPLMYDEQSPIHDFYPAEVQLDKNGKTADWEAVVLISFVDEKRLIEAMQPYLRKLSPEEKTRNQFGKDLIYSFNPQVDNLYKSPLGGIFSDIEHNHCVEKEYITIPLDSSEIRYGLLPNAKLGAEMLAGFPTLLSLPFTSSLEYNETMVFQQPSKQQSMVLQITDIYKTNNVTLEDFSKRHLNKVIYTRWPYLRESKLVSLTDGKTIYEYQESNDKKKFGFITKPAETQDKKLFNSLKNSMLRMYAKQKAVKIGPMEAIATVFPVTGLVRDSDGGYIKTFSPTPDYYPLQLVVESVVNEDERYKERGPIPIEEEFPLNSKVIFLGDYAYGGETTIDGYSSDRRLKITVEKKFLDSEPTIGKERLQMDHQAVKYYPSYIVSKNMHLHPLFLSKITSKFMITDATGKHINVGIPVKFEARHQKVLGYARRNPRGWEYSNLTLNLLKEYRQTFPDFFFRLSKVGNDIPVLEDLFPDTSTKDAMNLLDGIKQWLKYVSSKFIAVSLESDSLTKTSIAAVEDHIMKYAANIEGHERKQLAKVPREAVLNPRSSFALLRSQKFDLGDRVVYIQDSGKVPIFSKGTVVGYTTLSSSLSIQVLFDHEIVAGNNFGGRLRTNRGLGLDASFLLNITNRQFIYHSKASKKALEKKKQSNNRNNNTKTAHKTPSKQQSEEKLRKERAHDLLNFIKKDTNEKNSESVDNKSMGSQKDSKPAKKVLLKRPAQKSSENVQVDLANFEKAPLDNPTVAGSIFNAVANQYSDGIGSNLNIPTPPHPMNVVGGPIPGANDVADVGLPYNIPPGFMTHPNGLHPLHPHQMPYPNMNGMSIPPPAPHGFGQPISFPPPPPMTNVSDQGSRIVVNEKESQDLKKFINGKQHSNGSTIGGETKNSRKGEIKPSSGTNSTECQSPKSQSNAADRDNKKDEST |

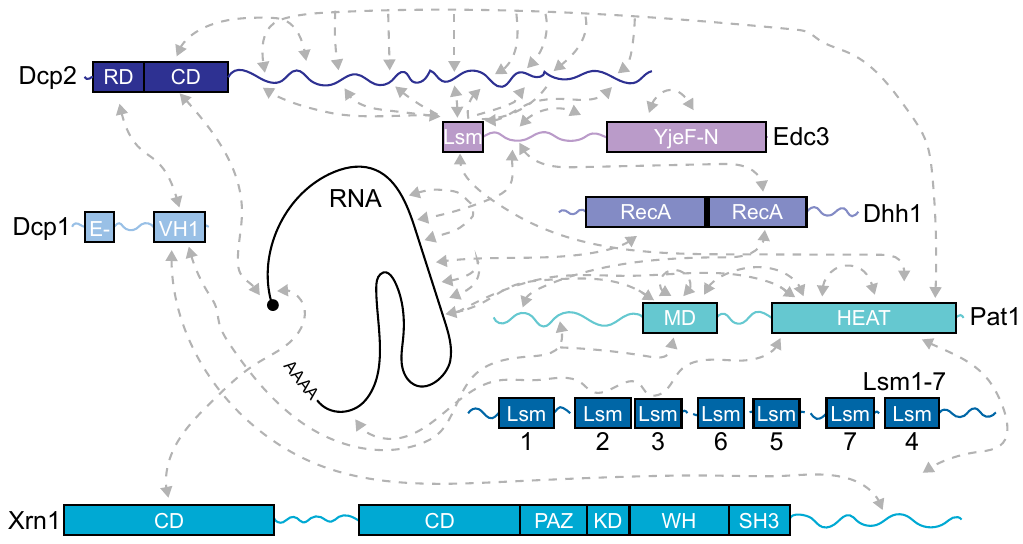

**Figure S1. Multivalent interactions between P-body proteins and RNA.**

Schematic of P-body proteins and RNA as in Figure 1A. Gray boxes indicate structured domains, and squiggly lines denote intrinsically disordered regions (IDRs). Domain abbreviations follow previously established nomenclature: Dcp1, Ena/VASP Homology 1 (EVH1)(5); Dcp2, Regulatory Domain (RD) and Catalytic Domain (CD)(6); Dhh1, RecA-like domain (RecA)(7); Edc3, Like Sm (Lsm)(8), YjeF N terminal protein domain (YjeF-N)(9); Lsm1-7, Like Sm (Lsm)(10); Pat1, Middle Domain (MD), Huntington-EF3-PP2A-TOR1 like domain super helix (HEAT)(10); Xrn1, Catalytic Domain (CD), Piwi-Argonaute-Zwille domain (PAZ), Kyrpides-Ouzounis-Woese domain (KD), Winged Helix(WH), SRC Homology domain 3 like domain (SH3) (11). Gray dotted lines denote interactions. References for interactions are: Dcp1 (6, 12, 13), Dcp2 (6, 8, 13-15), Dhh1 (7, 13, 16), Edc3 (8, 9, 14, 16, 17), Lsm1-7 (10, 13), Pat1 (10, 13, 15, 18), and Xrn1 (11, 12). Multivalent interactions between P-body proteins and RNA involve both structured domains and IDRs.

See also Fig. 1A, and S2-S3.

**Figure S2. Structured domains within P-body proteins.**

Structures for:

(A) Xrn1 CD, PAZ, KD, WH, and SH3 domains (11).

(B) Pat1 HEAT domain and Lsm1-7 (10).

(C) Dcp1, Dcp2 RD and CD domains, and Edc3 Lsm domain (19).

(D)Edc3 YjeF-N domain dimer (9).

(E) Dhh1 RecA domains and Edc3 M domain (16).

(F) Structural model for the middle domain (MD) of Pat1 (Pat1^241-422^) was generated using AlphaFold (20, 21). Secondary structure and disorder predictions are in agreement that Pat1 MD contains structured elements (Figure S3). However, these α-helices likely lack a defined and stable stereospecific three-dimensional structure due to: the lack of extensive tertiary contacts within the MD domain or to the HEAT domain, the flexible linkers between the α-helices, and consequently the low confidence of this model for the regions outside of the α-helices. See the following website for further details: <https://alphafold.ebi.ac.uk/entry/P25644>.

Metal ions are colored orange in Xrn1 (Mg^2+^), Dcp2 (Mg^2+^), and Pat1/Lsm1-7 (Co^2+^) structures, and RNA (oligo)nucleotides are colored black in Xrn1 and Dcp2 structures. The structured domains of P-body proteins are interaction surfaces that increase the valency within the P-body networks. Note that Pat1 HEAT domain also dimerizes (15), Pat1 N domain binds to Dhh1 in a similar position as Edc3 M domain (16), and Xrn1 C domain binds to Dcp1 (12); these structures have been omitted for brevity.

See also Fig. 1A, S3, and Methods.

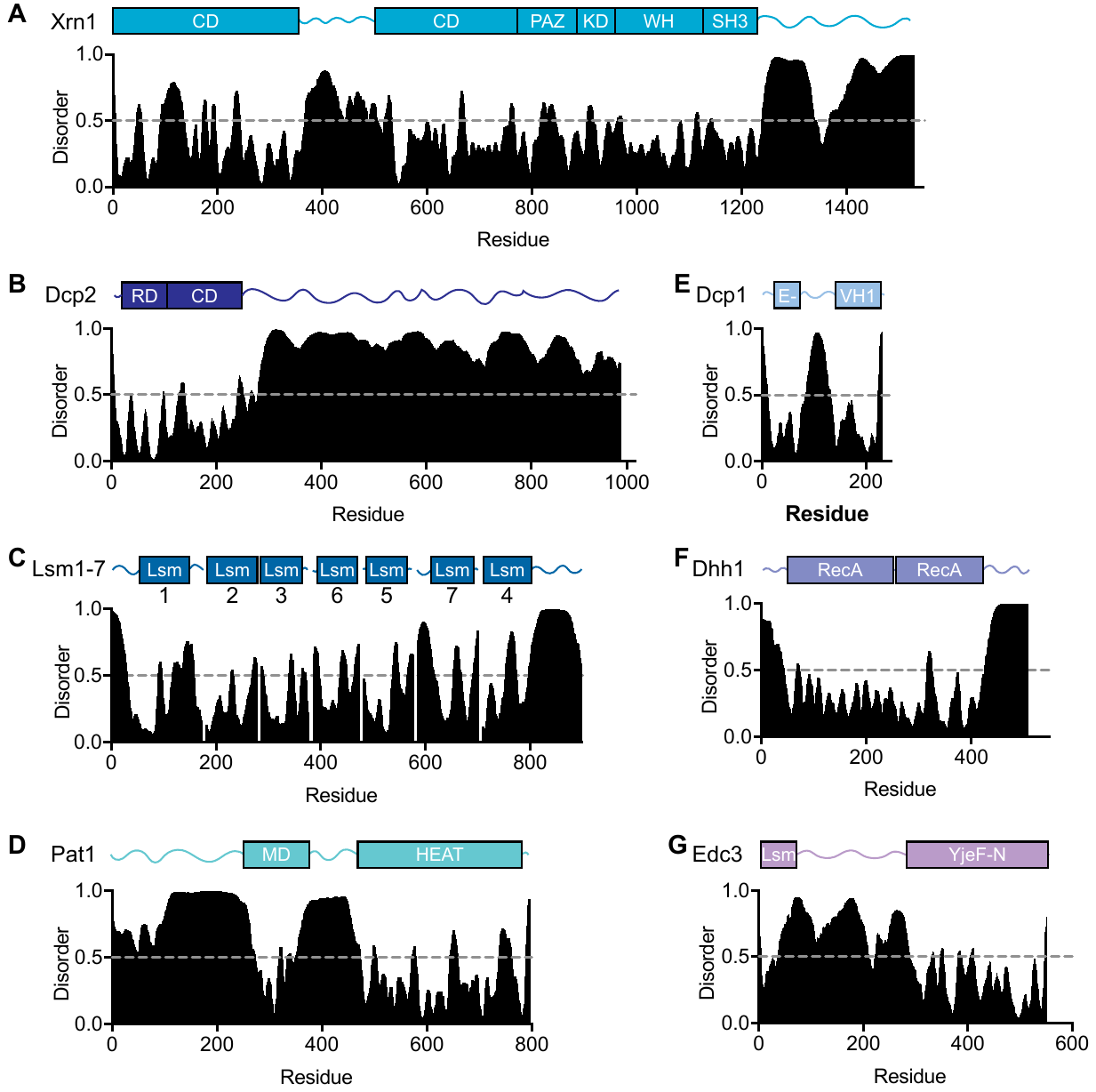

**Figure S3. Intrinsically disordered regions within P-body proteins.**

Disorder score was calculated for P-body proteins using Disprot VSL2 (22). Higher scores suggest disordered regions and lower scores suggest structured domains.

See also Fig. 1A and S2.

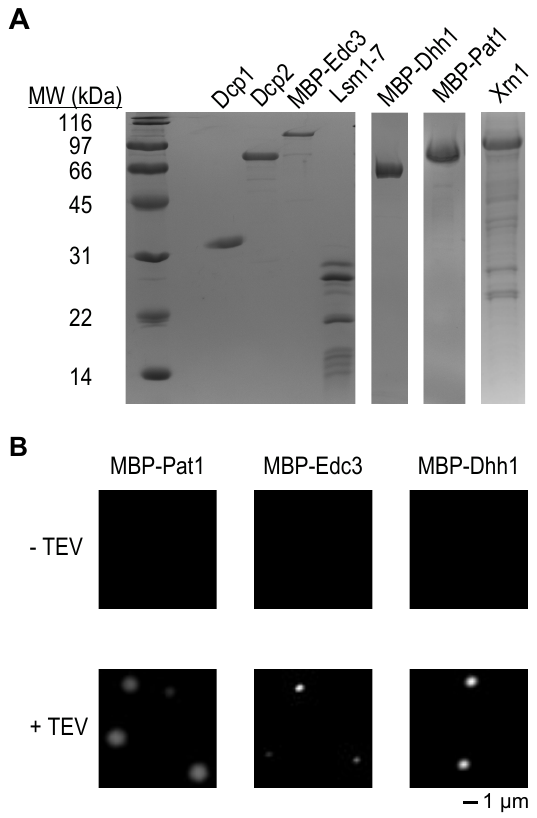

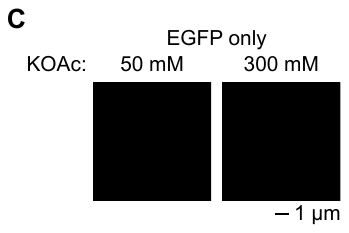

**Figure S4. Purified P-body proteins and TEV cleavage initiation of condensate formation.**

(A) SDS PAGE gel of purified P-body proteins used in this study. Proteins were purified using a four step purification process: Ni^2+^ affinity, amylose affinity, ion exchange, and size exclusion columns. Dhh1, Edc3, and Pat1 required the MBP tag for purification to prevent loss of protein. See methods for details.

(B) Using the purified proteins, TEV cleavage of the MBP tag was used to initiate phase separation. TEV was added at a 50:1 P-body protein:TEV ratio as higher amounts of TEV perturbed condensate formation and maturation.

(C) EGFP (1 μM) does not form condensates at 50 mM nor 300 mM KOAc, pH 7. This suggests that condensates are formed via P-body protein interactions rather than by EGFP.

See also Fig. 1B-1C.

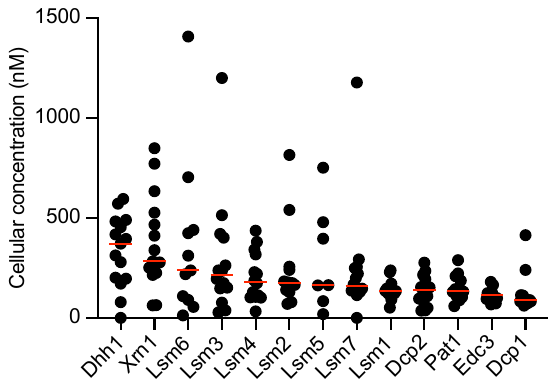

**Figure S5. Estimated cellular concentrations of P-body proteins.**

The absolute number of proteins in a Saccharomyces cerevisiae cell were obtained from Ho *et al.* (23). Briefly, Ho et al. combined mass spectrometry and fluorescent microscopy studies with quantitative proteome wide data. We converted the number of proteins per cell to concentration using the following assumptions: (i) cell volume of 62 µm^3^; (ii) P-body proteins were not restricted from any organelle as the cytoplasm makes up the majority of the cell volume, and most P-body proteins also reside in the nucleus which is the largest compartment after the cytoplasm (24). Each dot is from a single quantitative report and the red line indicates the median. The median cellular concentration estimates for P-body proteins range from 370 to 90 µM.

See also Fig. 1D.

**Figure S6. Disruption of homotypic condensates.**

(A) Representative micrographs for Pat1 (top), Edc3 (middle), and Dhh1 (bottom). All reactions were conducted at pH 7 with the following added: 50 mM KOAc buffer (1^st^ column), 300 mM KOAc (2^nd^ column), 10% glycerol (3^rd^ column), and 100 mM arginine (4^th^ column). All homotypic condensates were prevented by arginine, and all added molecules prevented Dhh1 condensates.

(B) Titration of arginine against Dhh1 (top), Edc3 (middle), and Pat1 (bottom) homotypic condensates. Arginine was added before condensate formation was initiated.

See also Fig. 1E.

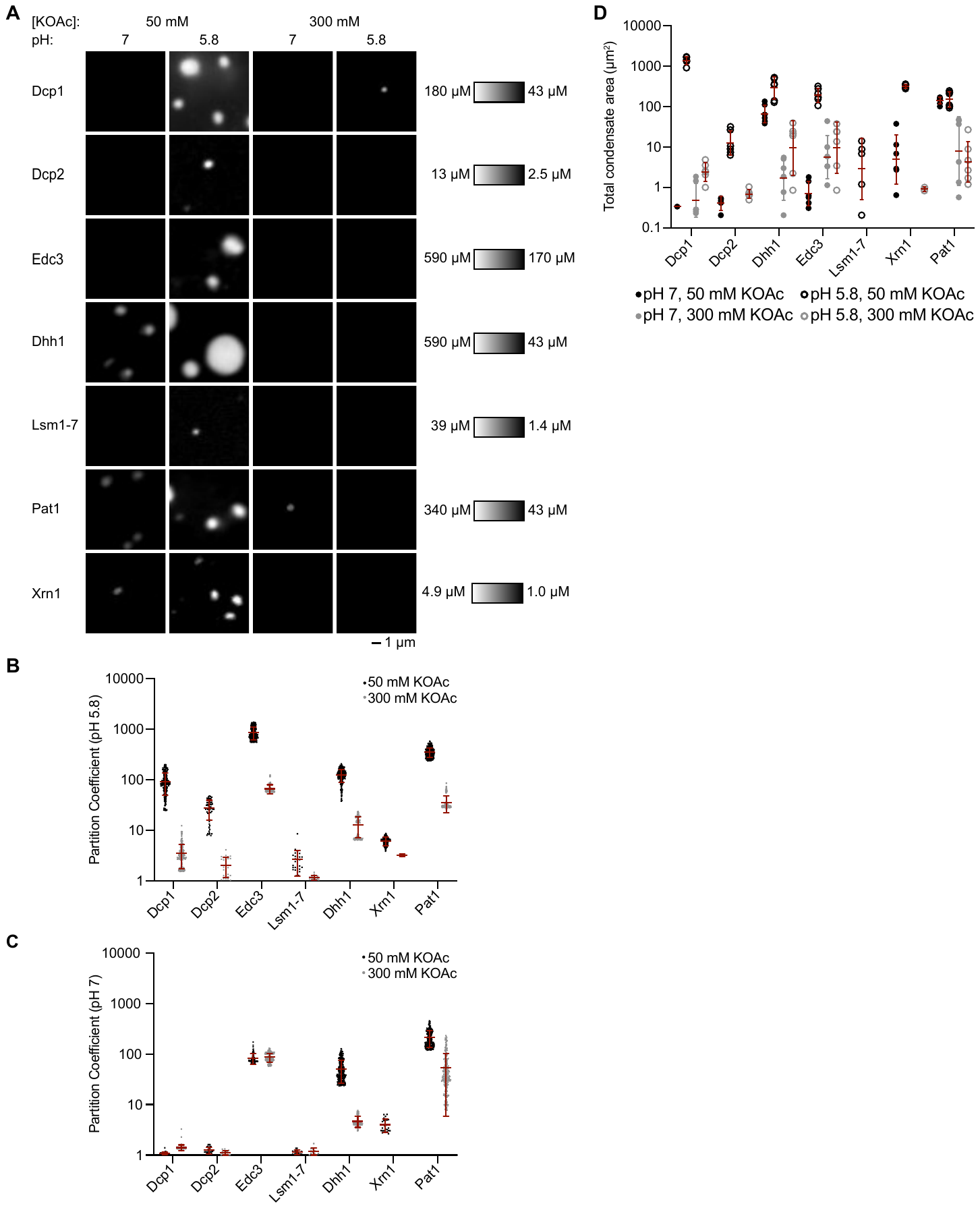

**Figure S7. Partition coefficients for individual P-body proteins at pH 7 and 5.8, and 50 and 300 mM KOAc.**

(A) Representative fluorescent micrographs for individual P-body proteins in all buffer conditions. Contrast is consistent for individual proteins across rows and is indicated on the right.

(B) Partition coefficients for individual P-body proteins at pH 5.8 for 50 mM KOAc (black) and 300 mM KOAc (gray).

(C) Partition coefficients for individual P-body proteins at pH 7 with 50 mM KOAc (black) and 300 mM KOAc (gray).

(D) Total condensate area for individual P-body proteins in all conditions. Individual data points represent the sum from one micrograph. Three micrographs were examined from each of two individual experiments for a total of six micrographs. Red bars indicate mean and standard deviation.

All individual P-body proteins exhibit enhanced partitioning into the condensed phase at pH 5.8, with the magnitude of the effect being most dramatic for Dcp1, Dcp2, and Edc3. Interestingly, pH 5.8 sensitizes Pat1 and especially Edc3 to electrostatic disruption with 300 mM KOAc.

See also Fig. 2A-B.

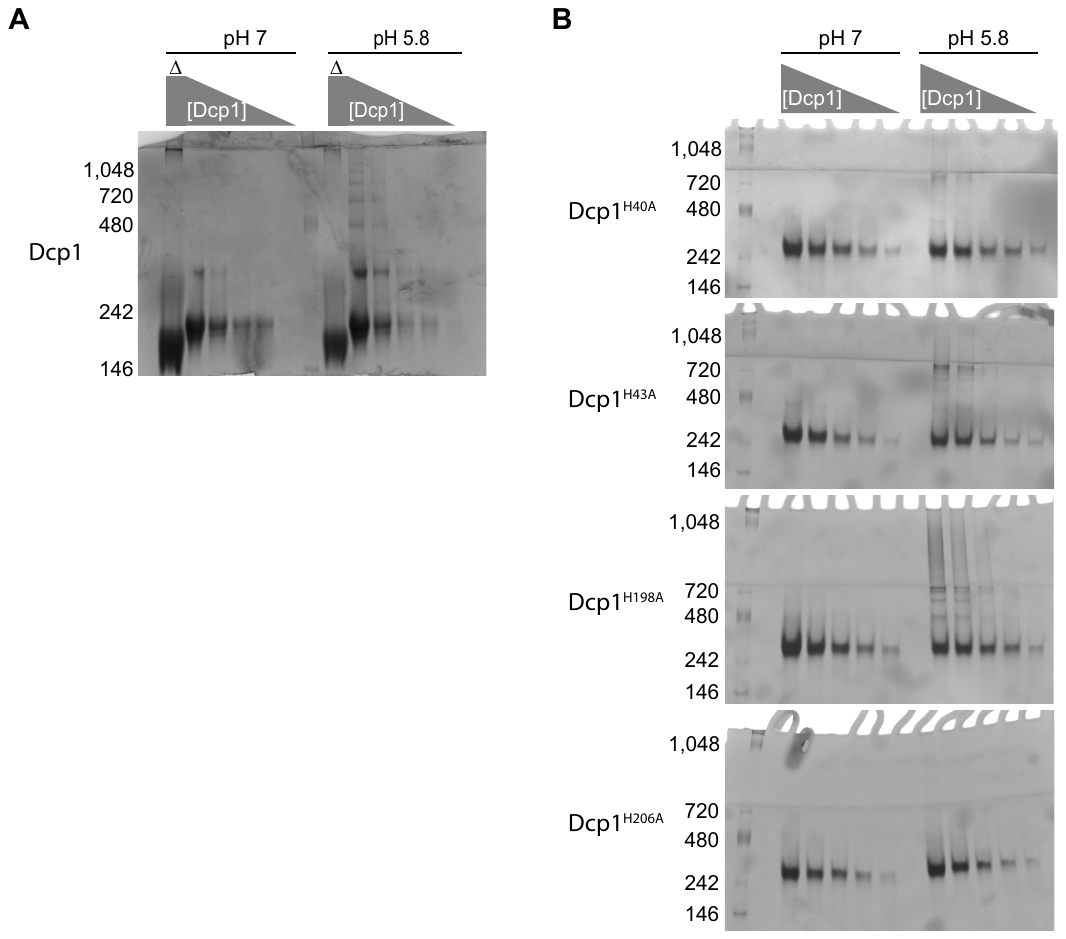

**Figure S8. H206A mutation disrupts Dcp1 oligomerization.**

(A) Native gel examining the oligomerization of Dcp1 at pH 7 (left) and pH 5.8 (right). Dcp1 concentration was diluted in twofold increments from 10 to 0.6 µM. The first lane has 10 µM Dcp1, and was incubated with SDS (Laemmli’s loading buffer) at 100 °C for 5 minutes before loading.

(B) Native gels examining oligomerization of Dcp1 histidine mutants. For histidine mutants the concentration range was from 5 to 0.3 µM.

Mutation of histidine 206 to alanine results in the most dramatic inhibition of Dcp1 LLPS and oligomerization at pH 5.8. The histidine 206 residues interact with each other at the dimer interface suggesting that this contact is important for Dcp1 LLPS and oligomerization. H40A and H43A mutants have a slight disruption in both assays, whereas mutation of histidine 198 has no impact. These data suggest that histidine 206 is particularly important for the pH-sensitivity of Dcp1 oligomerization and homotypic LLPS.

See also Fig. 2C-E .

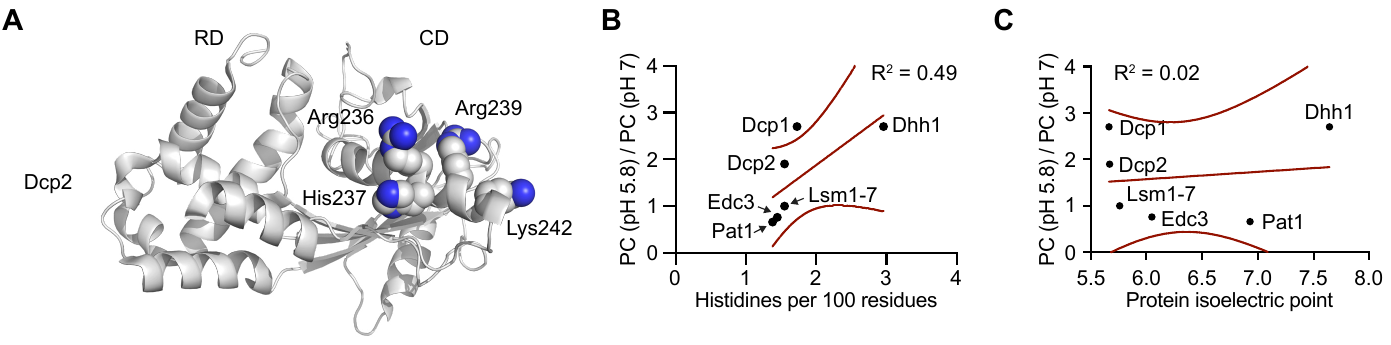

**Figure S9. Dcp2 structured domains and histidine density in P-body proteins.**

(A) Homology model of Dcp2 structured regulatory and catalytic domains with residues within previously identified RNA-binding region highlighted (6).

(B) Ratio of partition coefficients (pH 5.8 / pH 7 in 300 mM KOAc) versus the frequency of histidine residues (number of histidine residues per 100 amino acids) for all P-body proteins that were tested. Solid line indicates linear fit and dotted lines indicate 95% confidence interval.

(C) Ratio of partition coefficients (pH 5.8 / pH 7 in 300 mM KOAc) versus the protein isoelectric point for all P-body proteins that were tested. Solid line indicates linear fit and dotted lines indicate 95% confidence interval.

There is a reasonable correlation between histidine density and pH-sensitive condensate formation (R^2^ = 0.49) that is enhanced if Dcp1 is not considered (Figure 2I: R^2^ = 0.75). In the outlier, Dcp1, a single dominant histidine residue (H206) controls pH-sensitive condensate formation. In contrast, the protein isoelectric point does not correlate with pH-sensitive condensate formation (R2 = 0.02). These data suggest that multiple pH-sensitive histidine residues contribute to homotypic condensate formation and are distributed throughout Dcp2 and the other P-body proteins.

See also Fig. 2C-I.

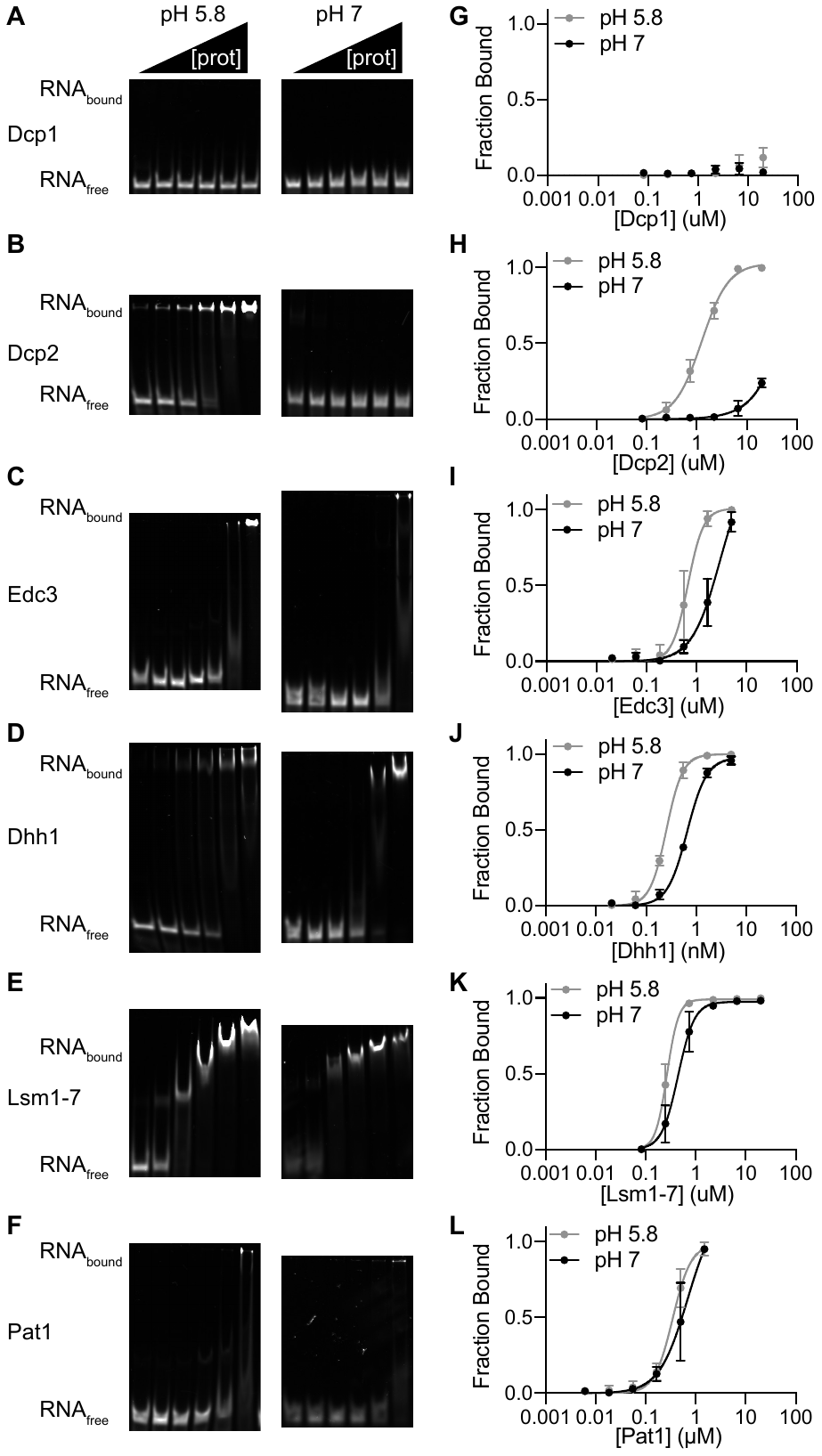

**Figure S10. Acidic pH promotes P-body protein binding to RNA.**

(A-F) EMSAs for P-body proteins binding to RPL41A RNA at pH 5.8 (left) and pH 7 (right). Gels are representative examples of three replicate experiments.

(G-L) Quantitation of *K_D_* values for P-body proteins binding to RNA. Data points and error bars correspond to the mean and standard deviation from three replicate experiments.

See also Table S2; Fig. 3A-C and S11.

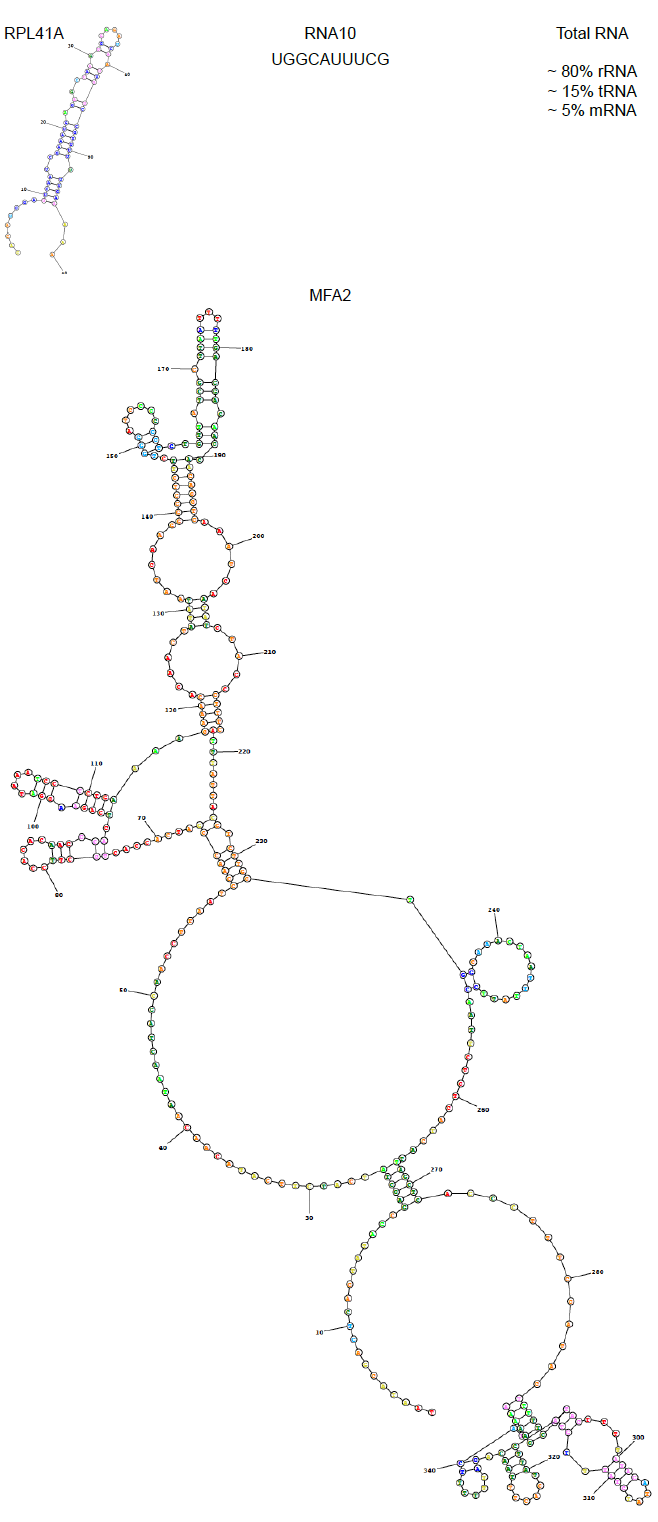

**Figure S11. Types of RNA used in this study.**

Portions of RPL41A covering the 5’ UTR and translational start site, and the 3’UTR and start of the polyA tail were combined in this 60 nucleotide RNA. RNA10 is a small single-stranded RNA previously used in RNA helicase studies (25). Total RNA contains more diverse RNA types and features (length, secondary structure, etc.). Yeast Mating Factor A (MFA2) is a full-length mRNA (348 nucleotides) that localizes to cellular P bodies (26). Predicted secondary structures for RPL41A and MFA2 from the RNAstructure website is shown (27).

See also Fig. 3A-C, 5B, S10, S19, S22, and S23.

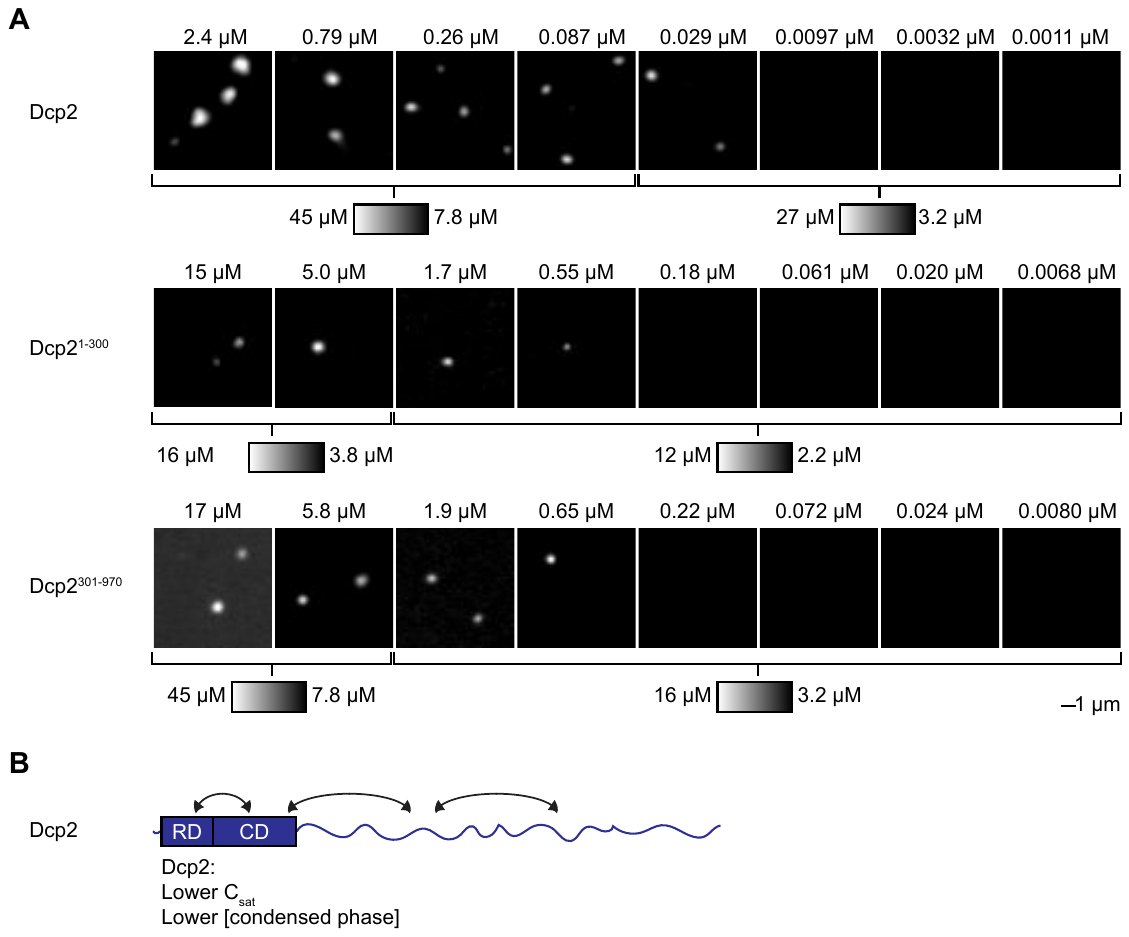

**Figure S12. Titration of Dcp2 truncations.**

(A) Representative micrographs for titration of Dcp2 and truncations. Total concentration of protein and contrast are labeled above and below the micrographs, respectively.

(B) Cartoon schematic representing: Dcp2 has a lower saturation concentration (C_sat_) than either truncation, suggesting that the structured N-terminal domains and disordered C-terminal regions synergize to drive C_sat_ lower in full-length Dcp2. The condensate concentration for Dcp2^301-970^ is higher, albeit at higher total protein concentrations.

See also Fig. 4A and 4K.

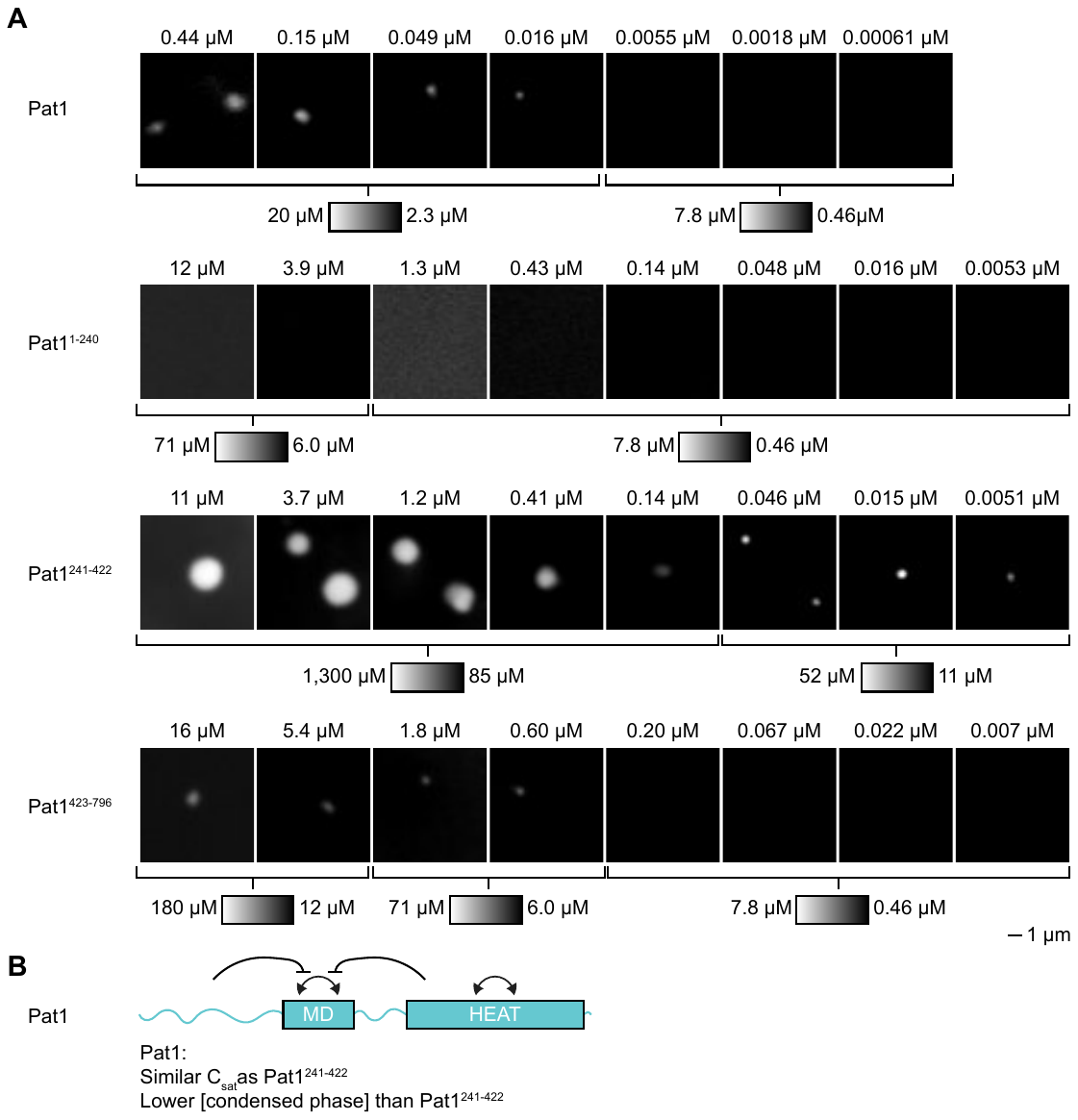

**Figure S13. Titration of Pat1 truncations.**

(A) Representative micrographs for titration of Pat1 and truncations. Total concentration of protein and contrast are labeled above and below the micrographs, respectively.

(B) Cartoon schematic representing: Pat1 has a similar saturation concentration (C_sat_) as Pat1^241-422^, and Pat1^241-422^ has a higher condensate protein concentration at comparable total protein concentrations. These data suggest that either the Pat1^1-240^ and/or Pat1^423-796^ interact with Pat1^241-422^ to modulate LLPS in Pat1.

See also Fig. 4B and 4L.

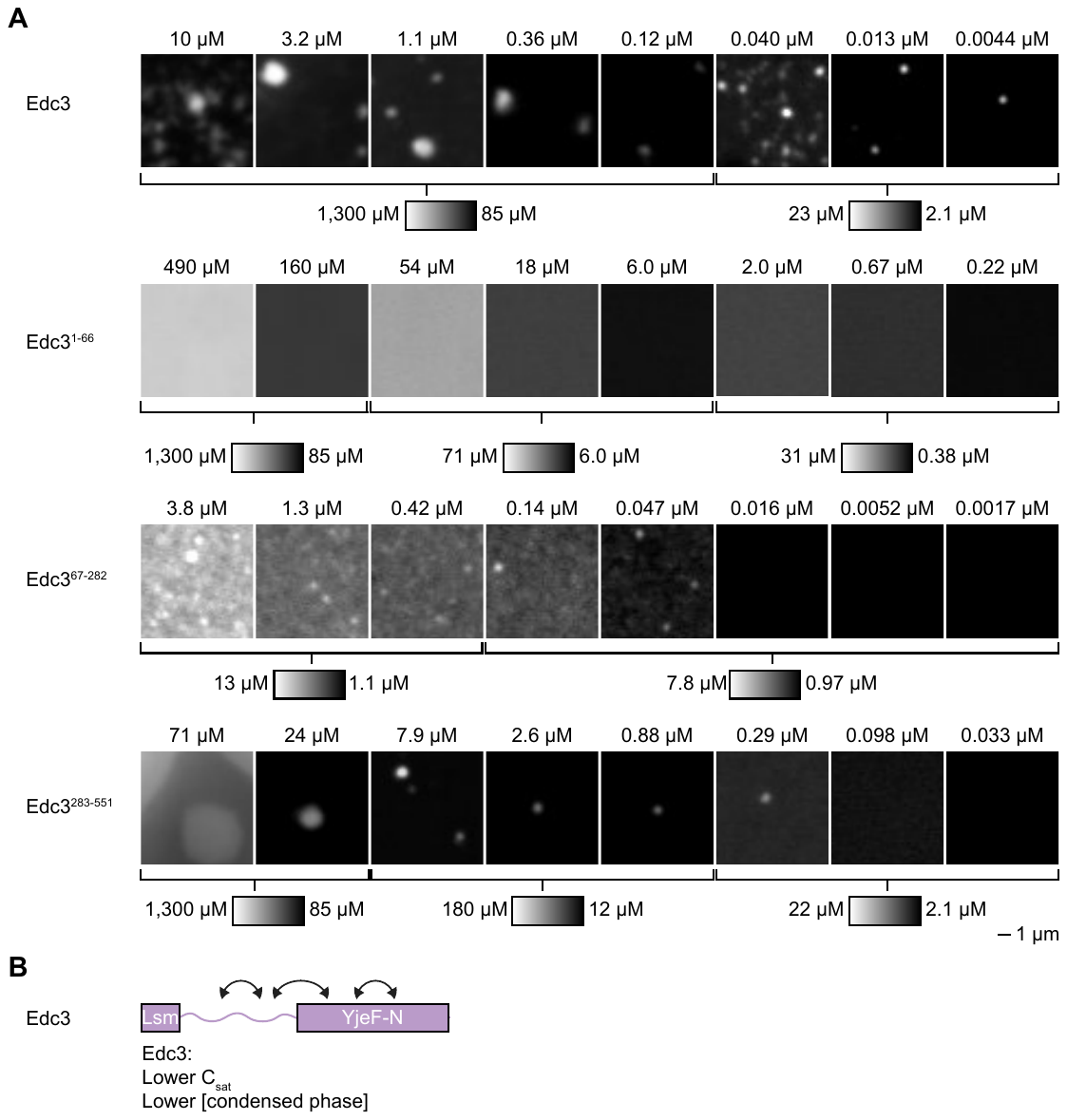

**Figure S14. Titration of Edc3 truncations.**

(A) Representative micrographs for titration of Edc3 and truncations. Total concentration of protein and contrast are labeled above and below the micrographs, respectively.

(B) Cartoon schematic representing: Edc3 has a lower saturation concentration (C_sat_) than any of the truncations, suggesting that the disordered middle region and the structured YjeF-N domain synergize to drive C_sat_ lower in full-length Edc3. The condensate protein concentration for Edc3^283-551^ is higher, albeit at higher total protein concentrations.

See also Fig. 4C and 4M.

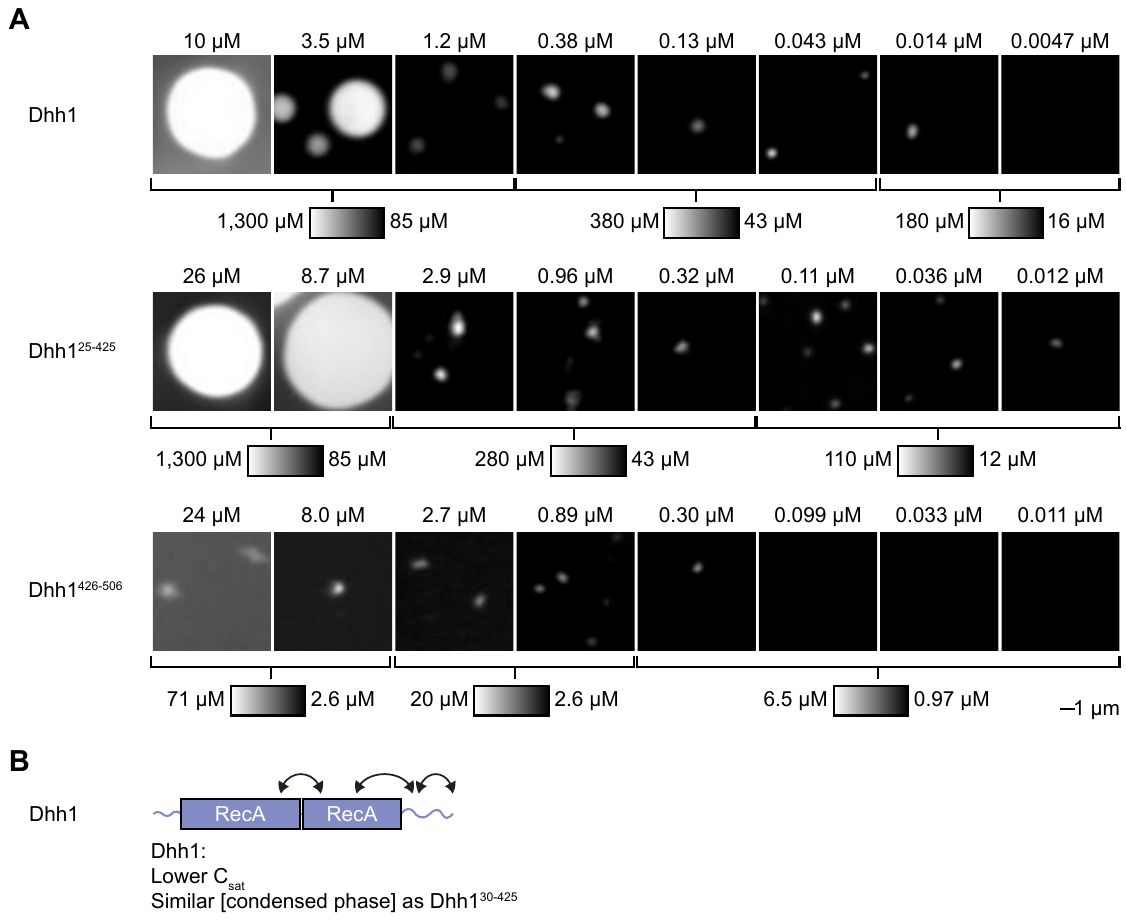

**Figure S15. Titration of Dhh1 truncations.**

(A) Representative micrographs for titration of Dhh1 and truncations. Total concentration of protein and contrast are labeled above and below the micrographs, respectively.

(B) Cartoon schematic representing: Dhh1 has a lower saturation concentration (C_sat_) than either of the truncations, suggesting that the structured tandem RecA domains and the disordered C-terminal region synergize to drive C_sat_ lower in full-length Dhh1. The condensate protein concentration for Dhh1^30-425^ is similar to Dhh1 at comparable total protein concentrations.

See also Fig. 4D and 4N.

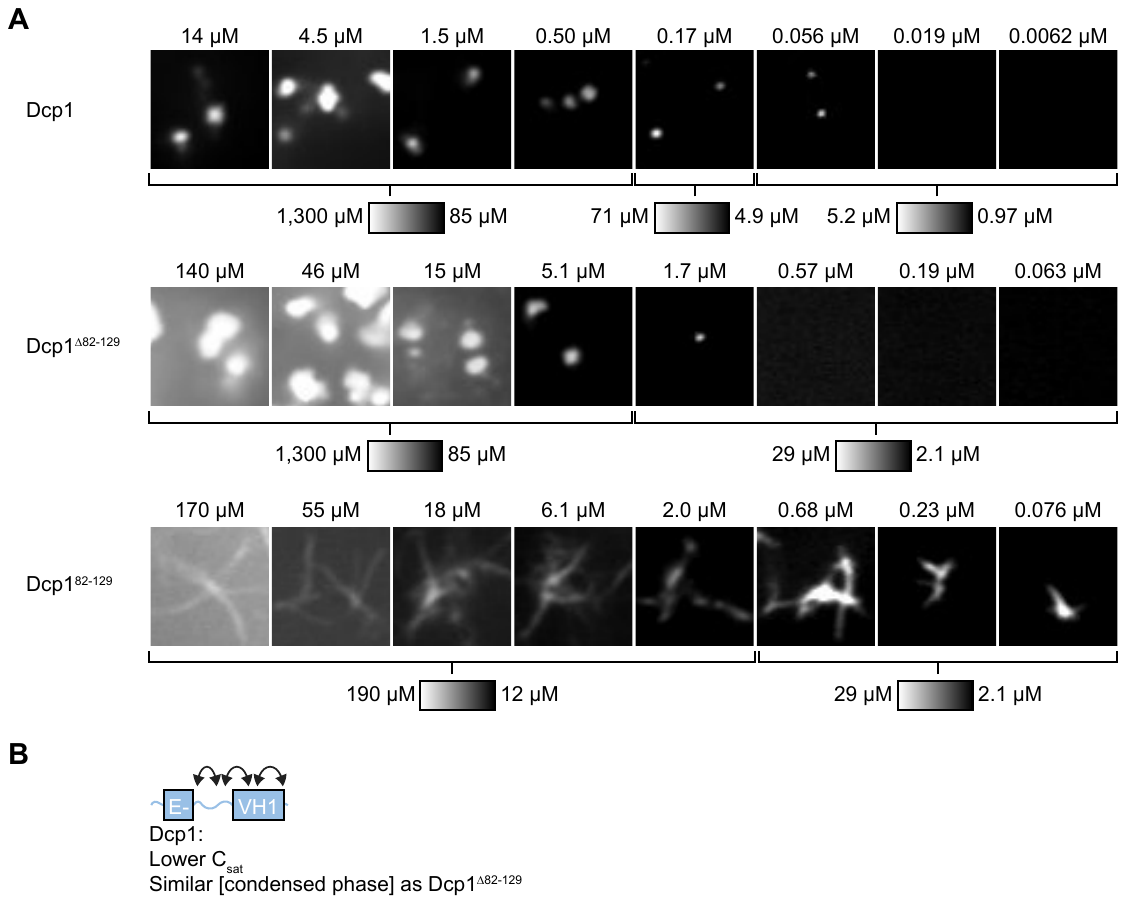

**Figure S16. Titration of Dcp1 truncations.**

(A) Representative micrographs for titration of Dcp1 and truncations. Total concentration of protein and contrast are labeled above and below the micrographs, respectively.

(B) Cartoon schematic representing: Dcp1 has a lower saturation concentration (C_sat_) than either of the truncations, suggesting that the structured EVH1 domain and the disordered middle linker synergize to drive C_sat_ lower in full-length Dcp1. The condensate protein concentration for Dcp1^Δ82-129^ is similar to Dcp1, albeit at slightly higher total protein concentrations.

See also Fig. 4E and 4O.

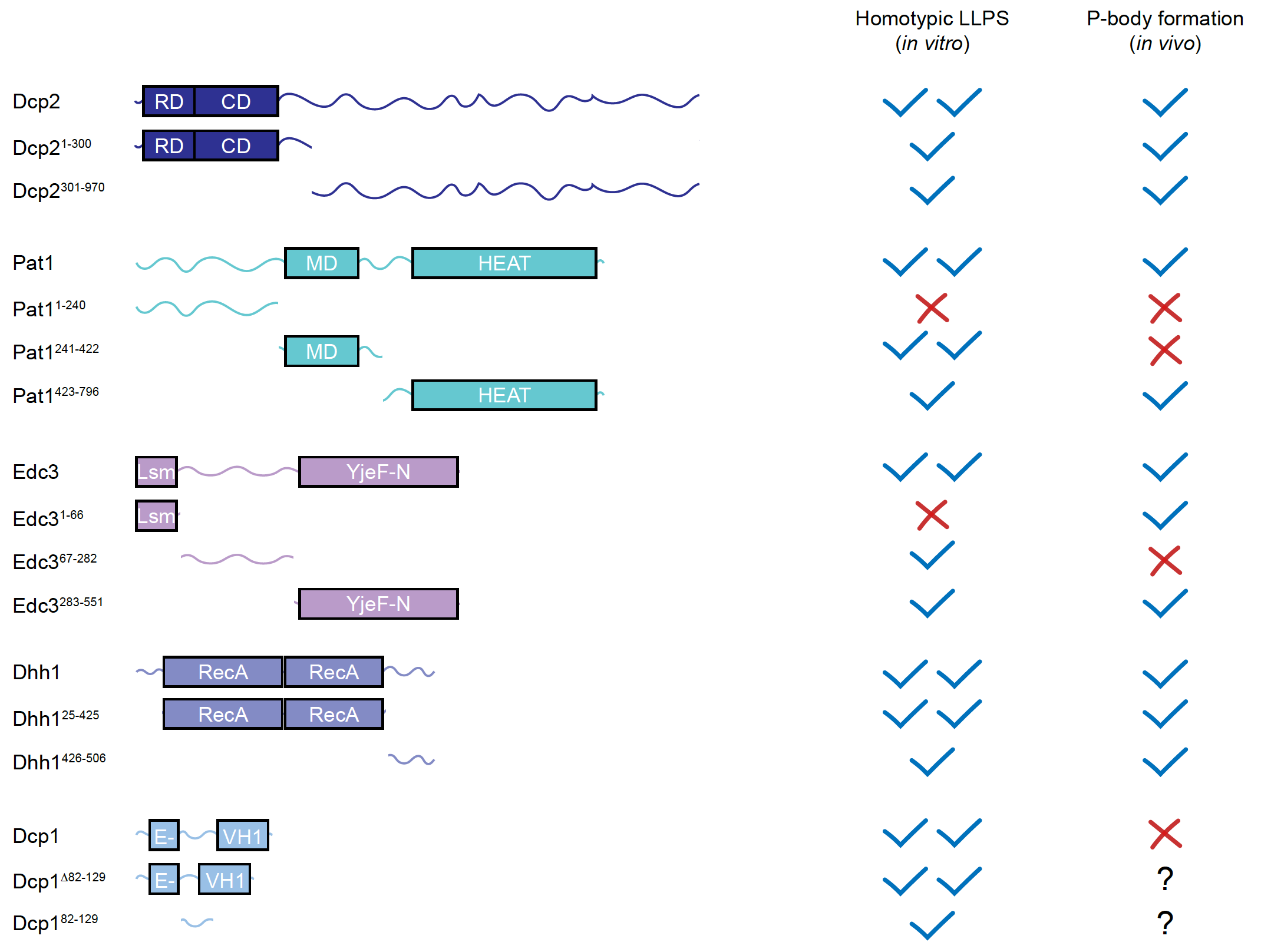

**Figure S17. Comparison between P-body protein regions forming homotypic condensates *in vitro* and contributing to P-body formation *in vivo*.**

The ability of P-body protein regions to exhibit homotypic LLPS *in vitro* (Fig. 4 and S12-16) and contribute to P-body formation *in vivo* (2-4, 18, 28) are scored in a qualitative fashion. Scoring for homotypic LLPS refers to experimental conditions and data from Figure 4. In general, there is reasonable agreement between forming homotypic condensates *in vitro* and contributing to P-body formation *in vivo*, despite the inherent differences in these experiments. Edc3^1-66^ does not form homotypic condensates but does contribute to P-body formation, likely via heterotypic interactions with helical-leucine motifs in Dcp2 that contribute to P-body formation (4, 8, 14). It is currently unclear why Pat1^241-422^ and Edc3^67-282^ form homotypic condensates but do not contribute to P-body formation in cells. It is possible that the ability of these domains to form condensates in cells is disrupted by interactions with other P-body components (3, 18), by less specific interactions with other cellular components (29), by posttranslational modifications (30), or via other mechanisms. Nonetheless, there is still reasonable overall agreement with the domains that exhibit LLPS *in vitro* and contribute to P-body formation *in vivo*.

See also Fig. 4, S12-16, and Discussion.

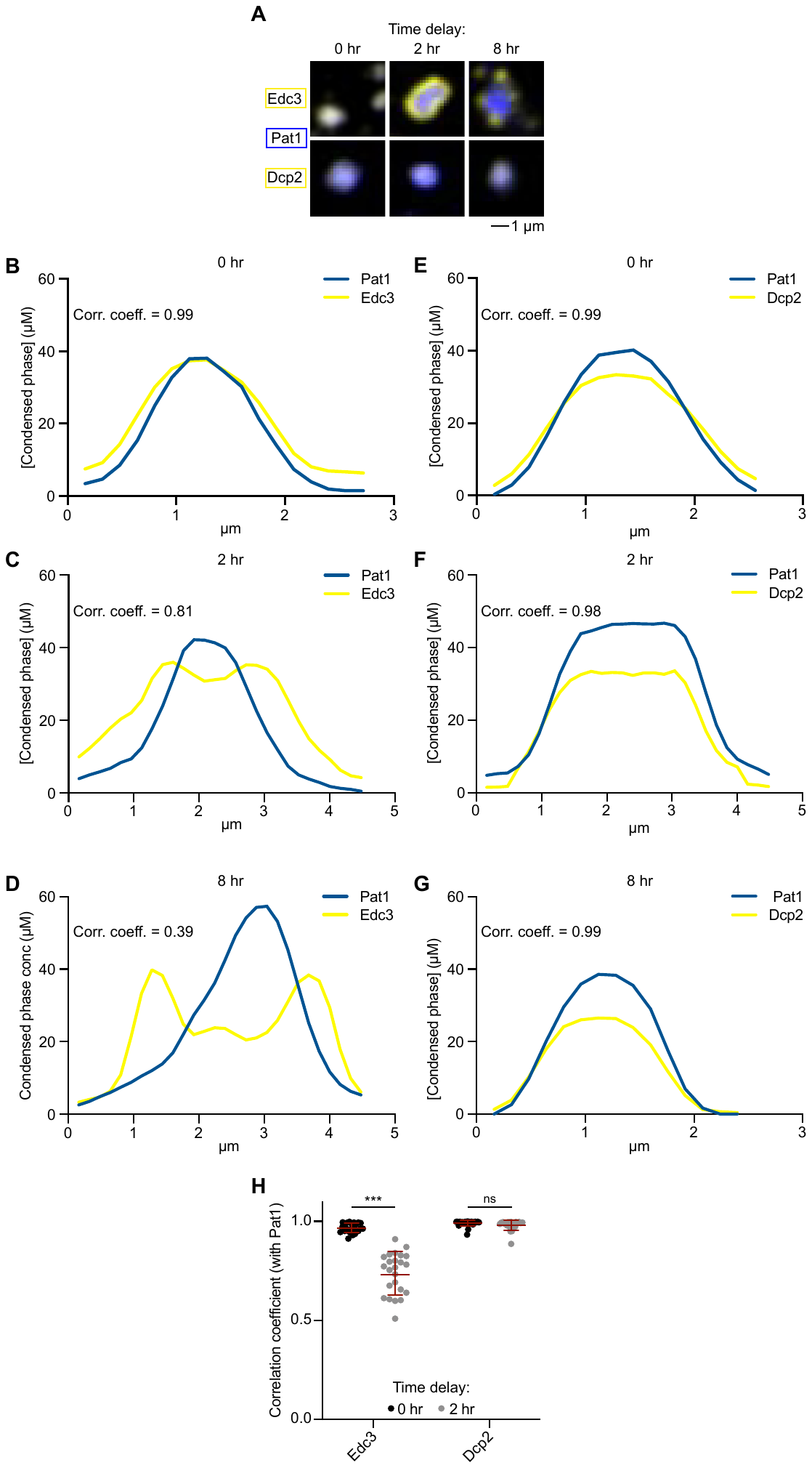

**Figure S18. Line profiles for overlap between Pat1 and Dcp2 or Edc3.**

(A) Representative micrographs for overlap between Edc3 or Dcp2 with Pat1 when Edc3 or Dcp2 are added at the time of condensate initiation (0 hr), and two or eight hours after condensate formation is initiated.

(B) Reactions were set up with all P-body proteins and RNA except Edc3. Edc3 was added back in as condensate formation was initiated (0 hr). Line profiles for Pat1 (blue) and Edc3 (yellow). Correlation coefficient between the two line profiles is displayed.

(C) As in (A) but Edc3 added 2 hours after condensate formation was initiated.

(D) As in (A) but Edc3 added 8 hours after condensate formation was initiated.

(E) As in (B) except Dcp2 was set aside instead of Edc3. Dcp2 was added back as condensate formation was initiated (0 hr).

(F) As in (D) but Dcp2 was added 2 hours after condensate formation was initiated.

(G) As in (D) but Dcp2 was added 8 hours after condensate formation was initiated.

(H) Quantification of the Pearson correlation coefficients of Edc3 and Dcp2 with Pat1 when added at condensate initiation (black) or after condensates have been formed for 2 hours (gray). Thirty condensates were measured from two replicate experiments.

During the course of our experiments, we observed that sufficient preincubation (1.5 hours) of all molecules together was required before initiating the reactions with TEV cleavage in order for different proteins to colocalize with one another. This suggests that a balance between heterotypic and homotypic interactions leads to P-body formation (and also that the *in vitro* system is slow to equilibrate). To further investigate this idea we intentionally investigated time as a variable in this set of experiments. We incubated different subsets of the P-body reconstitution separately for different amounts of time after TEV cleavage was initiated, before mixing the solutions together. We found that Dcp2 is readily recruited into the condensates containing the rest of the P-body proteins and RNA, at each timepoint tested. In contrast, Edc3 forms condensates that either coat (2 hr) or dock on the condensates with the rest of the P-body proteins and RNA. Images were taken 24 hr after TEV cleavage was initiated – and therefore 16-24 hr after the solutions were mixed together. Thus, these images likely represent the systems at equilibrium. One interpretation of these data is that homotypic condensates mature over time and fail to coalesce with heterotypic condensates. Indeed, alternative lines of investigation (Fig. 6 and S28-S31) also suggest that homotypic condensates mature more rapidly than heterotypic condensates. However, we note that additional plausible interpretations for this data exist – such as potential differences in surface tension between homotypic and heterotypic condensates (31).

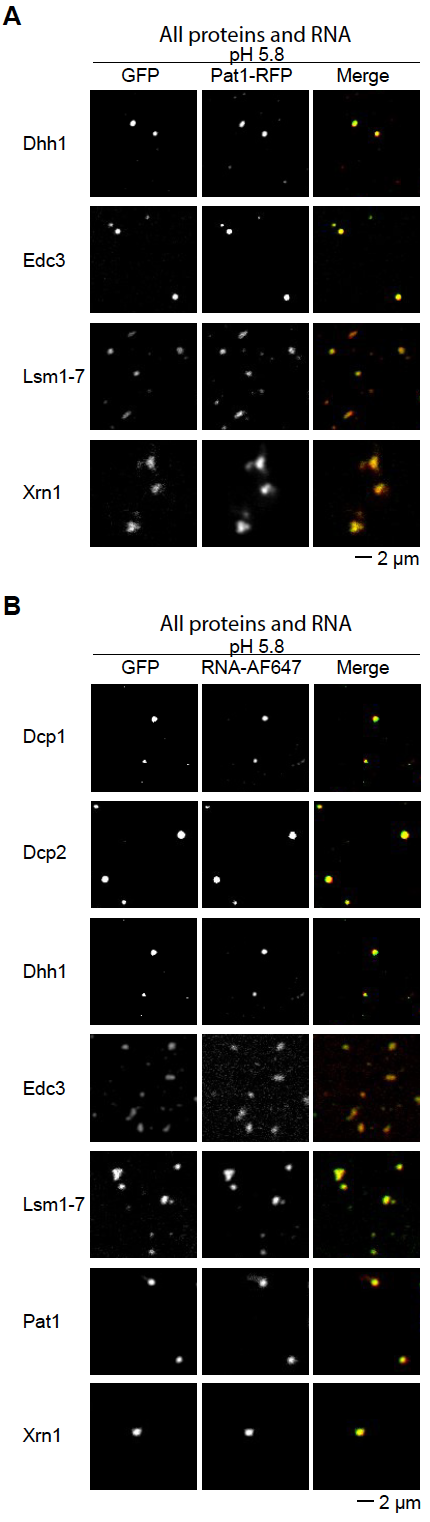

**Figure S19. All P-body proteins overlap with Pat1 and RNA in *in vitro* P-bodies under acidic (pH 5.8) conditions.**

(A) Representative micrographs for Dhh1, Edc3, and Lsm1-7 overlapping with Pat1 under acidic conditions (pH 5.8).

(B) Representative micrographs for Dcp1, Dcp2, Dhh1, Edc3, Lsm1-7, and Pat1 overlap with RNA under acidic conditions (pH 5.8).

Dcp1, Dcp2, Dhh1, and Edc3 are EGFP-fusion proteins and Lsm1-7 and Xrn1 are conjugated to AlexaFluor488.

See also Fig. 5A-B and S22.

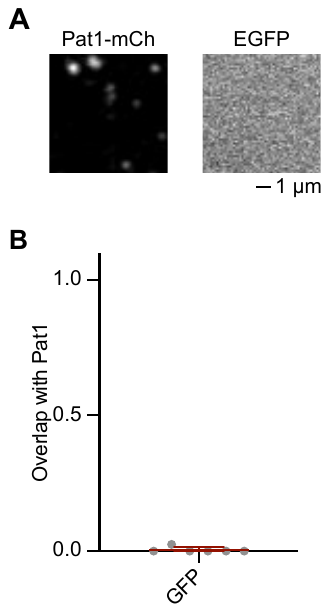

**Figure S20. EGFP does not partition into reconstituted P-bodies.**

(A) Representative micrograph of reconstituted P-bodies using Pat1-mCh as a marker, and with free EGFP (not tagged to any P-body protein) added. All P-body proteins and RNA were included in the reaction mixture, and P-bodies were formed at pH 5.8 and 300 mM KOAc.

(B) Quantification of overlap between condensates in the Pat1-mCh and EGFP channels.

EGFP does not partition into reconstituted P bodies, suggesting that the *in vitro* P bodies are formed by specific interactions involving P-body protein and RNA molecules.

See also Fig. 5.

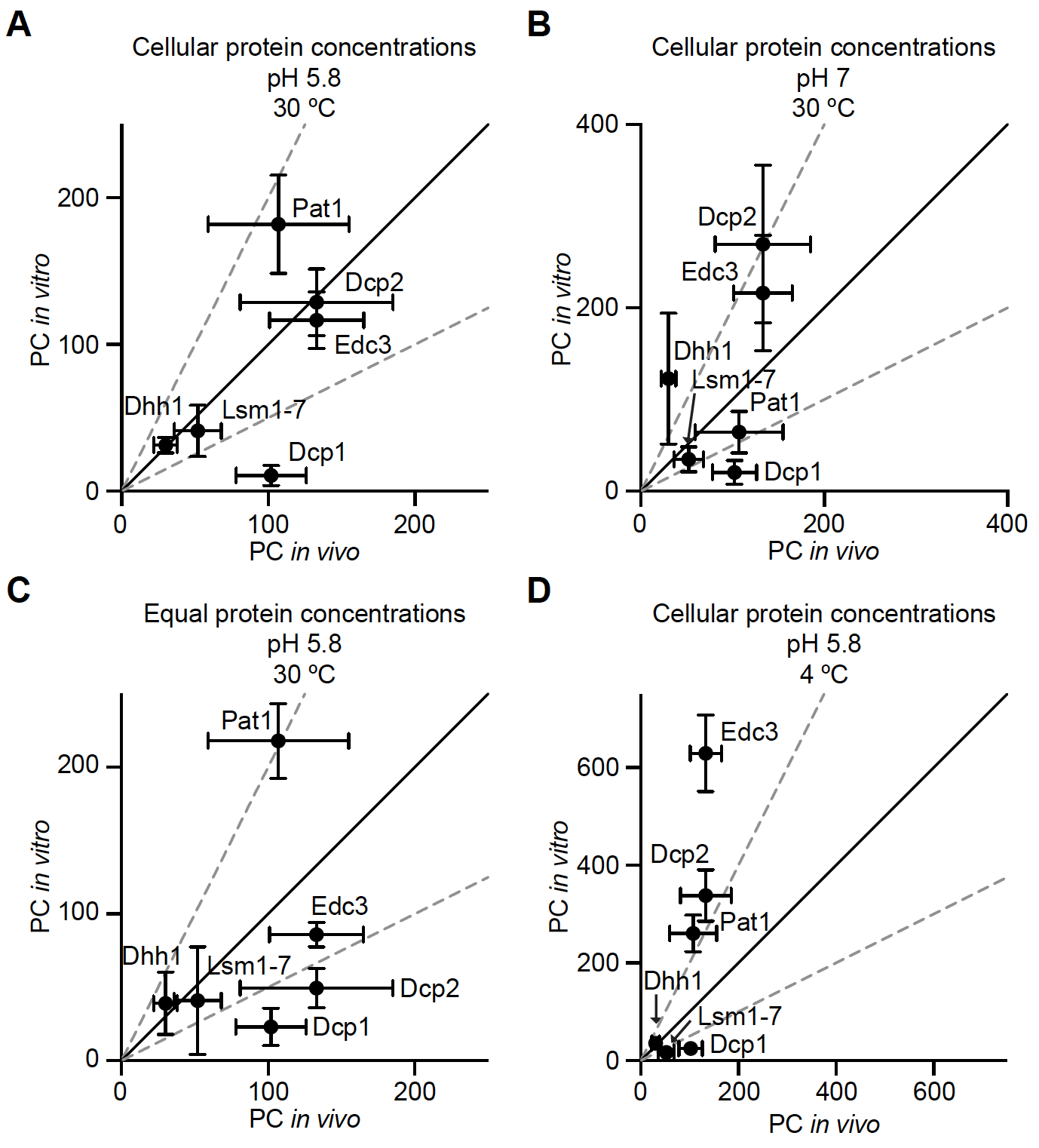

**Figure S21. Correlation between P-body protein partition coefficients *in vitro* and *in vivo*.**

We compared the partition coefficients of P-body proteins in *S. cerevisiae (4)*, to *in vitro* values observed under different experimental conditions.

(A) Cellular protein concentrations (range 90 – 370 nM, see Figure S5), pH 5.8, and 30 °C incubation temperature. Black diagonal line is where *in vivo* and *in vitro* values are equivalent, gray dotted lines indicate twofold differences. This is the same data as in Fig. 4C, included here for reference to other experimental conditions.

(B) Same as (A) but pH 7 instead of pH 5.8.

(C) Same as (A) but with 150 nM input concentrations for all proteins and RNA.

(D) Same as (A) but with 4 °C instead of 30 °C incubation.

Note the difference in axes scales in (B) and (D). Partition coefficients for all proteins are closest to *in vivo* values with cellular protein concentrations, pH 5.8, and 30 °C incubation, and provide validation for using these experimental conditions. Furthermore, these data suggest that the stoichiometry of P bodies are sensitive to parameters such as pH, protein concentrations, and temperature.

See also Fig. 5C and S5.

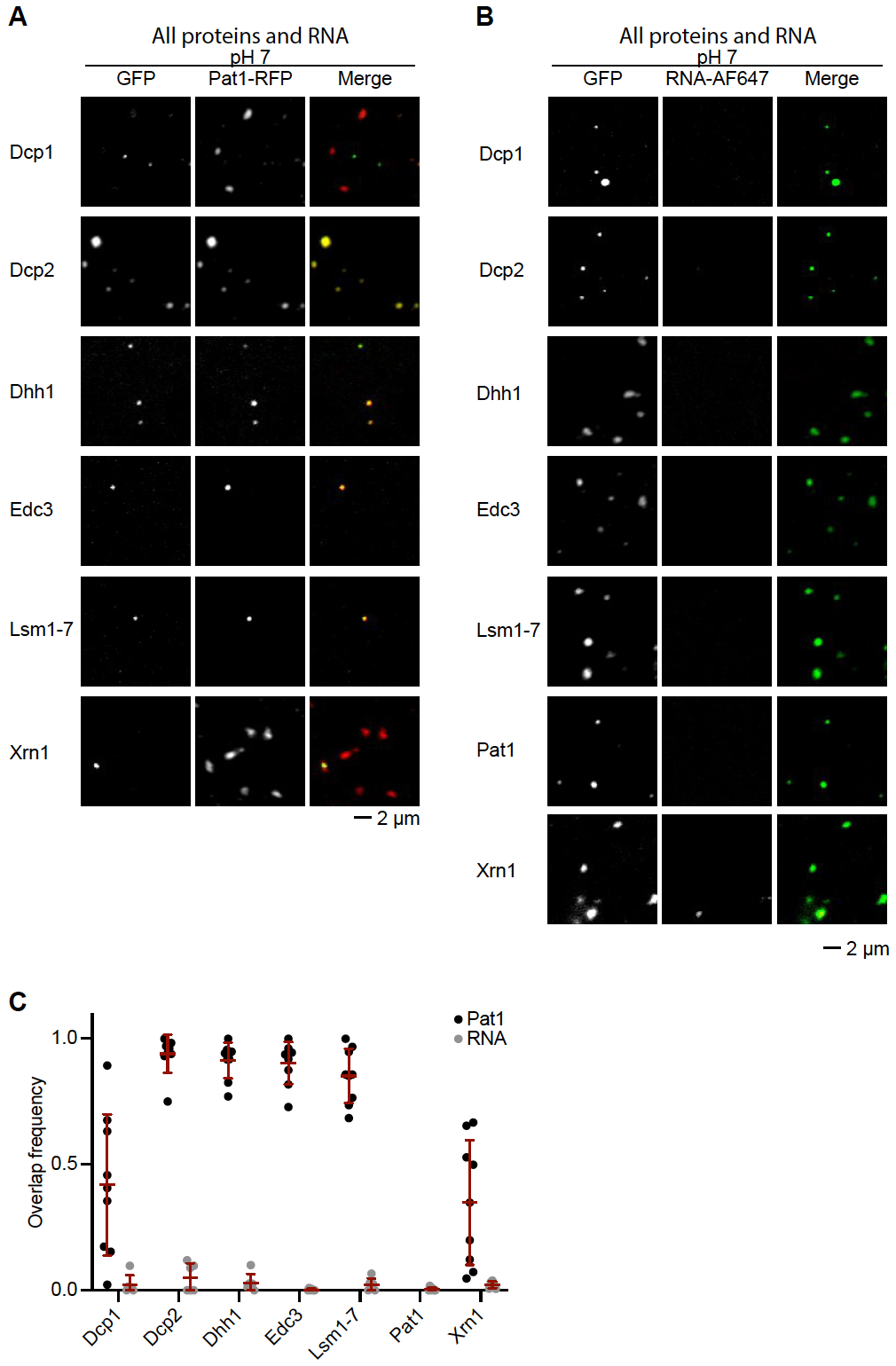

**Figure S22. Less overlap between P-body components at neutral pH (7.0).**

(A) Representative micrographs for Dcp1, Dcp2, Dhh1, Edc3, and Lsm1-7 overlapping with Pat1 under neutral conditions (pH 7.0).

(B) Representative micrographs for Dcp1, Dcp2, Dhh1, Edc3, Lsm1-7, and Pat1 overlap with RNA under neutral conditions (pH 7.0).

(C) Overlap between Pat1 (black) or RNA (gray) and P-body proteins.

Dcp1, Dcp2, Dhh1, and Edc3 are EGFP-fusion proteins and Lsm1-7 and Xrn1 are conjugated to AlexaFluor488.

Relative to acidic pH (5.8) conditions (Fig. 5A-B and S19), there is less overlap between RNA with all P-body proteins, and between Dcp1 and Xrn1 with Pat1. These results suggest that acidic pH are the more relevant conditions for P-body formation as all components overlap in that condition.

See also Fig. 5A-B and S20.

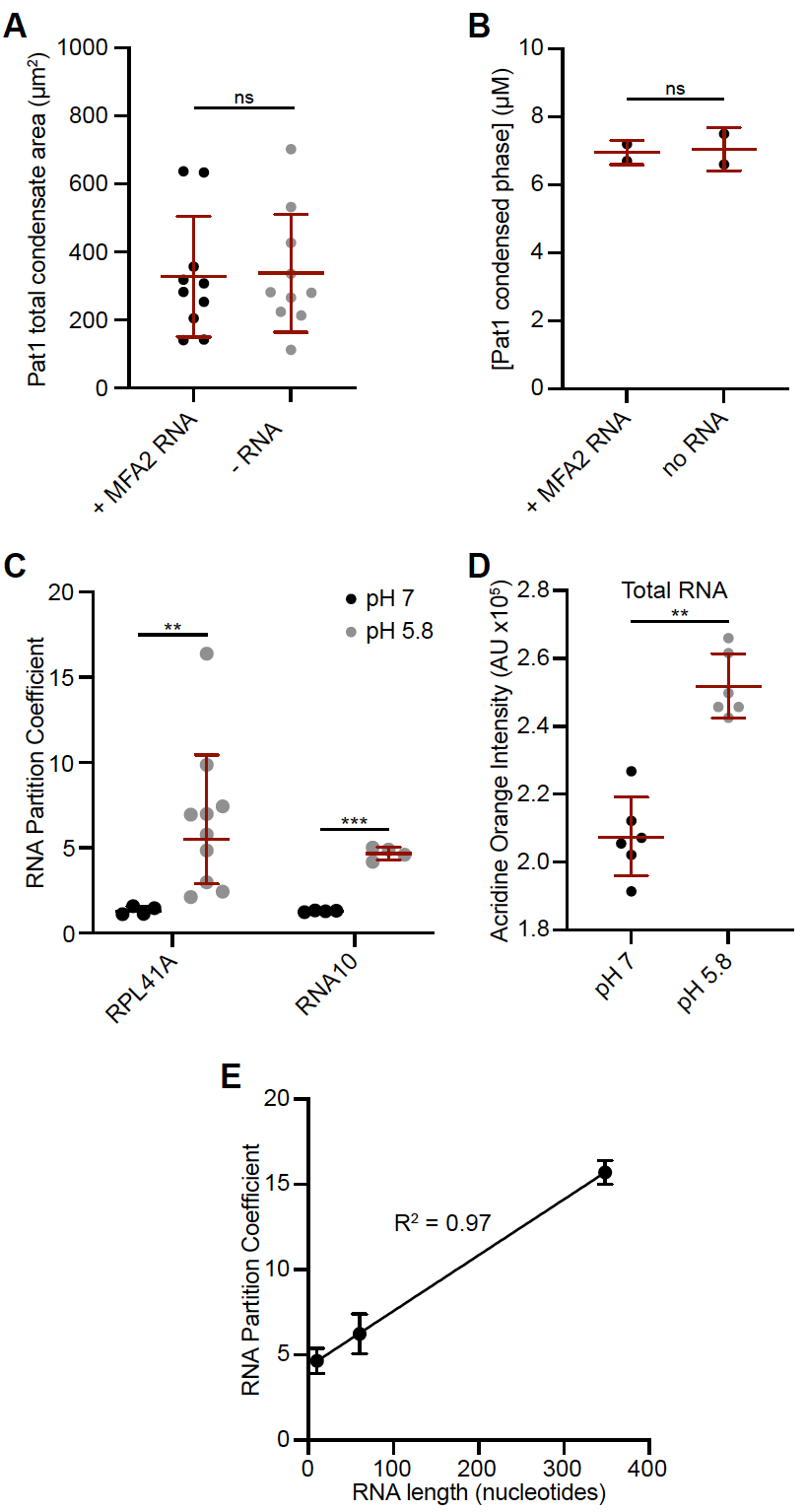

**Figure S23.** **RNA does not strongly impact *in vitro* P-body formation and is recruited in a length-dependent manner.**

(A) Condensate area, using Pat1 as a marker, is the same when all proteins are present with or without MFA2 RNA.

(B) Pat1 condensed phase concentration in condensates with all other P-body proteins is the same with or without MFA2 RNA.

(C) Partition coefficients for RPL41A and RNA10 RNAs at pH 7 (black) and pH 5.8 (gray).

(D) Acridine Orange intensity in P bodies when using total yeast RNA as the source of RNA.

(E) Correlation between RNA length and condensed phase concentration for different RNA species. Longer RNAs are more highly enriched in *in vitro* P bodies.

Four different types of RNA were tested in our reconstitution: RNA10, a 10 nucleotide single stranded RNA; RPL41A, a 60 nucleotide RNA consisting of portions of the coding and untranslated regions of ribosomal protein of the large subunit 41A; MFA2, a 348 nucleotide full-length mRNA of yeast Mating Factor Alpha that is known to localize to P bodies; and total RNA from yeast. For all RNA sources we found that RNA had little impact on condensate formation and partitioning of proteins into condensates (data for RPL41A and MFA2 shown in Fig. 5D and S23A-B). Furthermore, we observed that all RNAs were more highly enriched into condensates under acidic pH conditions (Fig. S23C-D), consistent with enhanced binding with P-body proteins under these conditions (Fig. 3). Lastly, the partitioning of RNA species correlates with their length (Fig. S23E). These data indicate that RNA does not strongly contribute to P-body formation in our reconstitution, but rather RNA is recruited into P bodies in a length-dependent manner.

See also Fig. 5D, S11, S19, and S22.

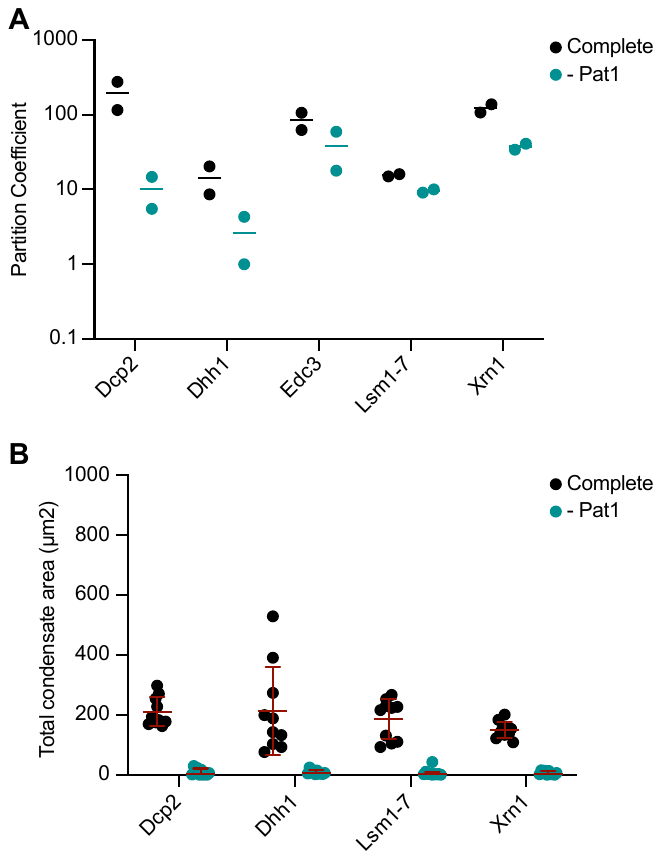

**Figure S24. Loss of Pat1 reduces the partition coefficient of other P-body proteins.**

(A) Partition coefficients with samples containing all proteins and RNA (Complete, black circles) and with all proteins, except Pat1, and RNA (- Pat1, teal circles).

(B) Total condensate area per micrograph with the same conditions as (A).

Pat1 contributes to P-body formation and increases the partitioning of the other molecules into P bodies.

See also Figure 5E.

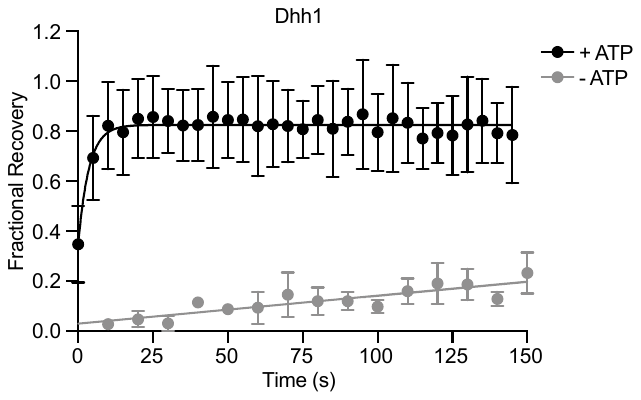

**Figure S25. ATP is required for rapid dynamics of Dhh1 *in vitro***.

Recovery of Dhh1 in FRAP experiments with (black) or without (gray) ATP included in the reaction mix. The condensates for these experiments contained all proteins and RNA. This finding suggests that ATP hydrolysis is necessary to mimic the cellular dynamics of Dhh1 *in vitro*, and is consistent with the finding that an ATP-binding deficient mutant of Dhh1 has reduced dynamics in cells (1, 4).

See also Fig. 5F.

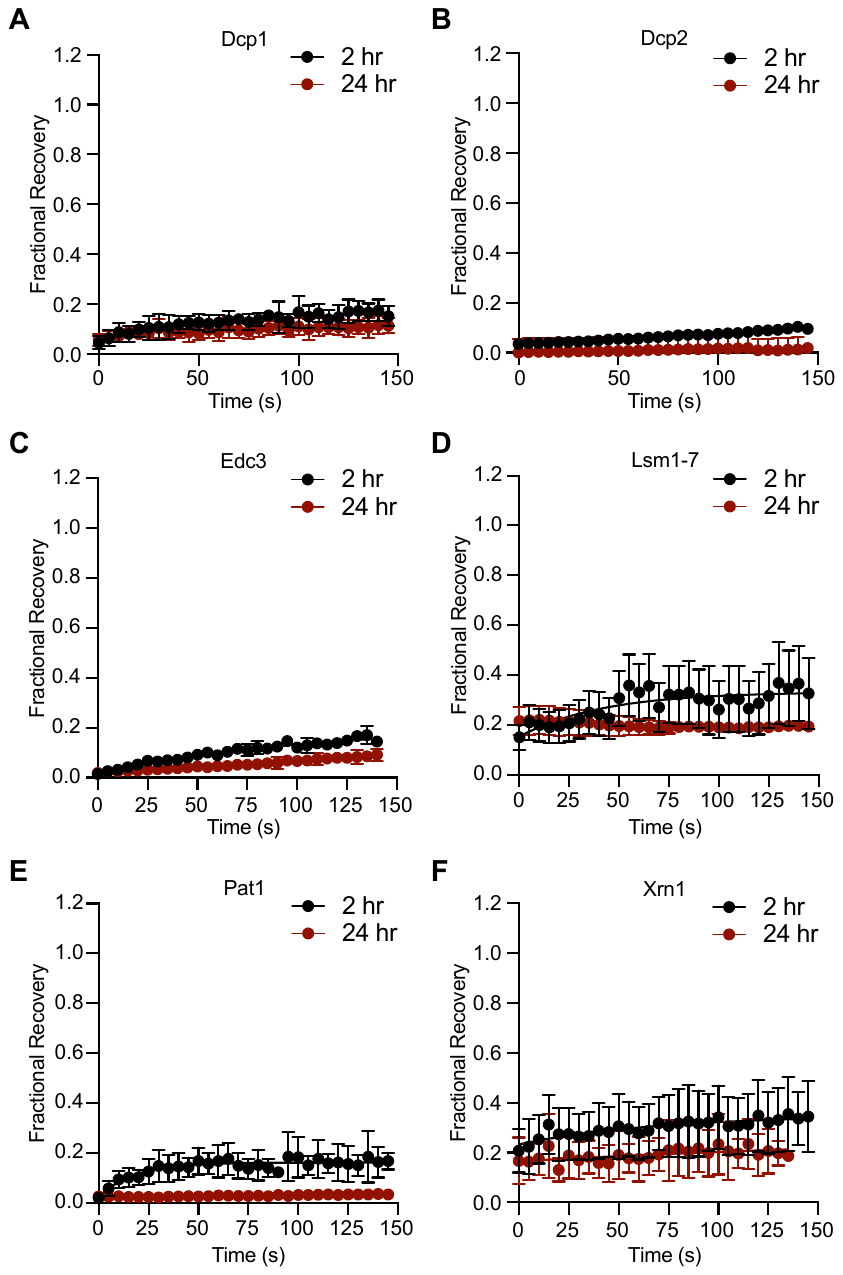

**Figure S26. Proteins become less dynamic as P-bodies mature.**

(A) Representative FRAP recovery curves for Dcp1 in heterotypic condensates with all P-body proteins and RNA at 2 hr (black) and 24 hr (red).

(B) Dcp2 FRAP recovery curves as in (A).

(C) Edc3 FRAP recovery curves as in (A).

(D) Lsm1-7 FRAP recovery curves as in (A).

(E) Pat1 FRAP recovery curves as in (A).

(F) Xrn1 FRAP recovery curves as in (A).

See also Fig. 5G and 5H.

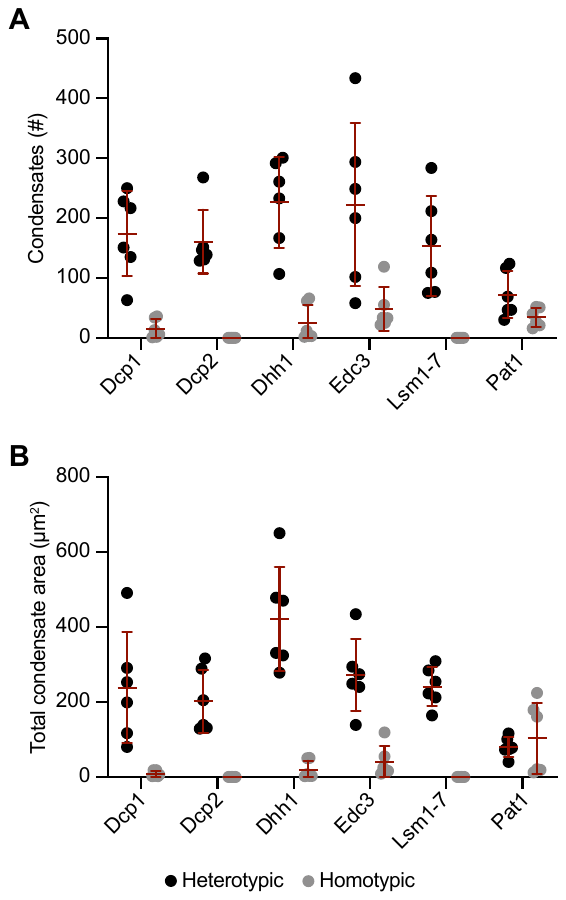

**Figure S27. Comparison between homotypic and heterotypic LLPS for P-body proteins.**

(A) Number of condensates were compared for P-body proteins either in the presence of all other P-body molecules (heterotypic, black) or as individual proteins (homotypic, gray).

(B) Total condensate area, as in (A).

The individual proteins Pat1, Edc3, and Dhh1 had the highest number of condensates and total condensate area, supporting their importance in P-body formation. The number and area of Pat1 alone is comparable to Pat1 in heterotypic condensates, suggesting the importance of Pat1 for P-body formation.

See also Fig. 6A.

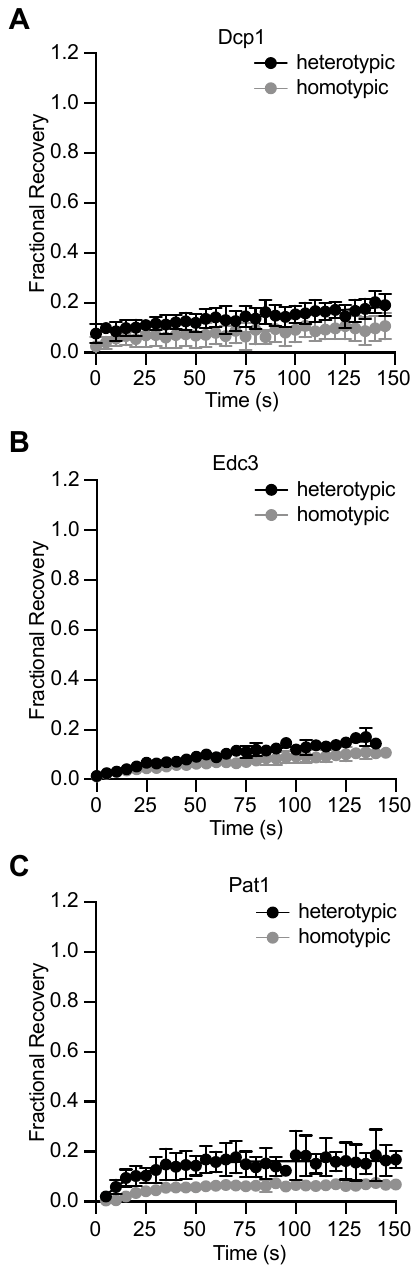

**Figure S28. Dcp1, Edc3, and Pat1 are slightly more dynamic in heterotypic condensates.**

(A) Representative FRAP recovery curves for Dcp1 in heterotypic (black) and homotypic (gray) condensates.

(B) Representative FRAP recovery curves for Edc3 as in (A).

(C) Representative FRAP recovery curves for Pat1 as in (A).

All three proteins are slightly more dynamic in heterotypic condensates as compared to homotypic condensates.

See also Fig. 6C.

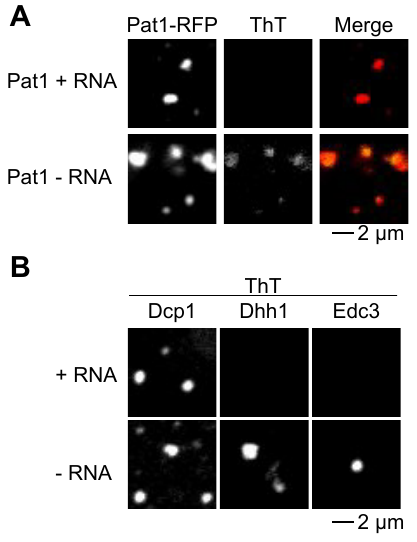

**Figure S29. RNA prevents ThT staining of single-protein condensates.**

(A) Representative micrographs monitoring Pat1-RFP (left), ThT staining (middle) and merge (left). Samples have RNA in top row and are without RNA in the bottom row.

(B) Representative micrographs of ThT staining for Dcp1 (left), Dhh1 (middle), and Edc3 (right). Fraction of ThT positive condensates for these proteins were estimated with comparison to GFP-tagged proteins measured in parallel.

See also Fig. 6D-E.

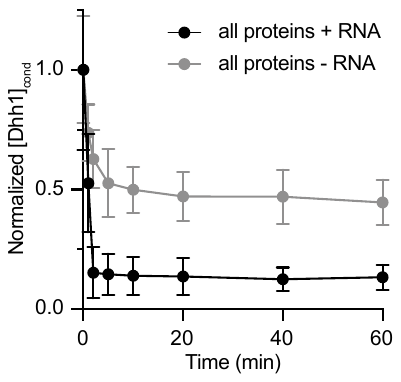

**Figure S30. RNA promotes proteolytic degradation of condensates.**

Relative Dhh1 condensate protein concentration before trypsin is added (0 min), and after trypsin is added. Black circles and line are for all proteins with RNA; gray circles and line are for all proteins without RNA. Condensates with RNA are more rapidly and completely proteolyzed by trypsin, suggesting that RNA promotes a more mobile and reversible condensate material state.

See also Fig. 6F-G.

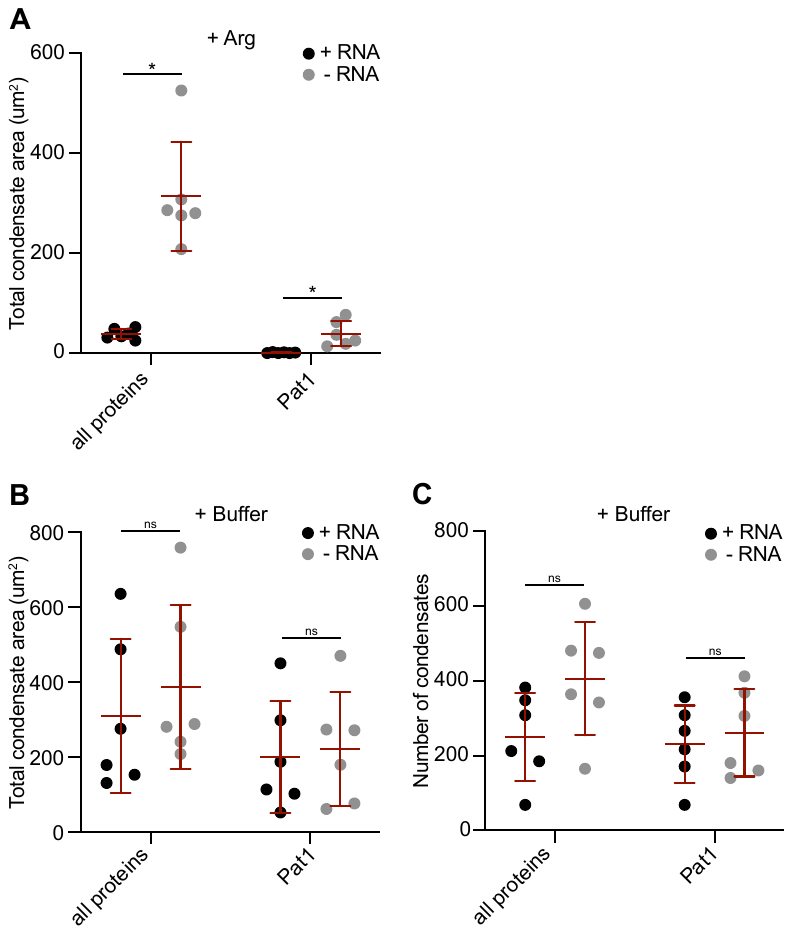

**Figure S31. Dissolution of RNA-containing condensates with arginine.**

(A) Total condensate area per micrograph for condensates with (black) and without RNA (gray) after addition of 500 mM arginine.

(B) Total condensate area per micrograph for condensates with (black) and without RNA (gray) after addition of an equivalent volume of buffer to control for dilution as compared to the arginine treatment.

(C) Number of condensates per micrograph for condensates with (black) and without RNA (gray) after addition of an equivalent volume of buffer to control for dilution as compared to the arginine treatment.

Arginine more readily dissolves condensates with RNA compared to those without RNA. The buffer control demonstrates that condensates with all proteins or with Pat1 have similar number of condensates and similar condensate area with or without RNA.

See also Fig. 6H-I.

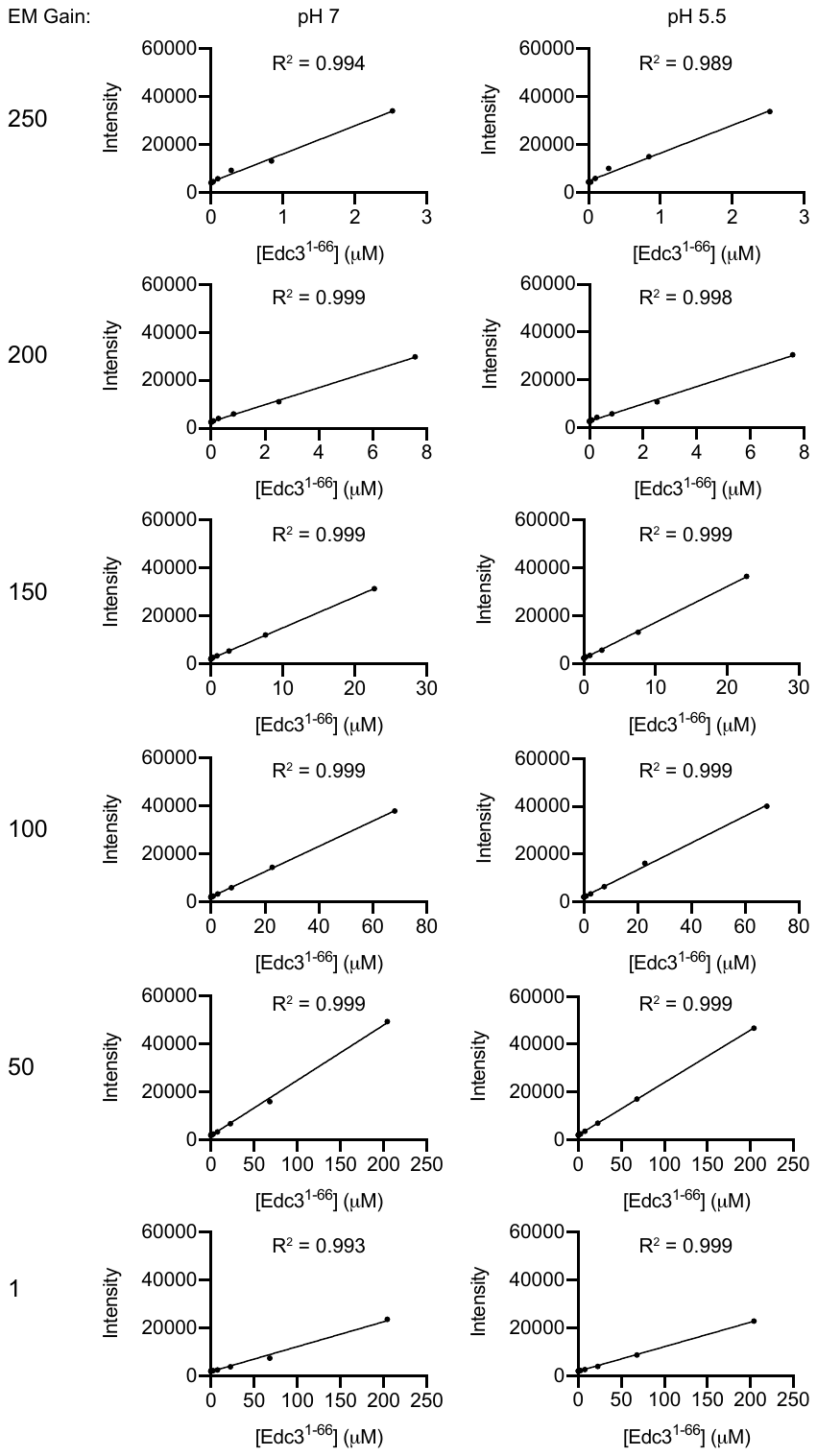

**Figure S32. EGFP standard curves for quantitative microscopy.**

Standard curves were measured using six different EM gain settings to obtain a linear relationship between protein concentration and pixel intensity over a wide range of protein concentrations. Note the change in x-axis scale between different EM gain settings. Standard curves were done in pH 7 (left) and pH 5.8 (right). MBP-Edc3^1-66^-EGFP was used to generate standard curves because it did not form condensates at any of these concentrations.

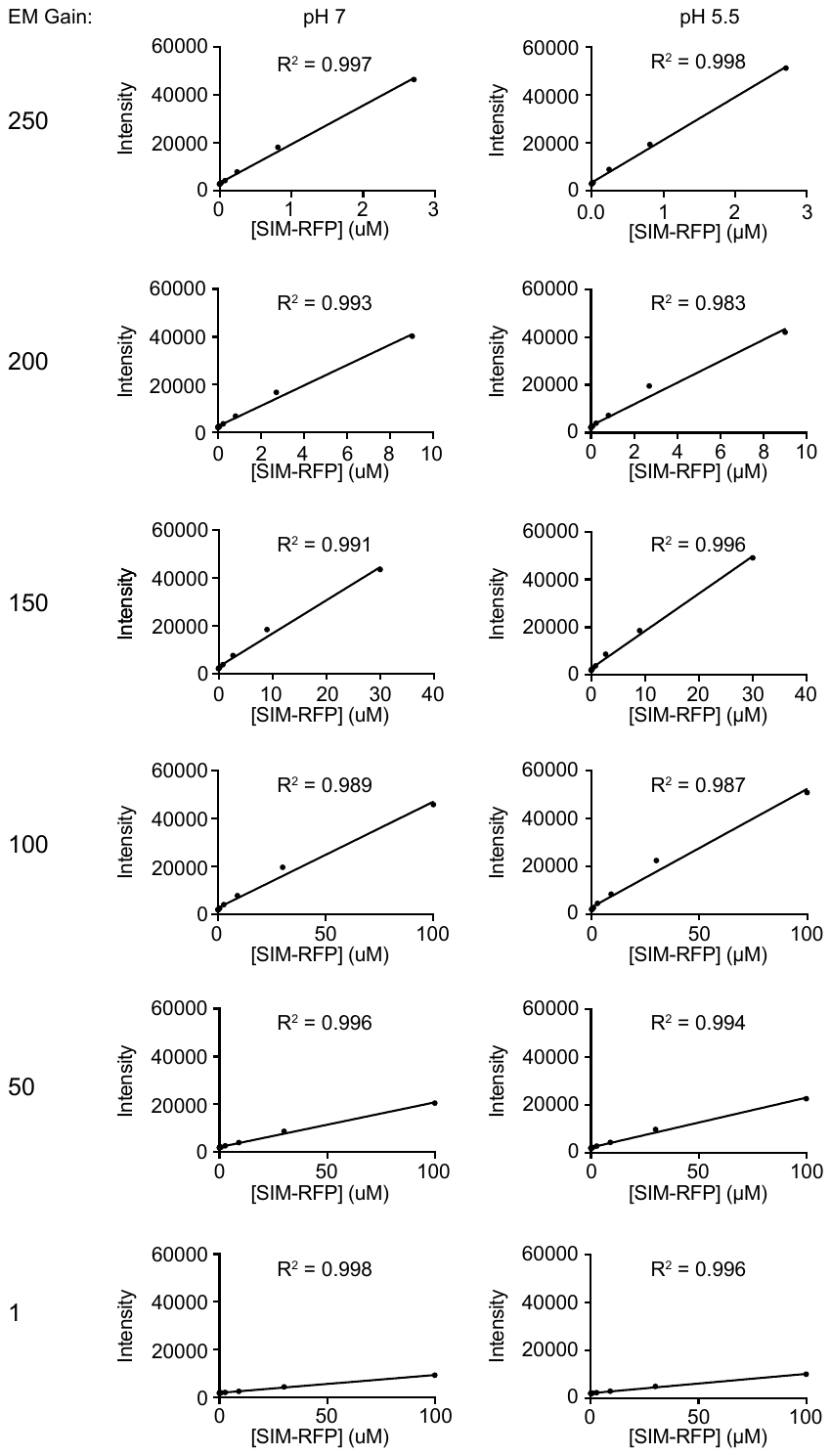

**Figure S33. RFP standard curves for quantitative microscopy.**

Standard curves were measured using six different EM gain settings to obtain a linear relationship between protein concentration and pixel intensity over a wide range of protein concentrations. Note the change in x-axis scale between different EM gain settings. Standard curves were done in pH 7 and pH 5.8. A single SUMO interacting motif (SIM) fused to RFP was used to generate standard curves because it did not form condensates at any of these concentrations.

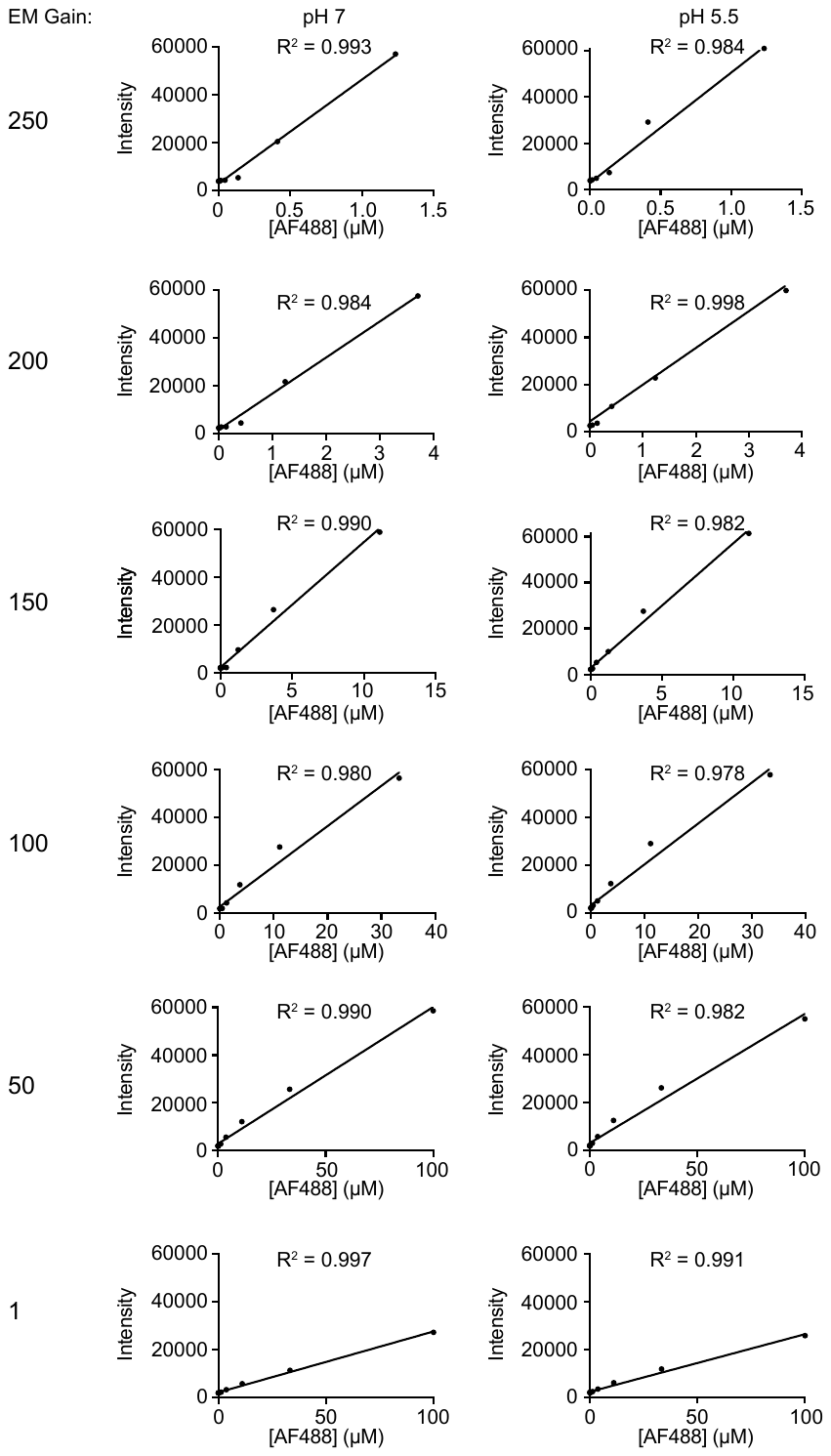

**Figure S34. AlexaFluor488 (AF488) standard curves for quantitative microscopy.**

Standard curves were measured using six different EM gain settings to obtain a linear relationship between protein concentration and pixel intensity over a wide range of protein concentrations. Note the change in x-axis scale between different EM gain settings. Standard curves were done in pH 7 and pH 5.8. Free AF488 (not bound to any protein) was used to generate standard curves.

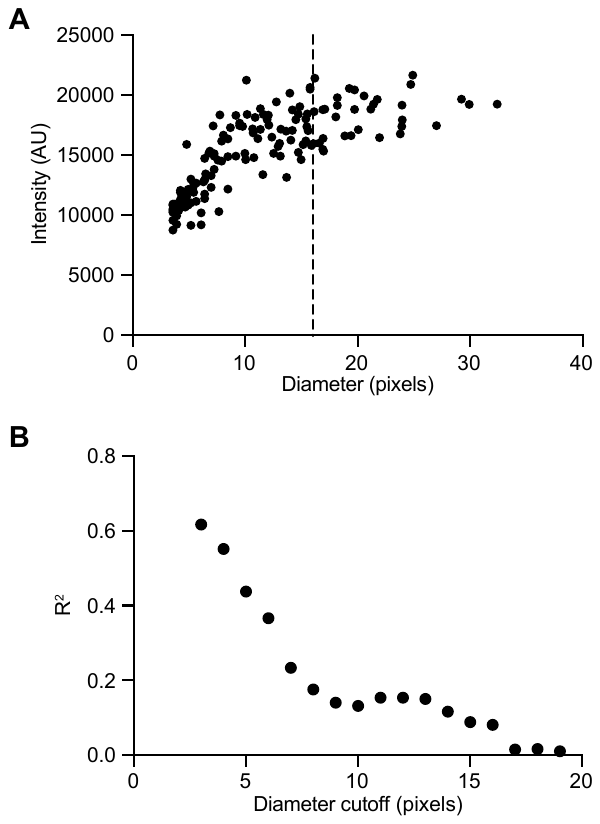

**Figure S35. Diameter cutoff for measuring condensate concentration.**

(A) Fluorescence intensity versus condensate diameter for Dhh1-GFP as a marker of *in vitro* P-bodies, i.e. with all other P body proteins and RNA.

(B) R^2^ values measuring the linear relationship for the data in (A) above the given diameter cutoff on the x-axis.

For diameters greater than 16 pixels the fluorescence intensity is unrelated to the size of the condensate. 16 pixels corresponds to approximately 2.5 μm on our microscope/detector. Therefore condensates with diameters greater than 2.5 μm were selected for concentration measurements to avoid dilution of intensities in condensates close in size to the PSF.
